# SpaCir-VDJ: a broadly compatible circularization strategy for spatial immune repertoire profiling

**DOI:** 10.64898/2026.04.26.720528

**Authors:** Zheyu Hu, Danni Guo, Xiuyuan Wang, Zhaoxuan Zhang, Zizhao Gao, Xueting Li, Maiyi Zhong, Xie Wang, Wei Wang, Zhe Wang, Xiao Liu

**Affiliations:** Institute of Biopharmaceutical and Health Engineering, Tsinghua Shenzhen International Graduate School, Tsinghua University, Shenzhen 518055, China; Department of Pathology, the First Affiliated Hospital of USTC, Division of Life Sciences and Medicine, University of Science and Technology of China, Hefei, Anhui, 230036, China; State Key Laboratory of Holistic Integrative Management of Gastrointestinal Cancer, Department of Pathology, Xijing Hospital and School of Basic Medicine, Air Force Medical University, Xi’an 710032, China; NeoImmune Co., Ltd., Shenzhen 518081, China

**Author notes:** These authors contributed equally to this work (Zheyu Hu, Danni Guo, Xiuyuan Wang, Zhaoxuan Zhang, Zizhao Gao). Correspondence: Xiao Liu; Zhe Wang.

## Abstract

Deciphering the spatial organization of immune clonal lineages is critical for understanding the evolution of adaptive immune responses. However, mainstream spatial transcriptomics platforms struggle to reconcile cost-effective short-read sequencing with the simultaneous and efficient capture of V(D)J sequences. To address this gap, we developed SpaCir-VDJ, an approach that utilizes a “multiplex PCR enrichment plus library circularization” strategy to integrate spatial barcodes and V(D)J CDR3 regions into a compact circular template optimized for standard short-read sequencing. This workflow is accompanied by an automated computational pipeline designed for circular-library data parsing, sequence reconstruction, spatial barcode deconvolution, and high-fidelity clonotype calling.

In human lymphoid tissues, SpaCir-VDJ delineated germinal center (GC)-associated clonal organization and isotype-linked somatic hypermutation (SHM) patterns. Notably, lineage-based spatial analyses inferred inter-GC clonal redistribution events, suggesting that GCs in the analyzed samples may not be fully isolated reaction units. Functional perturbation provided orthogonal support for candidate SHM regulators identified by spatial analysis, with knockdown of *MDM4*, *LYN*, *EAF2*, and *H2AFY* each leading to reduced SHM. Finally, in gastric cancer, SpaCir-VDJ localized dominant BCR lineages within tertiary lymphoid structures and tumor-margin plasma cell–rich regions, with lineage patterns consistent with extension toward the tumor interior. We further observed a significant positive correlation between cumulative tumor mutations and spatial TCR diversity, consistent with a spatial association between local mutational burden and T-cell remodeling within the tumor microenvironment. Collectively, SpaCir-VDJ provides a scalable and cost-effective framework for interrogating the spatiotemporal dynamics of immune microenvironments.

## Introduction

The adaptive immune system, composed of T cells and B cells, is central to host defense against pathogens and to immune surveillance of malignant tumors [1, 2]. This protective capacity arises from the extensive diversity of T-cell receptors (TCRs) and B-cell receptors (BCRs) generated through V(D)J recombination, enabling specific recognition of a broad range of antigens [3]. However, key steps of immune responses are not determined solely by “cell-type composition” but are jointly shaped by the histological microenvironment and its spatial organization. In secondary lymphoid organs, germinal centers (GCs) exhibit highly polarized spatial compartmentalization: B cells proliferate rapidly and undergo activation-induced cytidine deaminase (AID)–mediated somatic hypermutation (SHM) in the dark zone (DZ), then migrate to the light zone (LZ), where they undergo T follicular helper (Tfh) cell–mediated affinity selection. Cyclic trafficking between the DZ and LZ provides the kinetic basis for antibody affinity maturation [4, 5]. Ectopic tertiary lymphoid structures (TLS) that arise in tumors and chronic inflammation can likewise support GC-like reactions, promoting local antigen presentation and clonal evolution; their spatial maturation is closely associated with patient prognosis and responses to immunotherapy [6–9]. In recent years, integration of single-cell RNA sequencing (scRNA-seq) with immune repertoire sequencing (V(D)J profiling) has enabled direct linkage of clonal expansion to transcriptional phenotypes, differentiation states, and functional programs at single-cell resolution, thereby substantially advancing our understanding of clonal expansion, differentiation, and functional programs during immune responses [10, 11]. Nevertheless, the cell-dissociation step required for single-cell sequencing inevitably obliterates the spatial context between immune cells and complex tissue microenvironments, limiting the ability to resolve the spatial dynamics that underlie immune responses. Therefore, the absence of spatial information markedly limits our ability to dissect how clonotypes expand, migrate, and undergo functional differentiation within specific microenvironmental niches.

The emergence of spatial transcriptomics (ST) provides a new framework for in situ dissection of cell–cell interactions and tissue architecture and has become an important bridge between single-cell resolution and tissue-scale phenotypes [12, 13]. However, mainstream high-throughput platforms (e.g., 10x Genomics Visium and related 3′-capture schemes) still face practical bottlenecks in directly recovering TCR/BCR variable-region information [13]. Immune receptor transcripts are relatively low in abundance in complex tissues, and the V(D)J hypervariable region and spatial barcode often cannot be captured within the same read in short-read sequencing, thereby constraining high-throughput construction of spatial clonotype maps. Recently, Hudson et al. and Engblom et al. generated spatial immune maps using customized enrichment strategies, but scalability and accuracy remain limited. Hudson’s approach relies on relatively expensive MiSeq PE300 sequencing and requires inclusion of a defined fraction of PhiX to balance sequencing libraries, which restricts scalability for large studies [14]. By contrast, Spatial VDJ proposed by Engblom and colleagues combines hybrid-capture probes with long-read sequencing to couple spatial clonotypes to transcriptomes; however, the high cost of custom probes, the elevated base-error rate of long-read sequencing (which can inflate false-positive clonotype calls in highly diverse repertoires), and a comparatively complex workflow collectively raise barriers to broad adoption [15].

To overcome these limitations, we developed SpaCir-VDJ, a spatial immune repertoire sequencing technology. Building on the standard Visium library construction workflow, SpaCir-VDJ introduces a strategy of balanced multiplex PCR enrichment and library circularization. Circularization overcomes the length constraint of linear fragments by bringing distal spatial coordinate information into physical proximity with the V(D)J hypervariable region. We optimized TCR/BCR-targeted multiplex PCR primer sets and circularization efficiency to achieve efficient and balanced capture of diverse V(D)J subtypes with low bias. In parallel, we developed a matched bioinformatics pipeline for automated read processing, spatial barcode deconvolution, and downstream clonotype reconstruction and analysis.

Applied to human tonsil, lymph node, and gastric cancer tissues, SpaCir-VDJ enables joint profiling of whole-transcriptome maps and clonotype landscapes while preserving spatial context. In lymphoid tissues, SpaCir-VDJ localizes expanded clonotypes within germinal centers (GCs), delineates isotype-linked somatic hypermutation (SHM) patterns, and identifies lineage relationships consistent with inter-GC clonal redistribution, suggesting dynamic clonal exchange across GC-associated regions in the analyzed samples. Furthermore, spatial analyses prioritized candidate regulators of somatic hypermutation, and shRNA-mediated perturbation in Ramos B cells provided functional support for *MDM4*, *LYN*, *EAF2*, and *H2AFY*, with knockdown of each gene resulting in reduced SHM. In gastric cancer, SpaCir-VDJ delineates dominant BCR lineages in tertiary lymphoid structures and plasma cell–rich tumor-margin niches and reveals lineage patterns consistent with extension toward the tumor interior, while epithelial mutational burden is associated with spatial TCR remodeling. Collectively, this technology provides a methodological foundation for spatiotemporal immunogenomic studies of tumor immunity, autoimmune disease, and lymphoid malignancy.

## Results

### Development of SpaCir-VDJ for Spatial Immune Repertoire Profiling

Spatial transcriptomics remains limited in resolving immune receptor lineages, hindering precise tracking of clonal evolution in situ. To address this challenge, we developed SpaCir-VDJ, an integrated workflow for simultaneous capture of whole-transcriptome profiles and V(D)J sequences in poly(A)-capture spatial transcriptomics, implemented here using the standard 10x Genomics Visium workflow (Figure 1A). After on-slide reverse transcription, spatially barcoded cDNA was split for parallel transcriptome profiling and targeted immune receptor enrichment, enabling joint analysis of gene expression states and clonotype architecture from the same tissue section.

**Figure 1.**
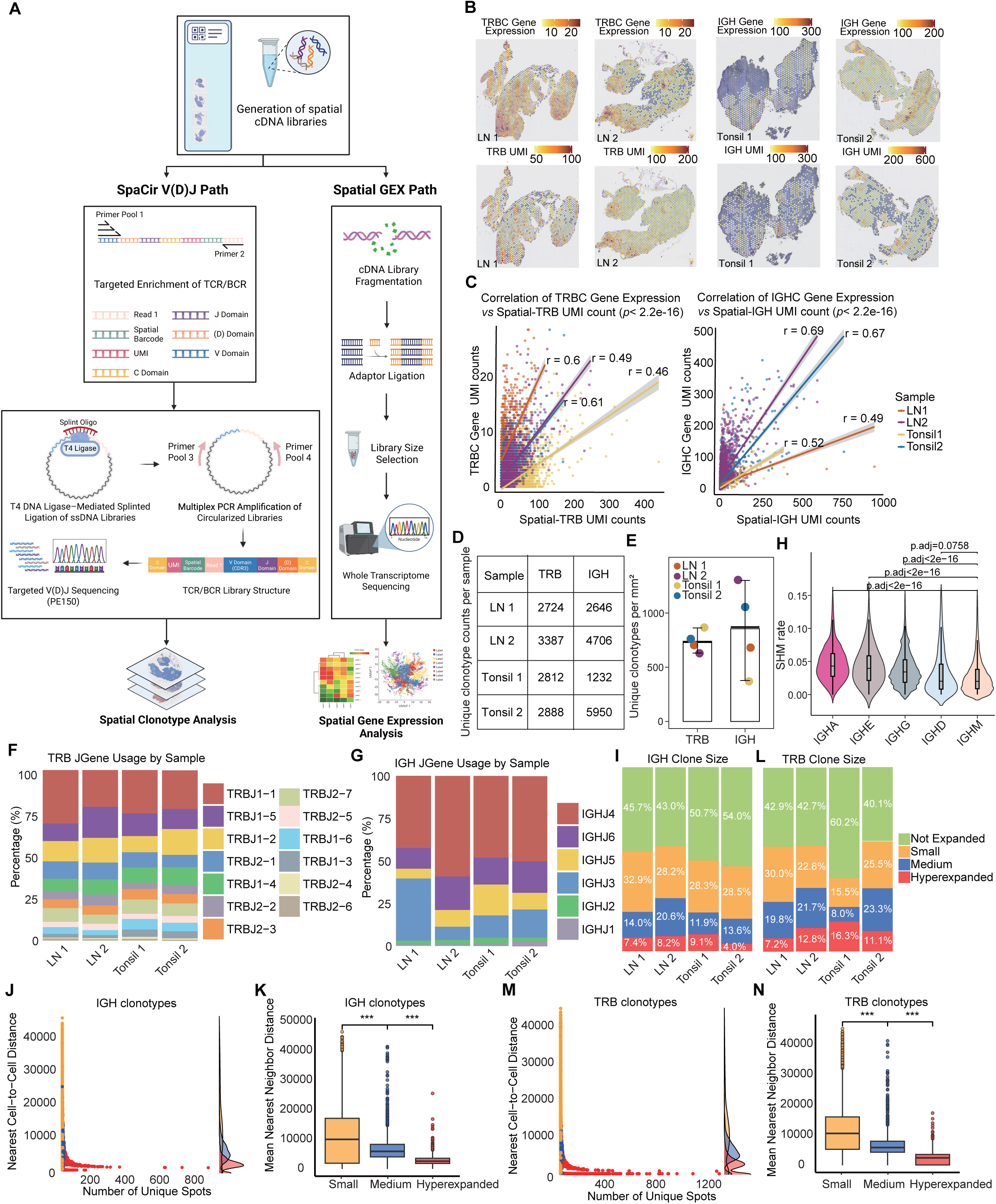
Overview and performance of SpaCir-VDJ for spatial immune repertoire profiling. (A) Schematic of the SpaCir-VDJ workflow, including targeted enrichment of TCR/BCR transcripts from Visium-derived cDNA, splint-assisted circularization, multiplex amplification, short-read sequencing, and integrated downstream spatial clonotype analysis together with whole-transcriptome profiling. (B) Representative spatial maps showing concordance between TRBC/IGHC gene expression and the corresponding spatial TRB/IGH UMI signals in human tonsil and lymph node tissues. (C) Correlation between transcript abundance and spatial repertoire capture for TRB and IGH. (D, E) Numbers and densities of unique TRB and IGH clonotypes across the four lymphoid tissue samples. (F, G) J-gene usage profiles of TRB and IGH across samples. (H) SHM rates across IGH isotypes. (I, L) Distribution of clone-size classes for IGH and TRB. (J, M) Relationships between clone spread and nearest-neighbor distance. (K, N) Hyperexpanded IGH and TRB clonotypes exhibit stronger spatial clustering than smaller clone classes.

For immune repertoire enrichment, we designed a forward multiplex primer pool (Pool 1), adapted from prior work[16, 17], targeting regions upstream of the TCR/BCR CDR3, and appended a shared fixed sequence to the 5′ end of each primer. We then introduced a reverse primer (Primer 2) complementary to the invariant Read 1 structure of the Visium cDNA, enabling targeted amplification of TCR/BCR transcripts by multiplex PCR. The enriched products were subsequently circularized using a splint oligonucleotide: the terminal 10 nucleotides of the fixed sequence and the terminal 10 nucleotides of the Read 1 end were each designed to anneal to opposite ends of a custom splint oligo, thereby “bridging” the molecule ends and enforcing their physical proximity. This configuration promotes efficient end-to-end ligation and circularization catalyzed by T4 DNA ligase (Figure 1A), generating a single-stranded circular DNA (ss-cDNA) library that brings the V-region CDR3 and the spatial barcode into a short-read–addressable arrangement on the same molecule.

Next, we performed a second round of multiplex amplification using an optimized primer set comprising Pool 3 (positioned upstream of CDR3) and Pool 4 (located in the constant region proximal to the J segment), with the circular ss-cDNA serving as the template. This step generated the final library containing the spatial barcode, CDR3, and J-segment sequences for PE150 sequencing (Figure 1A).

### Spatial Profiling of TCR and BCR Repertoires Reveals Localized Clonal Enrichment

Using the workflow described above, we performed spatial immune repertoire sequencing on two tonsil and two lymph node specimens obtained from four healthy donors (Figure 1A, B and Figure S1A, B). For each sample, IGH and TRB libraries were prepared and sequenced separately to ∼20 Gb, reaching near-saturation depth (Figure S1C). The mean expression of TRBC and IGHC in the spatial transcriptomic data was strongly and significantly correlated with the corresponding UMI counts recovered in the TRB and IGH repertoire libraries, respectively (Figure 1C), supporting concordance between transcript abundance and immune repertoire capture.

Clonotypes were defined by unique amino acid sequences of the CDR3 region. Across the four samples, the density of unique TRB and IGH clonotypes per mm² predominantly fell within the range of several hundred to over one thousand (Figure 1D, E). We next examined V and J gene usage and observed highly consistent IGH and TRB V/J usage profiles across samples (Figure 1F, G and Figure S1D, E). Finally, we assessed somatic hypermutation (SHM) across IGH isotypes. IGHD and IGHM exhibited the lowest SHM rates (Figure 1H and Figure S1F), in line with their predominant association with earlier B-cell differentiation states that have not undergone germinal-center affinity maturation.

Based on spatial UMI density and clonotype expansion levels, both IGH and TRB clones were classified into four categories: Not Expanded (clone size = 1), Small (2 ≤ clone size < 6), Medium (6 ≤ clone size < 20), and Hyperexpanded (clone size ≥ 20) (Figure 1I, J). Analysis of spatial nearest cell-to-cell distances revealed that Hyperexpanded IGH and TRB clones were significantly clustered in space (Figure 1K-N, Figure S1G), consistent with localized clonal expansion and spatial enrichment within the tissue microenvironment.

### Spatially Resolved Repertoire Diversity and Signaling Networks in Germinal Center-Associated Microenvironments

We performed histopathological annotation of tonsil and lymph node tissues by integrating H&E staining with spatial gene expression profiles [18]. Spatial cell-type deconvolution was performed using cell2location with the matched single-cell reference dataset of human tonsil and lymph node [19]. Considering tissue heterogeneity, region annotation was conducted independently for each tissue sample to ensure robustness of downstream analyses (Figure 2 and Figures S2-S4). Additionally, quality control was performed prior to downstream annotation to retain regions with reliable transcriptomic signal for subsequent analysis (Figure S4A).

**Figure 2.**
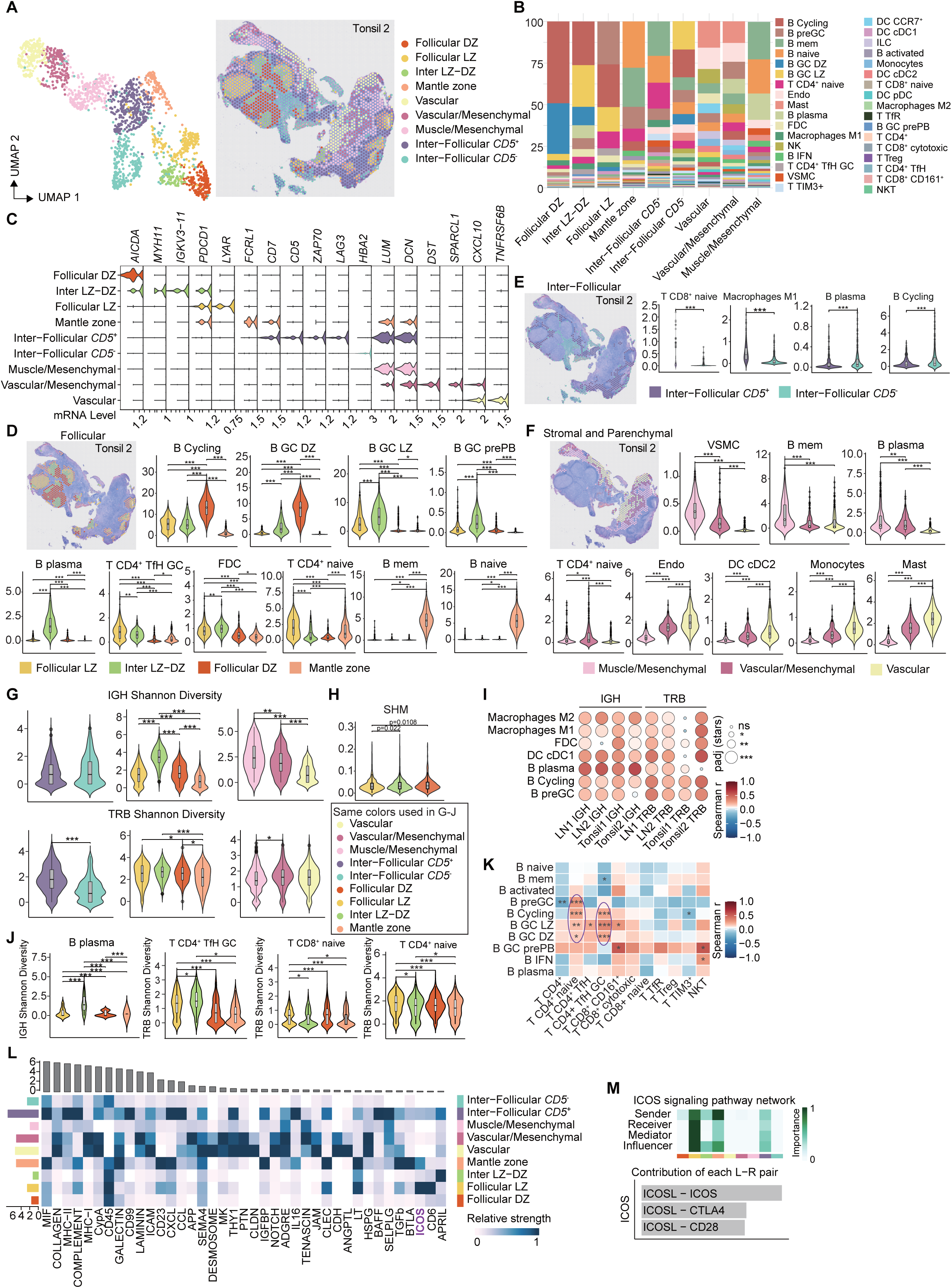
Spatial repertoire diversity and signaling architecture in the Tonsil 2 germinal-center microenvironment. (A) UMAP- and tissue-space annotation of major spatial regions, including follicular DZ, follicular LZ, inter LZ-DZ, mantle zone, vascular and mesenchymal compartments, and inter-follicular CD5+ and CD5− regions. (B) Cell-type composition across annotated regions. (C) Marker-gene expression patterns supporting regional annotation. (D–F) Regional enrichment of representative B-cell, T-cell, stromal, and myeloid populations in follicular, inter-follicular, stromal, and parenchymal compartments. (G) Region-specific IGH and TRB Shannon diversity. Colors denote tissue regions and are used consistently across panels G-J. (H) SHM enrichment in the DZ. (I) Correlations between repertoire diversity and local cell-type abundance. (J, K) Associations between repertoire diversity and selected lymphocyte subsets. (L, M) CellChat-based signaling analysis highlighting region-specific intercellular communication, including ICOS-centered interactions in the LZ.

Based on spatial gene expression and pathological annotations, we delineated major structural regions of tonsil and lymph node tissues. The dark zone (DZ) showed high AICDA expression and enrichment of B-cycling cells, whereas the LZ was enriched for LYAR, FDCs, and GC Tfh cells. In Tonsil 2, we further identified an intermediate region at the LZ-DZ interface, marked by co-expression of *MYH11* and *PDCD1*, and enriched for GC-derived pre-plasmablasts (B GC prePB) and plasma cells (B Plasma), suggesting a transitional niche for plasmablast differentiation (Figure 2A-D, Figure S2A, and Figure S3A-D). Surrounding the follicles, the mantle zone (MZ) was defined by high *CD7* and *FCRL1* expression and contained predominantly naive and memory B cells (Figure 2A-D, Figure S2A, Figure S3A-D, and Figure S4B-D). Additionally, inter-follicular (IF) regions showed substantial heterogeneity within samples, with distinct subregions enriched for different lymphoid and myeloid populations (Figure 2E, Figure S3E, and Figure S4C).

We also identified spatially organized stromal and parenchymal regions with distinct molecular and cellular profiles across samples (Figure 2A, C, F; Figure S3C, E; and Figure S4C, D). In Tonsil 2, three concentric stromal layers surrounded the GC: an inner muscle-mesenchymal zone marked by LUM/DCN and enriched in memory B and plasma cells, a vascular-mesenchymal zone with SPARCL1 and naive T cells, and an outer vascular zone characterized by CXCL10 with abundant dendritic and NK cells (Figure 2A, C, F). These observations suggest that spatially organized stromal compartments are associated with distinct immune cell compositions and may support specialized microenvironmental functions.

Furthermore, compared with the mantle zone, both IGH and TRB repertoires exhibited greater diversity in the LZ and DZ. Notably, plasma cell-enriched regions, such as the inter LZ-DZ interface and SDC1⁺ or XBP1⁺ inter-follicular zones, displayed markedly elevated IGH diversity. In contrast, TRB diversity was higher in T cell-rich regions, including the LZ and inter-follicular zones characterized by CD5⁺ or CXCL10⁺ expression (Figure 2G, Figure S3G-H, and Figure S4E). Consistent with their proliferative and affinity maturation activities, SHM levels were elevated in the DZ (Figure 2H and Figure S3I).

Correlation analysis between repertoire diversity and cell type abundance revealed that, across most tissue regions, IGH diversity was consistently positively correlated with plasma cell abundance, and in some regions also with M1/M2 macrophages (Figure 2I, Figure S2A, Figure S3I, and Figure S4F). This suggests that enhanced GC output (plasma cells) and their supportive macrophage microenvironment are associated with increased local antibody repertoire diversity. In parallel, cDC1 abundance was significantly correlated with both IGH and TRB diversity in most regions, while TRB diversity further showed positive associations with FDCs, pre-GC, and cycling B cells (Figure 2I, Figure S2A-B, Figure S3I, and Figure S4F). These findings indicate that the activity of antigen-presenting cells (cDC1, FDC), together with the accompanying T-B interactions, may indirectly promote diversification of the TCR repertoire and regulate the fate of early and proliferating B cells.

Mapping IGH/TRB repertoires to specific cell subsets revealed that CD4⁺ Tfh diversity was increased in the LZ, whereas T CD8⁺ naive diversity was higher in the DZ (Figure 2J, Figure S3J, and Figure S4G). Shannon diversity correlation further showed that T CD4⁺ naive and T CD4⁺ Tfh diversity were significantly associated with GC B cells (Figure 2K, Figure S3K, and Figure S4H). Spatially, T CD4⁺ Tfh cells co-localized with LZ B cells (Figure S2G). In contrast, T CD4⁺ naive cells were primarily co-localized with pre-GC B cells (Figure S2G). These results suggest that within the same GC, a broader TCR-epitope repertoire may provide helper signals to multiple BCR epitopes in the LZ. Moreover, prior to full GC establishment, abundant naive T cells may already encounter and become “primed” by early activated B cells in the pre-GC region, potentially facilitating subsequent diverse Tfh-B interactions and broader affinity maturation.

To further dissect the molecular basis of these associations, we applied CellChat analysis, which revealed region-specific signaling networks across lymphoid compartments (Figure 2L and Figure S2C, D). In the LZ, enrichment of the ICOS pathway highlighted its central role in Tfh-mediated B cell fate decisions, consistent with the diversity-associated findings (Figure 2L, M). The inter LZ-DZ interface exhibited strong APRIL signaling, supporting pre-plasmablast enrichment (Figure 2L). In addition, the CD5⁺ inter-follicular region adjacent to the GC was associated with BAFF-mediated survival signaling, CXCL13/CXCL12 signaling, and MHC-II–related antigen-presentation programs, underscoring its bridging role in B cell activation, migration into the GC, and subsequent differentiation (Figure 2L, M, and Figure S2E, F).

### Clonal Trajectories Reveal Spatially Distributed Clonal Redistribution Across Germinal Centers

Pseudotime analysis revealed a continuous transcriptomic transition from DZ to LZ and then to the Inter LZ-DZ region within germinal centers (GCs) (Figure 3A-C, Figure S5A). The DZ region exhibited high expression of proliferation- and SHM-related genes such as *AICDA*, *DUT*, and *HMGN2*. In contrast, the LZ region expressed *BATF*, *CXCL13*, and *FDCSP*, reflecting its role in antigen presentation and affinity selection. The Inter LZ-DZ region showed elevated expression of *XBP1*, *MZB1*, and several IGH class-switching genes, suggesting its function as a transitional niche prior to plasma cell differentiation (Figure 3A-C and Figure S5A).

**Figure 3.**
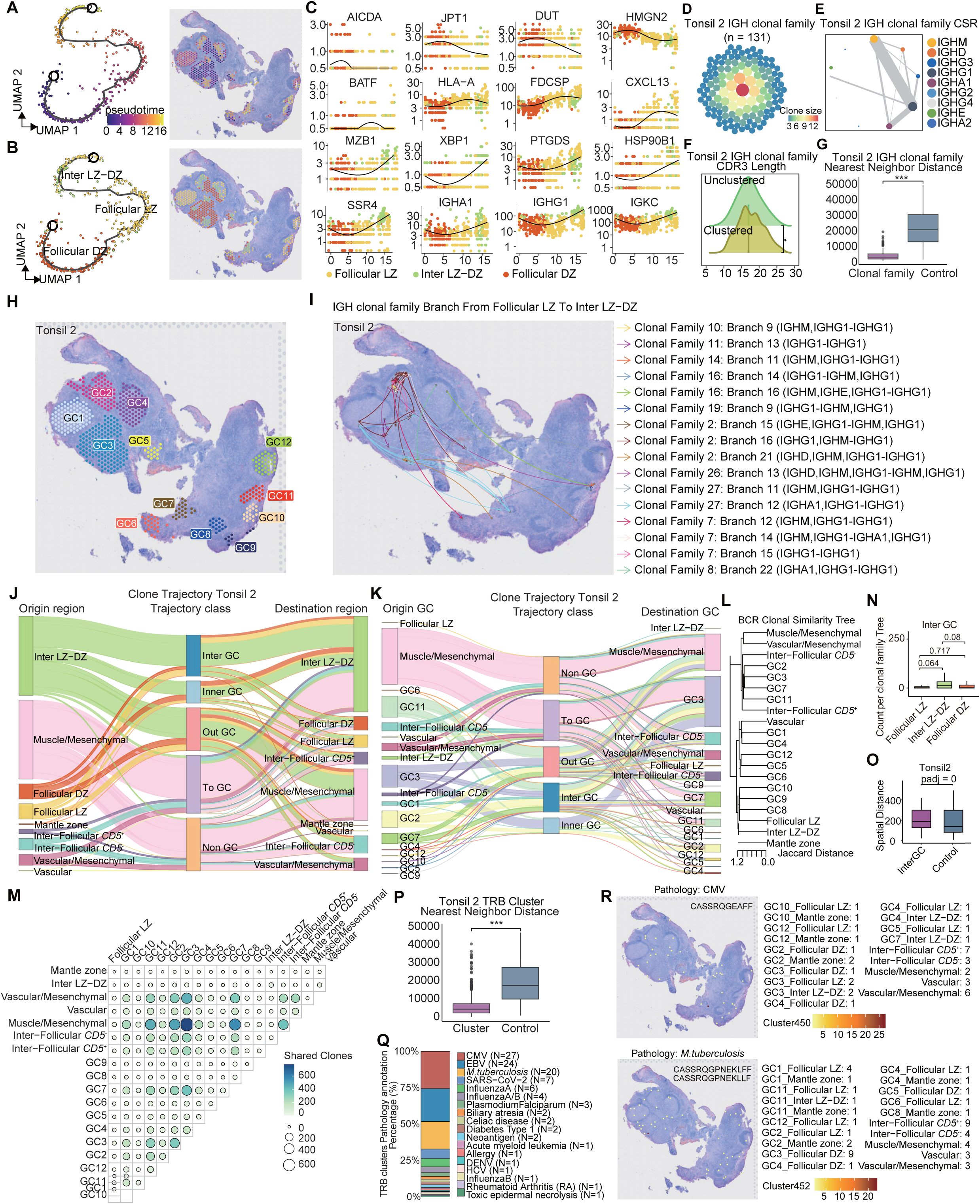
Clonal trajectories reveal spatially distributed clonal redistribution across germinal centers. (A, B) UMAP pseudotime and state annotation across follicular DZ, follicular LZ, and inter LZ-DZ compartments, together with their spatial projection onto tissue sections. (C) Dynamic expression of representative genes across the inferred DZ-to-LZ-to-inter LZ-DZ trajectory. (D–G) Example IGH clonal-family analyses showing spatial clustering, class-switch composition, CDR3-length structure, and reduced nearest-neighbor distances relative to controls. (H) Spatial map of annotated germinal centers in Tonsil 2. (I) Example lineage transitions from follicular LZ toward inter LZ-DZ. (J–K) Sankey plots summarizing inferred clone-level redistribution trajectories in Tonsil 2. In each plot, the left column indicates the origin region or GC, the middle column indicates the inferred trajectory class, and the right column indicates the destination region or GC. Flow width is proportional to the number of inferred lineage-linked transitions. (L–M) Analyses of region-pair clonal similarity and shared clonotype relationships associated with inferred inter-GC redistribution. (N) Quantification of inferred inter-GC redistribution events across GC-associated regions, highlighting preferential routing through inter LZ–DZ-associated states. (O) Spatial distances of inferred inter-GC events relative to controls. (P) Spatial clustering of GLIPH2-defined TRB clusters. (Q, R) Database-guided annotation and spatial visualization of representative TRB clusters with shared or related antigen-associated signatures.

To investigate clonal behavior, we reconstructed IGH clonal families and found that clonotypes from the same family were more spatially clustered than expected under random distribution (Figure 3D-G, Figure S6A-D, and Figure S7A-D). Based on histopathological and transcriptional features, we further delineated GC subregions (Figure 3H, Figure S6E, and Figure S7E). By constructing clonal lineage trees and projecting lineage relationships back onto tissue space, we inferred spatially distributed lineage transitions of IGH clonotypes (Figure S5B-D). These analyses suggested that many IGH families were redistributed from the LZ toward the Inter LZ-DZ region (Figure 3I), consistent with the possibility that this region represents a transitional niche following affinity selection and preceding terminal differentiation. Together, these findings support a spatially coordinated model of B-cell state transition within the GC.

We additionally identified putative inter-GC redistribution events of IGH clonotypes across distinct GCs (Figure 3H-I). To quantitatively characterize this pattern, we summarized the inferred origin and destination of lineage-linked spatial transitions and visualized the corresponding flux using Sankey plots (Figure 3J, Figure S6F, and Figure S7F). Region pairs with higher inferred transition frequency also showed increased numbers of shared clonotypes and greater clonal similarity (Figure 3K-M, Figure S6G-I, and Figure S7G-I). Moreover, these inferred inter-GC redistribution events were detected during the LZ-to-Inter LZ-DZ transition and appeared more frequently between Inter LZ-DZ regions of different GCs (Figure 3J, N). These data suggest that the Inter LZ-DZ region may represent a preferential, although not exclusive, route for cross-GC clonal redistribution. In particular, LZ-associated clones appeared more likely than DZ-associated clones to participate in these inferred cross-GC events (Figure 3J, N, Figure S6F, J). Overall, our data are consistent with the possibility that positively selected B-cell lineages can be redistributed between GC-associated regions in some samples, rather than remaining strictly confined to individual follicles. In addition, compared with randomly sampled GC-to-GC distances, some inferred inter-GC events spanned relatively longer ranges (Figure 3O, Figure S6K, Figure S7J), suggesting that such redistribution is not necessarily restricted to the nearest neighboring GCs.

Furthermore, we observed instances in which class-switched B cells appeared to re-enter the GC from other tissue regions, consistent with previous reports [20] (Figure S5F, Figure S6L, and Figure S7K). These events predominantly occurred in the IGHG1 isotype (Figure S5G, Figure S6M, and Figure S7L). Despite sample heterogeneity, the initial spots associated with GC re-entry showed higher expression of CSR-related genes than other spots, particularly *APEX1*, *UNG*, and *PRDM1* (Figure S5H, Figure S6N, and Figure S7M).

In addition to B-cell clonal dynamics, we further explored the spatial characteristics of T cells. Using GLIPH2 clustering based on shared V and J genes and CDR3 amino acid sequences, we identified TRB clusters that also exhibited strong spatial aggregation (Figure 3P, Figure S6O, and Figure S7N). Annotation of TRB sequences using the VDJdb and McPAS-TCR databases revealed that multiple GC regions contained TCR clonotypes with shared or related database-annotated specificity patterns (Figure 3Q, R, Figure S5I, Figure S6P, Q, and Figure S7O, P). These results suggest that, in addition to B-cell clonal redistribution, T cells may also participate in spatially shared antigen-associated programs across distinct GC regions, although direct antigen recognition cannot be established from sequence annotation alone.

Taken together, our findings support dynamic B- and T-cell clonal exchange across GC-associated regions and suggest that, in the analyzed tissues, GCs may participate in a broader tissue-level clonal communication network rather than behaving as completely isolated immune units.

### Stage-Associated BCR and TCR Clonal Remodeling During Germinal Center Maturation

To systematically categorize clonal behaviors across GCs, we grouped IGH clonotypes into four categories based on their spatial distribution across GCs and their expansion status (Figure 4A-B, Figure S9A-B, and Figure S10A-B). Given our earlier observation that positively selected clones may be redistributed across GCs (Figure 3J-K, Figure S6F), we speculated that shared and expanded clonotypes may constitute functional surveillance populations enabling inter-GC immunosurveillance. Interestingly, unexpanded clonotypes displayed significantly higher SHM frequencies than expanded clonotypes (Figure 4C, Figure S9C, Figure S10C). This pattern is consistent with the possibility that some highly mutated yet unexpanded clones may carry non-productive or non-selected variants.

**Figure 4.**
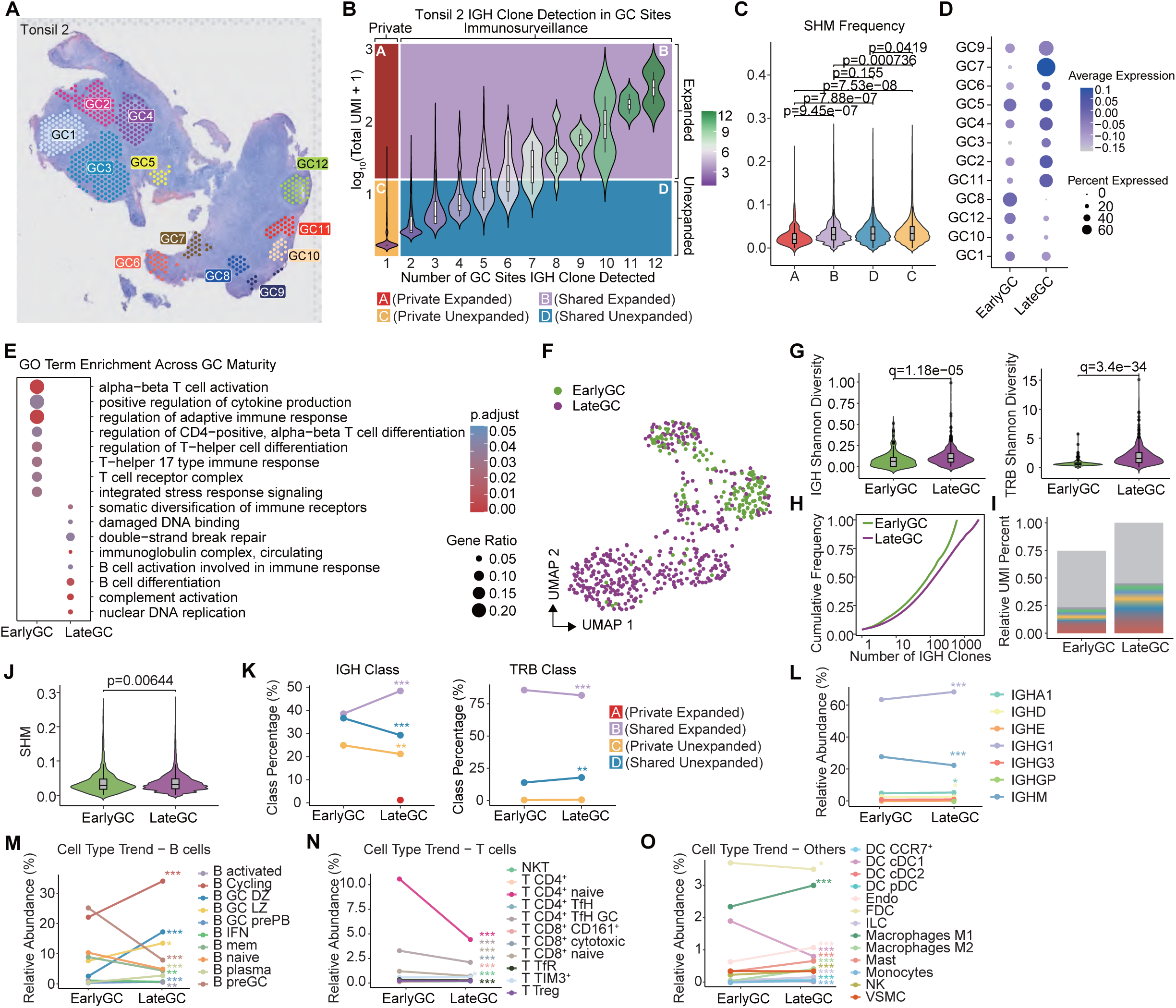
Stage-associated BCR and TCR remodeling during germinal-center maturation. (A) Spatial map of individual germinal centers in Tonsil 2. (B) IGH clonotypes were classified into four groups according to their GC distribution and expansion status: A, private expanded; B, shared expanded; C, private unexpanded; and D, shared unexpanded. Clonotypes detected in only one GC were defined as private, whereas those detected in two or more GCs were defined as shared. Expansion was determined using the 95th percentile (q95) cutoff of clone abundance. (C) SHM frequencies across clonotype classes. (D) Spatial distribution of early-GC and late-GC scores across GCs. (E) GO-term enrichment distinguishing early and late GC states. (F) UMAP separation of EarlyGC and LateGC spatial units. (G) IGH and TRB Shannon diversity in early and late GCs. (H, I) Cumulative-frequency and UMI-distribution summaries of IGH clone abundance across GC maturity states. (J) SHM comparison between early and late GCs. (K) Class composition of IGH and TRB clonotypes across GC maturity states. (L) IGH constant-region composition across GC maturity states. (M–O) Shifts in B-cell, T-cell, and other cell-type abundances between EarlyGC and LateGC regions.

We next examined the distribution of IGH clonotype classes (A-D) and constant-region usage across different GCs and found substantial heterogeneity among individual GCs (Figure S8A-B, Figure S9D-E, and Figure S10D-E). To further stratify GC states, we applied stage-associated transcriptional signatures to classify GC-associated spatial units into early-like and late-like groups. Using these signature scores, we separated GC-associated regions into EarlyGC and LateGC categories for downstream analysis (Figure 4D, Figure S9F, Figure S10F) [21–25].

Differential expression analysis between the two groups, excluding the marker genes used for classification, identified functionally distinct transcriptional signatures. As expected, EarlyGC-associated regions were enriched for pathways related to adaptive immune regulation and T-B cell interactions, including CD4⁺ T-cell help, Tfh differentiation, and Th17 regulation, consistent with early B-cell activation, interaction with T cells, and affinity selection (Figure 4E, Figure S9G). In contrast, LateGC-associated regions were enriched for pathways associated with antibody production, plasma-cell differentiation, proliferation, complement activation, DNA repair, and metabolic stress responses, consistent with later-stage differentiation following affinity selection (Figure 4E, Figure S9G). UMAP visualization further confirmed clear separation between the two groups (Figure 4F, Figure S9H). Furthermore, HdWGCNA analysis showed that hub genes identified in each group were associated with pathways consistent with their assigned GC stages, further supporting the robustness of the classification strategy (Figure S8D-E, Figure S9J-K).

We next investigated the clonal characteristics of early and late GCs. We found that both IGH and TRB clonotype diversity were increased in late GCs (Figure 4G-H, Figure S9L-M, and Figure S10G-H). This increase in clonotype diversity suggests that late GCs may retain a broader range of dominant and lower-abundance lineages than early GCs in the analyzed samples. Tracking clonotypes across stages further showed that many of the top 100 abundant IGH clonotypes in late GCs were already expanded in early GCs (Figure 4I, Figure S9N, and Figure S10I), indicating persistence of a subset of clonotypes across GC stages. By contrast, SHM frequencies did not display a consistent pattern across samples (Figure 4J, Figure S9O, and Figure S10J).

In addition, we observed that shared expanded IGH clonotypes and the IGHG1 isotype were more prevalent in late GCs, consistent with increased cross-GC clonal overlap in late GCs in these samples and more frequent class switching toward IgG1 in these samples. However, the abundance of shared expanded TRB clonotypes did not show consistent differences across GC stages (Figure 4K-L, Figure S9P-Q, and Figure S10K-L). Spatial deconvolution further suggested that preGC-associated B-cell programs were reduced, whereas LZ-, DZ-, plasma-, and cycling B-cell programs were relatively increased in late GCs, indicating a shift toward proliferation and differentiation states at the spatial-program level. By contrast, other cell populations did not show a consistent trend (Figure 4M-O, Figure S9R-T, and Figure S10M-O). Together, our results identify maturation-associated differences in clonal architecture and cellular composition between early and late GCs.

### Multi-layered Intrinsic Programs Associated with SHM in GC Dark Zones

We stratified DZ regions into three categories according to SHM ratio: High SHM (ratio > 0.05), Low SHM, and No SHM (Figure 5A-B, Figure S12A-B). AUC-weighted analysis identified distinct gene expression signatures associated with SHM intensity (Figure 5C-D, Figure S12C-D). High SHM regions were enriched for multiple regulatory factors acting at different levels of the process, including *AICDA*, which encodes the AID enzyme and represents the catalytic driver of cytidine deamination [26], chromatin remodelers and histone variants such as *CBX5*, *H2AFY*, and *HMGN3* that increase DNA accessibility and thereby facilitate AID recognition, and transcriptional/splicing-associated factors such as *POLR2F* and *DHX38* that promote transcriptional pausing and R-loop formation to guide AID targeting to hotspot loci [27, 28]. In addition, cell-cycle regulators including *CDC6*, *EAF2*, and *STAG1* were upregulated, extending dwell time in the G1/S phase and prolonging the mutational window [29], while DNA repair and nuclear structural factors such as *MDM4* and *NIBAN1* contributed to maintaining genomic stability and balancing mutation intensity to prevent excessive mutagenesis and cell loss [30]. By contrast, Low SHM regions were enriched for genes including *POU2F2*, a transcription factor implicated in B cell activation and potential AID induction, but which alone is insufficient to sustain efficient SHM in the absence of concurrent transcriptional pausing and chromatin accessibility. Other Low SHM-associated genes, such as *SNRPB*, enhanced splicing efficiency and reduced R-loop formation, thereby limiting AID targeting opportunities, while DNA repair factors such as *CHEK2* and *RNASEH2A* promoted error correction, and cell-cycle regulators such as *CDK4*, *PLK1*, and *MTHFD1* shortened G1/S transit, collectively restricting SHM efficiency [27–30]. Consistently, High SHM gene signatures showed strong positive correlation with SHM intensity, whereas Low SHM signatures did not exhibit significant correlation (Figure 5E-F, Figure S12E-F).

**Figure 5.**
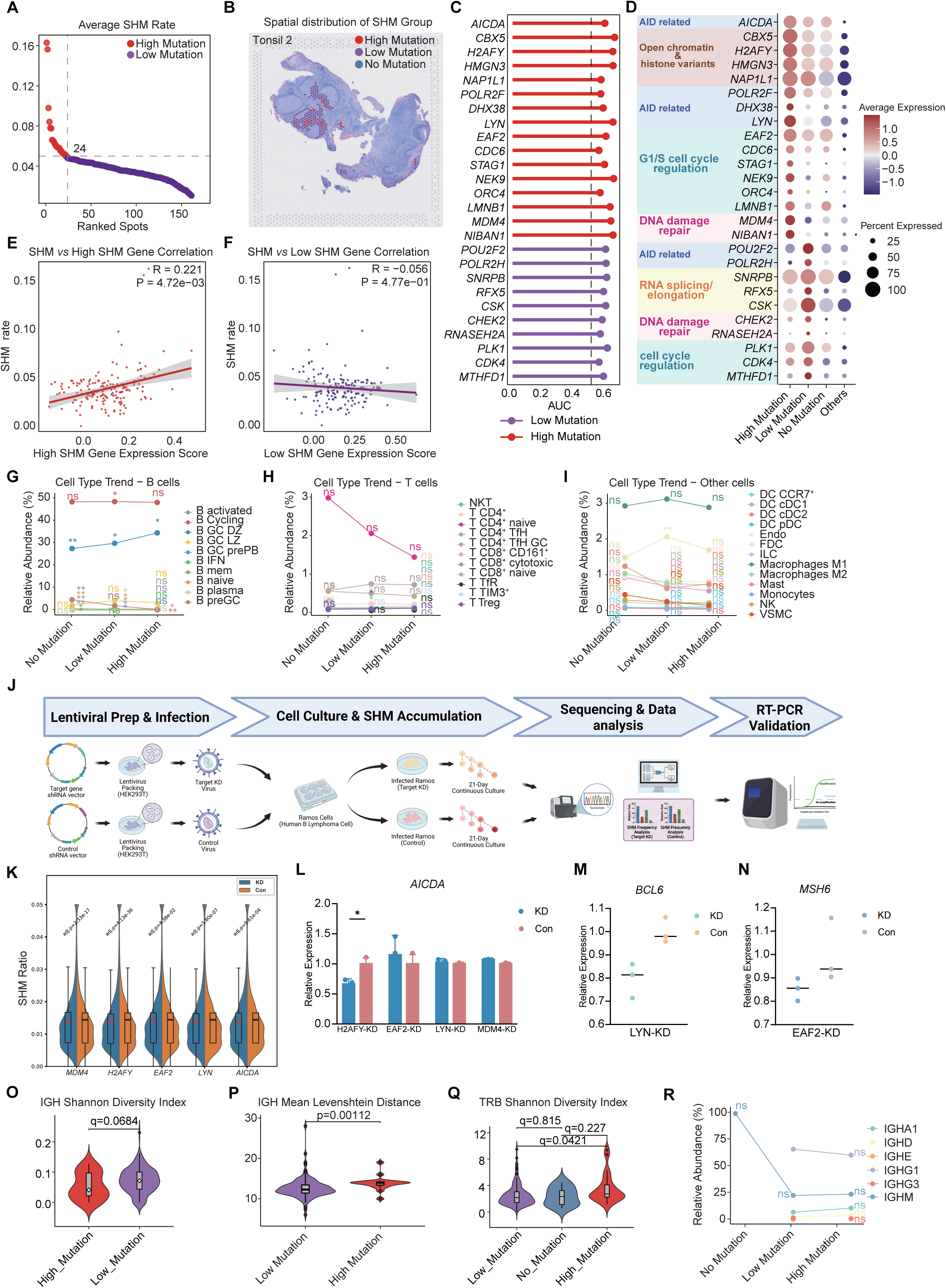
Intrinsic programs associated with SHM intensity in GC dark zones and their functional validation. (A, B) Ranking of average SHM rates and spatial mapping of SHM groups within Tonsil 2. (C, D) AUC-based prioritization and expression patterns of genes associated with high- and low-SHM states. (E, F) Correlations between SHM rate and high- or low-SHM gene-expression scores. (G–I) Changes in B-cell, T-cell, and non-lymphoid cell abundance across SHM groups. (J) Experimental workflow for lentiviral shRNA perturbation and SHM validation in Ramos cells. (K) SHM-frequency distributions after knockdown of candidate genes and the AICDA positive control. (L–N) RT-qPCR-based validation of candidate downstream molecular changes, including AICDA, BCL6, and MSH6. (O–R) Relationships between SHM intensity and IGH diversity, sequence dispersion, TRB diversity, and IGH isotype composition.

Most SHM-associated genes identified in this study, including chromatin remodelers, transcription/splicing factors, cell-cycle regulators, and DNA repair proteins, represent cell-intrinsic regulatory mechanisms that operate within the same cell rather than through intercellular signaling. SHM itself is a highly specialized diversification program restricted to GC B cells. Consistent with this, B cells accounted for ≥70% of DZ populations and were further enriched in high-SHM regions, while T cells decreased (Figure 5G-I, Figure S12G-I). Publicly available single-cell datasets further confirmed that the majority of these genes are predominantly expressed in B cells (Figure S11A). We therefore infer that these SHM-associated genes are predominantly expressed by B cells and are consistent with intrinsic molecular programs linked to SHM regulation in DZ regions. This process requires not only AID and a permissive transcriptional/epigenetic environment but also proper cell-cycle regulation to ensure the appropriate temporal window for mutation.

To validate the roles of candidate genes in the regulation of somatic hypermutation (SHM), we employed the Ramos cell line, a widely used in vitro model for ongoing SHM [31, 32], and generated shRNA-transduced Ramos cell populations targeting *MDM4*, *LYN*, *EAF2*, *H2AFY*, as well as *AICDA* as a positive control. After continuous culture for 21 days, the FR1–J region of the BCR heavy chain was sequenced in each knockdown group (Figure 5J). Compared with the control group, *AICDA* knockdown markedly reduced SHM levels, confirming that this experimental system can reliably and accurately capture changes in SHM. Further analysis showed that knockdown of *MDM4*, *LYN*, *EAF2*, and *H2AFY* also reduced SHM levels, suggesting that these candidate genes may participate in the regulation of SHM in Ramos cells (Figure 5K).

To investigate the molecular basis underlying these phenotypes, we further examined the expression of genes associated with SHM regulation. We first assessed *AICDA* expression in all four candidate-gene knockdown groups. Notably, *AICDA* downregulation was observed only in the *H2AFY* knockdown group, whereas no evident decrease in *AICDA* expression was detected following knockdown of *MDM4*, *LYN*, or *EAF2* (Figure 5L). This finding suggests that, among the candidate genes examined, the effect of *H2AFY* on SHM is most likely directly linked to the AID/AICDA axis. In contrast, *LYN* knockdown did not reduce *AICDA* expression but was accompanied by decreased *BCL6* expression (Figure 5M), suggesting that *LYN* may not regulate SHM by directly affecting *AICDA* expression, but rather indirectly by altering the germinal center-like molecular state required to sustain SHM. Similarly, *EAF2* knockdown did not lead to reduced *AICDA* expression, but was associated with decreased *MSH6* expression (Figure 5N), suggesting that its point of action may lie in the mismatch repair pathway and subsequent processing steps downstream of AID-induced lesions. By comparison, although *MDM4* knockdown also produced a reduced-SHM phenotype, no molecular change sufficient to explain this phenotype was detected within the current scope of analysis, and its precise mode of action remains to be determined.

Taken together, these results suggest that different candidate genes may regulate SHM at distinct levels, including SHM initiation, maintenance of cellular state, and post-lesion repair and processing. These functional validation experiments not only highlight the complexity of the SHM regulatory network, but also further support the reliability and potential utility of SpaCir-VDJ for identifying functional VDJ-associated regulatory factors.

At the clonal level, we next evaluated how SHM intensity was associated with repertoire architecture. High-SHM DZ-associated regions showed reduced IGH diversity together with increased sequence dispersion (higher mean nearest Levenshtein distance) (Figure 5O-P, Figure S12J), whereas TRB diversity was increased (Figure 5Q, Figure S12K). This pattern suggests a relative divergence between BCR and TCR compartments, with B-cell lineages appearing more selectively focused while T-cell clonotypes remain broader in these regions. Notably, the proportion of IGHG1 was decreased in the high-SHM group, whereas IGHM and other isotypes showed a slight increase (Figure 5R, Figure S12L), raising the possibility that some highly mutated clones may not undergo efficient selection and/or productive class-switch recombination.

### SpaCir-VDJ Delineates Spatial TCR/BCR Distributions and Candidate Immune-State Associations within the Tumor Microenvironment

We next applied SpaCir-VDJ to human gastric cancer tissues. Previous studies have shown that the spatial organization and functional states of tertiary lymphoid structures (TLS) and tumor-infiltrating lymphocytes are associated with clinical outcome in gastric cancer [33–35]. However, the spatial topology of T cell and B cell clonotypes and their relationship to local tumor heterogeneity remain incompletely defined. We analyzed gastrectomy samples from four patients who had not received chemotherapy or neoadjuvant treatment, and generated matched spatial gene-expression and TCR/BCR repertoire libraries for integrated analysis (Figure 6A). Based on spatial transcriptomic profiles, we annotated major tissue domains, including tumor epithelial regions, stromal regions, TLS, plasma cell–rich regions, and myeloid-enriched regions (Figure S13A-D). Across the four samples, we detected on average more than 500 unique BCR clonotypes and nearly 1,000 unique TCR clonotypes per sample (Figure 6B). BCR clonotypes were predominantly enriched in TLS and plasma cell–rich regions, which also showed upregulation of immunoglobulin-related genes, whereas TCR clonotypes showed a more diffuse spatial distribution and greater inter-sample heterogeneity (Figure 6C, Figure S14A, and Figure S16A). Using a published gastric cancer single-cell RNA-seq reference[36], we further performed cell2location-based annotation of major cell populations, particularly immune cell compartments, across the four samples (Figure S13E). Inferred T-cell and B-cell abundance was positively correlated with the spatial distribution of TCR and BCR signals, respectively (Figure S14B and Figure S16B), supporting consistency between transcriptomic and repertoire-derived spatial patterns. In three of the four samples, plasma cell–rich regions were spatially adjacent to TLS and located near the tumor margin (Figure S13D). We therefore examined how BCR abundance and diversity varied with distance to the tumor boundary in plasma cell–rich and tumor-interior regions (Figure S14C). BCR signal was concentrated near the tumor boundary (Figure S14D), whereas the number of unique BCR clonotypes and the degree of clonal expansion decreased with increasing distance from the boundary (Figure 6D and Figure S14E). Differential expression analysis of plasma cell–rich regions near the tumor boundary further identified genes such as *FN1*, *MMP11*, and *CCL18* (Figure 6E), consistent with the presence of fibroblast- and macrophage-associated immunoregulatory features at this interface.

**Figure 6.**
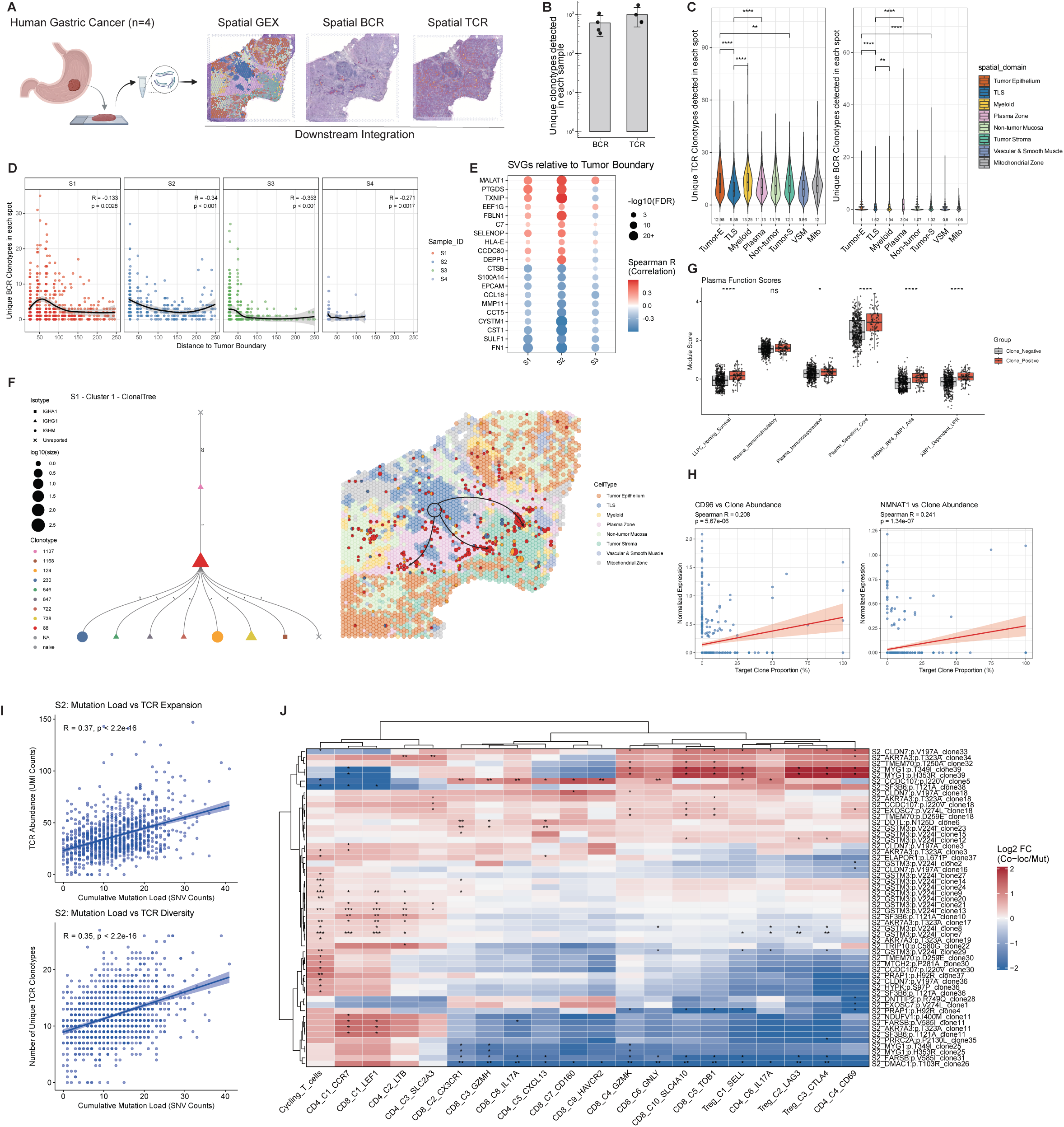
Spatial TCR/BCR distributions and candidate immune-state associations in gastric cancer. (A) Overview of the gastric-cancer SpaCir-VDJ analysis workflow. (B) Numbers of unique BCR and TCR clonotypes detected across gastric cancer samples. (C) Unique TCR and BCR clone numbers detected in each spatial domain across 4 samples. (D) Relationship between distance to the tumor boundary and local BCR clonotype richness. (E) Spatially variable genes associated with proximity to the tumor boundary. (F) Example clonal family showing spatial extension from TLS through plasma cell–rich regions toward the tumor interior. (G) Functional scores associated with clone-positive versus clone-negative spots for a representative dominant clone. (H) Correlations between CD96 or NMNAT1 expression and the proportion of clone 88 in plasma cell–rich regions. (I) Associations between cumulative tumor mutation load and TCR expansion/diversity in sample 2. (J) Heatmap showing the impact of different SNV mutation-TCR clonotype pairing relationships (in sample 2) on the abundance of T cell subtypes annotated by cell2location.

To further explore lineage structure, we reconstructed BCR clonal families in all four samples using fastBCR (Figure S15A, B). In sample 1, clone family 1 showed a spatial pattern extending from TLS toward the adjacent plasma cell–rich region and into the tumor interior (Figure 6F). Within this family, spots harboring the most expanded clone (clone 88) in the plasma cell–rich region showed upregulation of genes related to antibody secretion and immunoregulatory signaling (Figure 6G and Figure S15C). In addition, the abundance of this clone was positively correlated with genes including *CD96* and *NMNAT1* (Figure 6H), suggesting association with a distinct local immune context [37–39]. By contrast, clone family 2 displayed a different spatial configuration, with lineage-related spots present in both TLS and plasma cell–rich regions and showing features consistent with a more persistent plasma-cell program (Figure S15D-G). Together, these analyses highlight spatial heterogeneity in the distribution and transcriptional context of different BCR clone families.

In contrast to BCR, TCR showed comparatively smaller differences in quantity and diversity across spatial domains of the tumor, with aggregation observed in regions such as tumor tissue, TAM-rich regions, and TLS (Figure 6C and Figure S16A), indicating substantial T-cell infiltration. To explore whether local mutational landscapes might be associated with spatial TCR features, we used SpatialSNV to identify candidate mutation sites within tumor regions and applied stringent filtering criteria to derive cumulative mutation scores for each spot (see Methods) [40]. With the exception of sample 4, which showed lower sequencing quality, the remaining three samples exhibited a positive correlation between cumulative mutations in tumor epithelial regions and TCR expansion and diversity (Figure 6I and Figure S16C). This pattern is consistent with a spatial association between local mutational burden and TCR remodeling, but does not by itself establish tumor antigen-specific recognition. To further explore candidate antigen associations, we screened TCR clonotypes co-localized with mutation-enriched spots (see Methods) and integrated these results with cell2location-based annotations to examine relationships between mutation-TCR co-localization patterns and local T-cell states. These analyses revealed spatial differences in naïve and effector T-cell distributions among TCR groups with distinct candidate associations in the gastric cancer microenvironment (Figure 6J and Figure S16D). Furthermore, we used deepAntigen to predict putative interactions between candidate antigen epitopes and TCR CDR3 sequences (see Methods) [41]. Three clonotypes received higher prediction scores than non-colocalized controls and were associated with an increased proportion of cycling T cells with an expanded phenotype (Figure S16E). These observations nominate candidate neoantigen-associated clonotypes for future validation, but do not constitute direct evidence of antigen specificity. Together, these results provide a spatially resolved view of TCR and BCR distributions and candidate immune-state associations within the gastric cancer microenvironment.

## Discussion

Establishing a direct connection between immune clonal lineages and their native spatial organization is essential for dissecting the spatiotemporal dynamics of adaptive immune responses. However, most spatial transcriptomics (ST) assays are constrained by 3′-end capture chemistry and the low abundance of immune receptor transcripts. Moreover, in standard ST library architectures, clonotype-defining V(D)J features are physically distant from spatial barcodes, making it difficult to recover CDR3-containing sequences using short-read sequencing and thereby limiting in situ clonotype resolution. To address these challenges, we developed SpaCir-VDJ, an integrated workflow that enables joint profiling of whole-transcriptome states and spatially indexed immune receptor sequences within the standard 10x Genomics Visium pipeline. By introducing a circularization-based library design that brings distal receptor information into short-read–addressable proximity of spatial barcodes, SpaCir-VDJ provides a broadly applicable and cost-efficient approach for spatial immunogenomics.

SpaCir-VDJ addresses several practical bottlenecks in current technologies for spatial immune lineage analysis. Computational strategies that project bulk TCR/BCR sequencing data onto spatial maps, such as TRUST4-based approaches [42], generally require parallel assays and may be sensitive to variability introduced during cross-modal integration. Bead-based methods and other Visium-compatible workflows, including Slide-TCR-seq and related multimodal spatial barcoding workflows [43, 44], enable direct capture of immune receptor information but often rely on longer read configurations (for example, MiSeq PE300), which can increase per-sample cost and limit scalability for large-cohort studies. Long-read platforms such as Spatial VDJ [15] and SPTCR-seq [45] can span longer receptor segments, but are more susceptible to base-level sequencing errors, which may inflate apparent clonal diversity and compromise inference of somatic hypermutation (SHM) and isotype usage. In addition, workflows such as Stereo-XCR-seq typically adopt a circularization-first strategy [46]. Because circularization of low-abundance templates may result in efficiency loss, such designs are more prone to stochastic molecular loss before PCR amplification, thereby reducing sensitivity for rare clonotypes. In contrast, SpaCir-VDJ adopts an enrichment-first, circularization-later design. By recovering low-abundance receptor templates before circularization, this workflow may reduce stochastic molecular loss and better preserve tissue-level clonal complexity.

An additional strength of SpaCir-VDJ is its compatibility with the broader class of poly(A)-capture spatial transcriptomics platforms. Although implemented here using the current Visium format, the underlying workflow should be directly adaptable to higher-resolution systems, including 10x Genomics Visium HD (3′) and other array-based platforms, thereby extending spatial immune repertoire profiling from multicellular neighborhoods toward increasingly refined resolution. This forward compatibility positions SpaCir-VDJ to enable progressively finer spatial dissection of in situ immune clonal diversification as spatial resolution continues to improve.

Applying SpaCir-VDJ to lymphoid tissues revealed fine-grained spatial heterogeneity in immune clonal organization. SpaCir-VDJ resolves B-cell clonal maturation trajectories within germinal centers (GCs) and delineates isotype-associated patterns of somatic hypermutation (SHM), while shRNA-mediated knockdown in a Ramos B-cell model provides functional support for roles of *EAF2*, *H2AFY*, *MDM4*, and *LYN* in modulating SHM-associated phenotypes. In addition, the inferred inter-GC clonal redistribution events observed in our analyses suggest that GCs may not always behave as fully isolated reaction units. In gastric cancer, the platform further delineates spatial patterns in which dominant clonotypes are enriched in tertiary lymphoid structures (TLS) and tumor-margin niches, with lineage structures consistent with extension toward the tumor interior. We also observed a positive association between local mutational burden and spatial TCR diversity, a pattern compatible with links between tumor mutational landscapes and immune remodeling. Together, these findings position SpaCir-VDJ as a useful framework for interrogating immune response dynamics in complex tissue architectures.

SpaCir-VDJ also provides clear opportunities for further expansion. Although the current workflow robustly captures receptor segments spanning FR3–J, applications requiring more complete V(D)J information, including FR1, will benefit from further optimization of multiplex primer design and library architecture. In addition, the present study focuses primarily on TRB and IGH, which in part reflects the resolution boundary of the current implementation: at spot-level resolution, robust spatial pairing of additional chains such as TRA and IGL remains challenging. Importantly, because SpaCir-VDJ operates on poly(A)-captured, spatially barcoded cDNA, the workflow should be readily transferable to higher-resolution platforms, including newly developed single-cell–resolution spatial transcriptomics systems such as 10x Genomics Visium HD (3′). Continued advances in both spatial transcriptomics platforms and associated computational frameworks should therefore further extend SpaCir-VDJ toward more comprehensive in situ immune receptor pairing and finer dissection of clonal diversification.

In summary, SpaCir-VDJ provides a scalable and workflow-efficient framework for spatial immune repertoire profiling. By enabling in situ detection of clonal expansion, supporting SHM-associated diversification analyses, and linking transcriptional phenotypes with spatially indexed clonotypes, SpaCir-VDJ expands the experimental toolkit for interrogating immune organization and dynamics in complex tissues, with broad applications in tumor immunity, autoimmunity and spatially informed immunotherapy development.

## Materials and Methods

### Human Tissue Specimen Collection

Human tissue specimens were obtained from two clinical centers in China. Two tonsil samples and two lymph node samples were collected from the First Affiliated Hospital of the University of Science and Technology of China (USTC, Hefei, China) with approval from the Institutional Clinical Ethics Committee (No. 2025KY-192). In addition, four gastric cancer (GC) specimens were obtained from Xijing Hospital (Xi’an, China) under clinical ethics approval No. KY20254023-1. All procedures involving human participants were conducted in accordance with the Declaration of Helsinki, and written informed consent was obtained from all individuals before enrollment.

Immediately after surgical excision, specimens were rinsed in ice-cold PBS to remove residual blood and debris, embedded in optimal cutting temperature (OCT) compound, snap-frozen on dry ice, and stored at −80 °C until further processing for spatial transcriptomic and immune repertoire profiling. All sample numbers used throughout the study refer to internal specimen identifiers assigned during sample processing and are retained for consistency with laboratory records; therefore, the numbering is not necessarily consecutive and does not reflect the total number of analyzed specimens.

### Spatial Visium Library Preparation and Sequencing

Spatial transcriptomic libraries were prepared using the Visium Spatial Gene Expression Slide & Reagent Kit (10x Genomics). Briefly, 10-μm-thick cryosections were mounted onto Visium slides, and library preparation was performed according to the manufacturer’s instructions (Document No. CG000239 Rev G) with minor modifications. Based on optimization using the Tissue Permeabilization Kit, the permeabilization time was set to 12 min for most samples, whereas LN 2 (internal sample identifier) was permeabilized for 18 min. Libraries were sequenced on the DNBSEQ-T7 platform (MGI).

### SpaCir-VDJ Library Construction and Sequencing

To capture TCR and BCR repertoires from Visium-derived cDNA, 1 μL of the cDNA library was used as input for targeted amplification. V-region amplification was performed using multiplex forward primer pools targeting human TRB and human IGH FR3 regions, together with a universal reverse primer (5′-CTACACGACGCTCTTCCGATCT-3′). The primer sequences have been described previously [16, 17]. Following PCR enrichment, TCR and BCR libraries were size-selected and purified using SPRIselect beads (Beckman Coulter, B23318).

For library circularization, 100 ng of the PCR-enriched TCR/BCR library was processed using the Cyclization Kit (13341ES, Yeasen Biotechnology). The splint oligonucleotide provided in the kit was replaced with a custom-designed oligonucleotide (5′-GCGTCGTGTAGACACGACGCTC-3′). Circularized products were subsequently amplified using Multiplex PCR Enzyme (PM101, Vazyme) to enrich fragments containing both the spatial barcode and the CDR3 region. Primer sequences used in this step are listed in Supplementary Data. PCR was performed under the following conditions: 95 °C for 5 min; 30 cycles of 95 °C for 30 s, 60 °C for 90 s, and 72 °C for 30 s; followed by 72 °C for 10 min. Final libraries, with a target size of approximately 300 bp, were purified using SPRIselect beads and quantified using Qubit (Thermo Fisher Scientific). Libraries were prepared for sequencing using the VAHTS Universal DNA Library Prep Kit for MGI (NDM607, Vazyme), supplemented with specific adapters and indices (13350ES01, Yeasen Biotechnology), and sequenced on the DNBSEQ-T7 platform using a PE150 strategy. Read 1 captured the spatial barcode, whereas Read 2 covered the CDR3 and J regions.

### TRB/IGH Amplicon Preprocessing and V(D)J Clonotype Calling

Paired-end FASTQ files from TRB and IGH amplicon libraries were processed using pRESTO (v0.7.2; FilterSeq.py and MaskPrimers.py), MiXCR (v4.6.0; repseqio.v4.0 reference library)[47], and custom Python scripts (Python 3.9.13; Biopython 1.85; pandas 2.2.2; NumPy 1.24.4; matplotlib 3.9.4). Processing steps included read-level quality filtering, primer tagging, orientation normalization, poly(A)-based selection and trimming, read-pair concatenation, V(D)J alignment, and clonotype assembly.

Reads were first filtered using pRESTO with a minimum Phred score threshold of 20, followed by primer identification and tagging. Primer-tagged paired-end reads were then grouped by read identifier and classified into valid forward/reverse primer orientations. Poly(A)-containing molecules were enriched by requiring a poly(A) stretch in read 1, followed by trimming of the upstream sequence and corresponding qualities. For each sample, valid read orientations were merged, and read 2 was reverse-complemented and concatenated to read 1 to reconstruct amplicon sequences.

Concatenated reads were aligned and assembled into clonotypes using MiXCR with human RNA amplicon settings. For TRB, the floating left alignment boundary was set to VEnd; for IGH it was set to FR1Begin, together with permissive alignment settings to retain partially aligned molecules when necessary. Alignment tables were post-processed in Python to remove non-clonal reads (cloneId = −1) and to retain only chain-consistent records (TRB for TRB libraries and IGH for IGH libraries), yielding filtered alignment tables for downstream analyses.

Additional preprocessing parameters and implementation details are provided in the Supplementary Methods.

### Spatial T/BCR Barcode Matching

To link MiXCR-derived V(D)J alignments to Visium spatial barcodes, filtered alignment tables were matched to the Cell Ranger/Space Ranger barcode whitelist (barcodes.tsv) using a custom barcode-recovery workflow. Original Visium barcodes were converted into the orientation compatible with barcode sequences embedded in repertoire reads by removing the “-1” suffix and taking the reverse complement.

For each MiXCR alignment record, candidate barcode windows were scanned within the target sequence and matched against the whitelist. Exact matches were retained directly, whereas single-edit barcode correction was allowed under quality constraints. UMIs were extracted relative to the matched barcode window, and matching/correction was executed in parallel across records. Read-level annotations included matched barcode, correction type, edit distance, barcode-window quality, and UMI sequence.

At the spatial-barcode level, reads assigned to the same barcode were aggregated, and the corresponding original Visium barcode notation was restored. Downstream filtering retained only barcode assignments with edit distance ≤ 1, removed low-support events (<3 reads per UMI), and resolved ambiguous assignments by keeping the entry with the highest cumulative support for each (spatial barcode, UMI) combination. This procedure generated spot-resolved TCR/BCR clonotype and UMI count tables for subsequent spatial analyses.

Detailed barcode-recovery procedures, UMI extraction rules, and filtering criteria are provided in the Supplementary Methods.

### Visium Spatial Transcriptomics Preprocessing, Clustering, and Cell Type Mapping Visium preprocessing, dimensionality reduction, and clustering

Visium spatial transcriptomics data were processed using Seurat v5.1. For each sample, the filtered expression matrix (outs/filtered_feature_bc_matrix.h5) was loaded using Load10X_Spatial() with the Spatial assay as default. Mitochondrial read percentage per spot was calculated using PercentageFeatureSet() with the pattern ^MT-. Data were normalized and variance-stabilized with SCTransform (method = “glmGamPoi”), regressing out mitochondrial content (vars.to.regress = “percent.mt”). Principal component analysis was performed on SCT-transformed data using the top 30 principal components, followed by shared nearest-neighbor graph construction (FindNeighbors, dims = 1:30), Leiden clustering (FindClusters, algorithm = 4), and UMAP visualization (RunUMAP, dims = 1:30). Clustering resolution was adjusted per sample according to dataset complexity.

### Cell type mapping with cell2location

Cell type mapping of Visium data was performed using cell2location (v0.1.3) [19]. For tonsil and lymph node samples, reference cell-state signatures were derived from the tutorial-provided single-cell reference dataset distributed with cell2location (sc.h5ad) together with its pretrained RegressionModel. For gastric cancer samples, reference signatures were derived from a published human gastric cancer single-cell RNA-seq dataset [36]. The Visium Seurat objects were converted to AnnData format, and gene symbols were mapped to Ensembl identifiers using biomaRt to ensure compatibility with the corresponding reference. Mitochondrial genes were flagged and excluded from the feature set used for cell2location fitting, while remaining in the object for quality-control bookkeeping. In addition, the LN 1 sample lacked a well-defined germinal center (GC) structure based on both H&E staining and transcriptomic signals and was therefore excluded from subsequent GC-focused analyses.

Reference signatures were extracted as mean expression per cell state/cell type, and both spatial data and signatures were restricted to intersecting genes prior to model fitting. The model was trained on the full spatial dataset (batch_size = None, train_size = 1) using N_cells_per_location = 15 and detection_alpha = 20 to accommodate spot-to-spot technical variation in detection sensitivity, for 20,000 epochs. Posterior cell abundance estimates were obtained using export_posterior (num_samples = 1000), and spot-level abundance summaries were exported for downstream visualization and association analyses. Unless otherwise specified, q05_cell_abundance_w_sf was used as the spot-level abundance estimate for each cell type.

### Pseudotime trajectory inference and gene module analysis

Pseudotime and trajectory-associated gene module analyses were performed on LN 2 Visium spots [48], as LN 2 exhibited a well-defined GC architecture and a clear continuum across DZ, LZ, and Inter LZ-DZ regions. Raw spatial counts and spot-level metadata were extracted from the Seurat object, and a Monocle3 cell_data_set was generated with corresponding gene metadata. Based on Seurat-derived regional annotations, analysis was restricted to germinal center–related populations (Follicular DZ, Follicular LZ, and Inter LZ-DZ).

The subsetted cell_data_set was preprocessed using preprocess_cds (num_dim = 200), embedded with UMAP, clustered, and used to learn the principal trajectory graph. Spots were ordered using order_cells to obtain pseudotime values, which were then written back to Seurat metadata and visualized in tissue space. To identify trajectory-associated genes, graph_test was applied on the principal graph and significant genes (q < 0.05) were grouped into co-expression modules using find_gene_modules. Module-level expression was aggregated by regional annotation and visualized as module-by-cluster heatmaps. Top trajectory-associated genes ranked by Moran’s I were visualized along pseudotime to characterize continuous transcriptional programs associated with DZ/LZ transitions.

## Spatial Clonotype and Germinal Center Analyses

### Spatial proximity analysis of expanded IGH and TRB clonotypes

Using the barcode-matched spot × clonotype UMI count matrix, clone size for each clonotype was defined as the total UMI count summed across all spots. IGH and TRB clonotypes were categorized into four expansion groups: Not Expanded (clone size = 1), Small (2 ≤ clone size < 6), Medium (6 ≤ clone size < 20), and Hyperexpanded (clone size ≥ 20). To quantify spatial clustering, spatial coordinates of all spots containing each clonotype were extracted, pairwise Euclidean distances were computed, and the nearest-neighbor distance for each spot was determined. The mean nearest-neighbor distance for each clonotype was defined as the average of these values across occupied spots (set to 0 for clonotypes present in only one spot). Group-level comparisons excluded the Not Expanded group to avoid inflation by single-spot clonotypes and were assessed using two-sided Wilcoxon rank-sum tests.

### Clone family clustering, lineage reconstruction, and class-switch analysis of spatial IGH repertoires

IGH sequences were analyzed using fastBCR v1.1 for clone family clustering and clonal lineage inference [49]. Representative CDR3 amino acid sequences were used to summarize UMI support, and each UMI-supported event was indexed by a composite sequence identifier consisting of spatial_barcode_ori + UMI. Clone families were identified using data.BCR.clusters() with cluster_thre = 3, overlap_thre = 0.1, and consensus_thre = 0.8. Representative families were visualized by multiple-sequence alignment and sequence logo analysis. Clonal lineage trees were reconstructed using ClonalTree through the fastBCR interface, generating Newick-formatted trees for downstream visualization. For class-switch recombination (CSR) analyses, within-family isotype composition and inferred switching patterns were summarized using fastBCR plotting functions.

To infer lineage-consistent spatial redistribution of IGH clones, lineage directionality along reconstructed clonal trees was integrated with the spatial coordinates of the corresponding UMI-supported sequence events. This approach enabled visualization and interpretation of clonal expansion and redistribution patterns across GC regions based on the localization of successive lineage states.

To quantify similarity of IGH clonotype composition across GC subregions (GC_part), spatial clonotype information was aggregated from the spot level to the GC_part level. Pairwise similarity between GC_part groups was measured using the Jaccard similarity coefficient, and similarity was converted into a distance metric as D(A,B) = 1 − J(A,B). The resulting distance matrix was subjected to hierarchical clustering and visualized as a clonal similarity tree.

### GLIPH2-based clustering and public-database annotation of spatial TRB CDR3β repertoires

Spatial TRB CDR3β amino acid sequences were functionally clustered using turboGliph v0.99.2 (GLIPH2) [50]. CDR3β sequences together with their TRBV/TRBJ assignments were formatted as GLIPH2 inputs, and UMI-deduplicated unique counts were used as sequence support. Clustering and motif-based scoring were performed with GLIPH2.0 settings using the built-in reference database and V-gene matching.

For antigen/disease annotation, human entries from VDJdb [51] and McPAS-TCR [52] were curated and merged into a unified reference table containing harmonized fields for CDR3β, TRBV, TRBJ, and pathology/epitope metadata. GLIPH2 cluster members were annotated by exact CDR3β sequence matching against the merged reference.

## Spatial clonotype stratification and GC maturity assignment

### Spatial IGH/TRB clonotype categorization by sharing breadth and clonal expansion

Spatial IGH/TRB clonotypes were stratified along two axes: sharing breadth across GC regions and degree of clonal expansion. For each clonotype, we summarized the number of distinct GC subregions in which it was detected (num_GC_sites, counted across GC_part) and its total UMI support (total_UMI). Clonotypes were dichotomized as expanded versus unexpanded using the 95th percentile of total_UMI as the expansion threshold, and classified into four groups: Private Expanded, Shared Expanded, Private Unexpanded, and Shared Unexpanded.

### GC maturity labeling using stage-associated gene signatures

To infer GC maturity states in spatial transcriptomic profiles, spots annotated as non-GC (GC_part = “Others”) were removed. GC maturity was then assigned using published stage-associated gene sets. The Early GC signature comprised CD83, CD86, CXCR5, GPR183, SLAMF1, and BCL6. The Late GC signature comprised IRF4, PRDM1, XBP1, ZBTB20, DUSP2, IRF8, GADD45B, JCHAIN, TOX2, and SDC1. Module scores for the Early and Late signatures were computed per spot using Seurat AddModuleScore (nbin = 10) [21–25]. For each GC_part, mean Early and Late module scores were summarized across spots, and a maturity label (EarlyGC or LateGC) was assigned according to the higher mean score.

### Differential expression and GO enrichment to support maturity grouping

To evaluate the biological coherence of the maturity assignment, differential expression analysis was performed between EarlyGC and LateGC groups using Seurat FindAllMarkers (only.pos = TRUE, min.pct = 0.1). Upregulated genes were filtered by adjusted P value < 0.05 and avg_log2FC > 0.5, and mitochondrial genes were excluded. To avoid circularity, maturity marker genes used for classification were removed from the DEG lists prior to enrichment analysis. GO enrichment was performed using clusterProfiler v4.12.0 and org.Hs.eg.db v3.19.1, retaining terms with p.adjust < 0.05.

### GC maturity–stratified clonal composition and diversity analysis

To visualize stage-wise clonal composition, UMI counts were aggregated per stage × clonotype and converted to within-stage relative abundances. The top 100 clonotypes in LateGC, ranked by relative abundance, were selected and tracked across stages, with all remaining clonotypes collapsed into an “Other” category. Relative clonotype composition was displayed using stacked bar plots in the order EarlyGC → LateGC.

To compare repertoire concentration and diversity across stages, clonotypes within each stage were ranked by decreasing relative abundance and cumulative frequency curves were calculated and plotted on a log10 x-axis.

### Stage-wise compositional trend analysis

To quantify GC maturity–associated changes in immune features, including isotype usage, cell-type abundance composition, and clone-class composition, spot-level raw signals were first calculated. For visualization only, raw signals were converted into within-stage compositional percentages by normalizing each category to the total signal within that stage. Stage-wise compositional trajectories were displayed using line plots or stacked composition plots.

Statistical analyses were performed at the spot level, the standard analytical unit for Visium-based spatial transcriptomic data. For prespecified contrasts, including EarlyGC versus LateGC comparisons, two-sided Wilcoxon rank-sum tests were applied to spot-level raw measurements for each category independently. P values were used to support observed spatial trends across groups and were summarized on plots using conventional significance notation.

### Spatial HdWGCNA network construction and functional annotation

Spatial HdWGCNA was performed using HdWGCNA v0.4.05 on Seurat objects after excluding spots annotated as non-GC[53]. Genes were pre-selected using SetupForWGCNA with an expression-frequency strategy (gene_select = “fraction”, fraction = 0.05), and spatial spots were aggregated into metaspots by GC_Maturity_Label to reduce spot-level noise. The Spatial assay was used as network input, and soft-thresholding power was selected based on approximate scale-free topology. Networks were constructed using ConstructNetwork, followed by calculation of module eigengenes and module connectivity. Module eigengenes were added to Seurat metadata and visualized using DotPlot and SpatialFeaturePlot.

Module co-expression relationships were embedded using RunModuleUMAP. Hub genes were identified on the basis of module membership (kME) using GetHubGenes, retaining the top 150 hub genes per module. To avoid circularity, mitochondrial genes and maturity marker genes used for GC labeling were excluded before enrichment analyses. GO enrichment for filtered hub-gene lists was performed using clusterProfiler and org.Hs.eg.db, with significance defined as BH-adjusted p.adjust < 0.05. For visualization, a combined score integrating enrichment magnitude and statistical significance was computed as GeneRatio × [−log10(p.adjust)].

### Somatic hypermutation–stratified analysis of follicular DZ spots

Spatial spots annotated as Follicular DZ were extracted from the Seurat object for downstream analyses. SHM was quantified using MiXCR-derived IGH VDJ_mutation_rate. For each spot, SHM was summarized as the mean VDJ_mutation_rate across all IGH entries assigned to that spot (avg_15G_VDJ_mutation_rate). Spots were then ranked by this value and classified using an empirical cutoff of 0.05 into High_Mutation (>0.05), Low_Mutation (0–0.05), or No_Mutation (0). This three-level classification was stored in Seurat metadata as SHM_VDJ.

To characterize SHM-associated transcriptional programs within the DZ subset, differential feature screening across SHM_VDJ groups was performed using Seurat FindAllMarkers with the ROC test (test.use = “roc”), yielding an AUC-like discriminative metric (myAUC) for each gene. For each gene, the record with the maximal myAUC across groups was retained, and mitochondrial genes were excluded. SHM-associated genes were visualized using AUC-ranked plots, and predefined gene sets were displayed across SHM_VDJ groups using DotPlot. In addition, SHM-associated gene sets were scored per DZ spot using Seurat AddModuleScore, and module scores were correlated with spot-level SHM using Spearman rank correlation.

## SpatialSNV, TCR-SNV Pairing, and Neoantigen Prediction

### SpatialSNV data integration and candidate somatic variant prioritization

To characterize the spatial landscape of somatic alterations, spatial transcriptomic RNA profiles were integrated with spatial single-nucleotide variant (SNV) matrices using Scanpy (v1.9.0). Spatial coordinates were retained to preserve tissue architecture. Variant allele frequency and sequencing depth matrices were aligned to RNA-seq barcodes, and only shared spots were retained.

Variants were annotated using the SpatialSNV pipeline integrated with ANNOVAR and the refGene (hg38) database[40]. Functional consequences, including genomic region and mutation class, were assigned to each variant. SNV matrices were normalized against total RNA counts using normalize_with_rna to account for spot-to-spot variation in transcript capture efficiency.

To prioritize candidate somatic variants while reducing likely germline polymorphisms and technical artifacts, a multi-step filtering strategy was applied: (i) only exonic nonsynonymous SNVs resulting in amino acid substitutions were retained; (ii) spatial clustering was assessed using Moran’s I, and variants with Moran’s I > 0.1 were prioritized; and (iii) tumor-to-TLS enrichment was used as an additional filter, retaining variants with a tumor-to-TLS enrichment ratio > 2.0 or variants detected exclusively within tumor-rich domains.

Candidate variants were further resolved at the transcript level by expanding ANNOVAR annotations into one-to-many mappings to specific Ensembl/RefSeq transcripts. The resulting candidate variant matrix and associated metadata were exported for downstream spatial co-localization analyses with TCR repertoires.

Detailed variant-filtering criteria, spatial co-localization procedures, and neoantigen prediction settings are described in the Supplementary Methods.

### Spatial co-localization and TCR-SNV pairing

To identify candidate interactions between SNVs and TCR clonotypes, SNV and TCR matrices were integrated into a unified Seurat object. A UMI-based spatial pairing algorithm was used to calculate UMI precision, defined as the proportion of co-localized UMIs within shared capture spots. High-confidence pairs were retained using the following criteria: UMI precision ≥ 0.5, spatial co-localization P < 0.05 by permutation testing, and at least three co-localized UMIs.

### Immune perturbation and cell-type specificity

To evaluate the potential immune-context changes associated with specific TCR–mutation engagement, cell-type abundances per spot derived from cell2location were compared between interaction sites and background regions. For each validated pair, log-fold changes of specific cell lineages were calculated.

### Predicting TCR specificity using deepAntigen

Based on candidate variants identified by SpatialSNV, corresponding protein sequences were retrieved from UniProt, and a 20-residue window flanking each mutation site was extracted. Candidate 9-mer and 10-mer neoepitopes containing the substituted residue were generated using a sliding-window approach. Candidate neoantigens and their spatially co-localized TCR clonotypes were exhaustively paired and evaluated using deepAntigen to generate interaction scores[41].

### Ramos Cell Perturbation and SHM-Associated Functional Assays Cell culture

Ramos cells (Cat. No. CL-0483; Procell Life Science & Technology Co., Ltd.) were cultured in RPMI-1640 medium supplemented with 10% fetal bovine serum (FBS) and 1% penicillin/streptomycin (P/S) in a humidified incubator at 37 °C with 5% CO2.

### Construction of shRNA lentiviral vectors

To generate shRNA-expressing lentiviral vectors, the pLL3.7 backbone maintained in our laboratory was linearized by dual digestion with HpaI and XhoI (Takara). Target-specific shRNA oligonucleotides (Sangon Biotech; sequences listed in Supplementary Data) were resuspended in annealing buffer (10 mM Tris-HCl, pH 8.0, 50 mM NaCl, 1 mM EDTA). Sense and antisense oligonucleotides were mixed at an equimolar ratio, heated to 95 °C for 5 min, and gradually cooled to room temperature (25 °C) at a rate of 0.1 °C/s to form double-stranded inserts.

Annealed inserts were ligated into the linearized pLL3.7 vector at a 5:1 molar ratio using T4 DNA ligase (Takara) at 16 °C overnight. Ligation products were transformed into chemically competent *E. coli* Stbl3 cells. Positive clones were selected by ampicillin resistance, screened by colony PCR, and validated by Sanger sequencing using a U6 promoter–specific forward primer.

### Lentivirus production

Lentiviral particles were generated by transient transfection of HEK293T cells using a second-generation packaging system. Briefly, HEK293T cells were seeded in 10-cm dishes and transfected at 70–80% confluence with pLL3.7, psPAX2, and pMD2.G at a mass ratio of 4:3:1. For each dish, 15–20 μg total DNA was mixed with Opti-MEM (Gibco), and linear PEI (MW 25,000) was added at a 3:1 (w/w) ratio relative to DNA. Complexes were incubated for 15–20 min at room temperature before being added dropwise to the cells.

After 16 h, the medium was replaced with fresh complete DMEM supplemented with 10% FBS. Viral supernatants were collected 48 h post-transfection, cleared by centrifugation at 3,000 rpm for 5 min, and filtered through a 0.45-μm PVDF low-protein-binding filter (Millipore). Filtered viral supernatants were aliquoted and stored at −80 °C until use. Repeated freeze-thaw cycles were avoided.

### Lentiviral transduction

Ramos cells were seeded in 6-well plates at 2 × 10^5 cells per well in complete RPMI-1640 medium. Lentiviral supernatant was added together with 8 μg/mL polybrene (Santa Cruz Biotechnology), and plates were centrifuged at 800 × g for 1 h at 30 °C to enhance virus-cell contact. After 6 h of incubation, the virus-containing medium was replaced with fresh complete medium. No antibiotic selection was applied. Successful transduction was confirmed by fluorescence microscopy after infection, indicating that the majority of cells were transduced. Knockdown efficiency was evaluated by RT-qPCR at approximately 5–7 days post-infection. Downstream analyses were performed on pooled transduced cells, and the 21-day culture period for SHM-associated assays was counted from day 1 after infection.

### RNA extraction and RT-qPCR

Total RNA was extracted from Ramos cells using the FastPure Cell/Tissue Total RNA Isolation Kit (Vazyme Biotech) according to the manufacturer’s instructions. RNA concentration and quality were assessed using a NanoDrop 2000 spectrophotometer (Thermo Fisher Scientific). For reverse transcription, 500 ng total RNA was used to synthesize cDNA in a 20-μL reaction using PrimeScript RT Master Mix (Perfect Real Time; Takara Bio).

Quantitative PCR was performed using TB Green Premix Ex Taq II (Tli RNaseH Plus; Takara Bio) on a CFX96 Real-Time PCR Detection System (Bio-Rad). Cycling conditions were 95 °C for 30 s, followed by 40 cycles of 95 °C for 5 s and 60 °C for 30 s, followed by melting-curve analysis. Relative mRNA levels were normalized to TBP and calculated using the 2^−ΔΔCt method. Primer sequences are listed in Supplementary Data. RT-qPCR analyses were performed using two independent biological replicates, each measured in technical triplicate.

### BCR repertoire library preparation and sequencing

Following lentiviral transduction, Ramos cells were cultured continuously for 21 days from day 1 post-infection to allow accumulation of somatic hypermutations (SHM) within the BCR locus. Knockdown efficiency was evaluated by RT-qPCR at approximately 5–7 days post-infection. Cells were monitored daily and passaged every 2 to 3 days in complete RPMI-1640 medium. At each passage, knockdown and control groups were reseeded at the same starting density to minimize density-dependent variation, and cultures were maintained at 2 × 10^5 to 1 × 10^6 cells/mL. Total RNA was subsequently isolated from each experimental group for downstream analysis.

BCR repertoire libraries were generated by targeted PCR amplification of the IGH variable region spanning framework region 1 (FR1) to the J region. The forward primer sequence was 5′-TGTCCCTCACCTGCGCTGTC-3′, and the reverse primer sequence was 5′-ATACCGTAGTAGTAGTAG-3′. The resulting amplicons were purified and used for downstream library construction and sequencing. Libraries were sequenced on the DNBSEQ-T7 platform (MGI) using a PE150 strategy.

## Acknowledgements

This work was supported by the Scientific Research Start-up Funds (QD2021005N), the Department of Chemical Engineering–iBHE Special Cooperation Joint Fund (DCE-iBHE-2025-2), and the Shenzhen Science and Technology Program (WDZC20220819134430002). The funders had no role in the study design, data collection and analysis, decision to publish, or preparation of the manuscript.

## Author Contributions

Zheyu Hu conceived the experimental strategy and developed the technical workflow. Danni Guo designed the bioinformatics analysis framework. Xiuyuan Wang contributed to bioinformatics analysis. Zheyu Hu, Xueting Li, and Maiyi Zhong performed the spatial transcriptomics experiments and associated validation assays. Xie Wang provided technical support. Zhaoxuan Zhang, Zizhao Gao, Zhe Wang, and Wei Wang contributed to clinical specimen collection and sample provision. Zhe Wang and Xiao Liu supervised the study and served as co-corresponding authors. Xiao Liu also contributed to overall study conception and funding acquisition. All authors read and approved the final manuscript.

---

**Figure S1. Additional characterization of SpaCir-VDJ in human tonsil and lymph node tissues.** (A) Study design showing analysis of two human tonsils and two lymph nodes using joint spatial gene-expression and spatial clonotype profiling. (B) H&E images of the four analyzed tissues. (C) Rarefaction and extrapolation curves showing near-saturation sequencing depth for IGH and TRB libraries. (D, E) Sample-wise IGHV and TRBV usage profiles. (F) Sample-specific SHM distributions across IGH isotypes. (G) Sample-wise nearest-neighbor distance analyses showing spatial clustering of larger IGH and TRB clonotypes.

**Figure S2. Additional spatial-correlation and signaling analyses for Tonsil 2.** (A, B) Heatmaps showing associations between IGH/TRB diversity and cell-type abundance across tissue regions. (C, D) Global CellChat interaction strengths and signaling-network organization across annotated regions. (E, F) Contributions of BAFF-, MHC-II-, and CXCL-related ligand–receptor pairs to regional communication. (G) Spatial maps illustrating co-localization of selected B-cell and T-cell subsets within the LN 2 and Tonsil 2 tissue.

**Figure S3. Regional annotation and repertoire features in LN 2.** (A, B) UMAP- and tissue-space annotation of follicular, inter-follicular, epithelial/muscle, and subepithelial/mesenchymal/vascular compartments. (C) Marker-gene expression supporting regional definitions. (D–F) Region-specific enrichment of representative immune and stromal cell populations. (G, H) Regional IGH/TRB diversity and SHM patterns. (I) Correlations between repertoire diversity and cell-type abundance. (J, K) Associations between T-cell subset diversity and B-cell programs.

**Figure S4. Quality filtering, regional annotation, and repertoire analyses in Tonsil 1.** (A) Quality-control distributions and spatial maps before and after filtering of low-capture regions. (B, C) UMAP- and tissue-space annotation of follicular, inter-follicular, epithelial-like, and subepithelial/mesenchymal/vascular regions. (D) Regional marker-gene expression. (E) Regional IGH/TRB diversity. (F) Correlations between regional repertoire diversity and cell-type abundance. (G) Associations between selected T-cell subset diversity and B-cell programs. (H) Heatmap summarizing correlations between T-cell and B-cell diversity programs.

**Figure S5. Additional pseudotime, lineage, and T-cell annotation analyses in Tonsil 2.** (A) Expression dynamics of representative pseudotime-associated genes spanning proliferation, affinity-selection, and plasmablast-transition states. (B–D) Sequence-logo and lineage-tree representations of selected IGH branches, together with spatial projection of branch transitions. (E–G) Relationships between clone classes, germinal-center identity, constant-region usage, and SHM-state association. (H) Expression of CSR-related genes in spots linked to putative GC re-entry versus other regions. (I) Representative spatial distributions of annotated TRB clusters with putative shared antigen associations.

**Figure S6. Detailed inter-GC clonal redistribution analyses in LN 2.** (A–D) Example LN 2 IGH clonal-family clustering, class-switch structure, CDR3-length distribution, and nearest-neighbor comparison. (E) Spatial annotation of germinal centers and non-GC regions in LN 2. (F, G) Sankey analyses summarizing inferred region-to-region and GC-to-GC clonal redistribution. (H, I) Clonal-similarity trees and shared-clone matrices across GC and non-GC compartments. (J, K) Quantification of inferred inter-GC event frequencies and spatial distances. (L, M) Relationships between shared expanded clonotypes, isotype composition, GC maturity, and SHM group. (N) CSR-related gene expression in GC re-entry–associated spots and comparison regions. (O–Q) Spatial clustering, pathology annotation, and representative spatial maps of TRB clusters in LN 2.

**Figure S7. Detailed inter-GC clonal redistribution analyses in Tonsil 1.** (A–D) Example Tonsil 1 IGH clonal-family clustering, class-switch structure, CDR3-length distribution, and nearest-neighbor comparison. (E) Spatial annotation of GCs and surrounding non-GC regions. (F, G) Sankey analyses summarizing redistribution across regions and across individual GCs. (H, I) Clonal-similarity trees and shared-clone matrices. (J) Quantification of spatial distances for inferred inter-GC events. (K, L) Associations of shared expanded clonotypes with GC identity and isotype composition. (M) CSR-related gene expression in candidate GC re-entry regions. (N–P) TRB-cluster spatial aggregation, pathology annotation, and representative spatial maps in Tonsil 1.

**Figure S8. Additional GC-level compositional and network analyses.** (A) Constant-region composition across individual GC parts. (B) Class proportions across GC parts. (C) Module-level enrichment between early and late GC states. (D) Spatial distribution of representative HdWGCNA modules. (E) GO-term enrichment of hub genes from maturity-associated modules.

**Figure S9. Detailed GC-maturity analyses in LN 2.** (A, B) Spatial map of individual GCs and class assignment of IGH clones according to GC breadth and abundance. (C) SHM frequencies across clone classes. (D, E) Constant-region and class composition across GC parts. (F–K) EarlyGC/LateGC scoring, GO enrichment, module activity, and representative module spatial patterns. (L–N) Diversity and clone-abundance distributions across GC maturity states. (O–Q) SHM, clone-class, and constant-region comparisons between early and late GCs. (R–T) Changes in B-cell, T-cell, and other cell-type abundances with GC maturity.

**Figure S10. Detailed GC-maturity analyses in Tonsil 1.** (A, B) Spatial map of germinal centers and class assignment of IGH clones across GC sites. (C) SHM frequencies across clone classes. (D, E) Constant-region and class compositions across GCs. (F–I) EarlyGC/LateGC abundance distributions and clone-abundance summaries. (J–L) SHM, clone-class, and constant-region comparisons between early and late states. (M–O) B-cell, T-cell, and other cell-type changes associated with GC maturation.

**Figure S11. Public single-cell validation of SHM-associated candidate genes.** (A) Dot plot showing expression of SHM-associated candidate genes across major immune and stromal cell populations, highlighting their predominant enrichment in B-cell compartments.

**Figure S12. Detailed SHM-group analyses in LN 2.** (A, B) Ranking and spatial projection of SHM groups in LN 2. (C, D) AUC-based gene prioritization and group-wise expression patterns for high- and low-SHM signatures. (E, F) Correlations between SHM rate and high- or low-SHM gene scores. (G–I) Changes in B-cell, T-cell, and other-cell abundances between SHM groups. (J–L) Effects of SHM intensity on IGH diversity, mean Levenshtein distance, TRB diversity, and IGH isotype composition.

**Figure S13. Spatial-domain annotation and cell-composition landscape of gastric cancer samples.** (A) UMAP representations of clustering, sample origin, and spatial-domain annotation across the four gastric cancer tissues. (B) Heatmap of top marker genes defining major gastric cancer cell types and spatial domains. (C, D) Tissue-space projections of cell-type and spatial-domain annotations for each sample. (E) Predicted cell-type composition across spatial domains using cell2location, with expanded summaries for plasma-cell-rich regions and TLS.

**Figure S14. Additional BCR spatial analyses relative to the tumor boundary.** (A) Distribution of unique BCR clonotypes across annotated spatial domains in each gastric cancer sample. (B) Correlation between detected BCR UMI counts and spatial cell-state abundance inferred from transcriptomic deconvolution. (C) Definition of spatial classes relative to the tumor boundary. (D) Distance distributions of unique BCR clone numbers within spots located internal and external to the tumor boundary. (E) Total BCR abundance in relation to distance from the tumor boundary.

**Figure S15. Detailed clonal-family analyses in gastric cancer BCR data.** (A) Cluster size of BCR families across all samples. (B) Class-switch structure of all clone families. (C, E) Differentially expressed genes between spots harboring the target BCR clone and background spots within the plasma cell region. (D) Representative clone tree and tissue projection for a spatially extended clonal family. (F) Comparison of plasma-cell functional scores between clone-positive and clone-negative regions. (G) Functional landscape of genes correlated with clone abundance.

**Figure S16. Additional TCR spatial analyses in gastric cancer.** (A) Distribution of unique TCR clonotypes across annotated spatial domains in each gastric cancer sample. (B) Correlation between detected TCR UMI counts and inferred T-cell abundance. (C) Sample-wise associations between cumulative mutation load and TCR expansion or diversity. (D) Heatmap showing the impact of different SNV mutation-TCR clonotype pairing relationships on the abundance of T cell subtypes annotated by cell2location.

