## Supplementary Methods for "SpaCir-VDJ: a broadly compatible circularization strategy for spatial immune repertoire profiling"

**Detailed Preprocessing of TRB and IGH Amplicon Libraries**

Paired-end FASTQ files derived from TRB and IGH amplicon libraries were processed using pRESTO (FilterSeq.py and MaskPrimers.py, v0.7.2), MiXCR (v4.6.0; built-in V/D/J/C library: repseqio.v4.0), and custom Python scripts (Python 3.9.13; Biopython 1.85; pandas 2.2.2; NumPy 1.24.4; matplotlib 3.9.4). Read processing included quality filtering, primer tagging, primer-orientation normalization, poly(A)-based selection and trimming, read-pair concatenation, V(D)J alignment, and clonotype assembly.

Reads were first filtered using FilterSeq.py quality with a minimum Phred score threshold of 20 (-q 20). Primer sequences were then identified using MaskPrimers.py align and recorded in read headers in tag mode (--mode tag) with BARCODE and PRIMER annotations (--barcode --bf BARCODE --pf PRIMER). Primer matching allowed a maximum error rate of 0.3 (--maxerror 0.3) and a maximum primer match length (--maxlen) of 27 for TRB and 20 for IGH.

Primer-tagged paired-end reads were parsed with Biopython (SeqIO) and grouped according to the core read identifier (the portion of the read ID preceding /1 or /2). Read pairs were classified into two valid orientations according to their PRIMER tags: Rprimer(/1)+Fprimer(/2) and Fprimer(/1)+Rprimer(/2). Pairs lacking a valid primer-tag combination, or carrying the same primer type on both ends, were discarded.

To enrich poly(A)-containing molecules, read1 sequences were retained only if the first 70 bp contained at least 8 consecutive adenines. Read2 entries were then filtered by retaining only those whose core IDs were present in the retained read1 set. Read1 sequences were trimmed by identifying, within the first 80 bp, the last occurrence of a run of at least 8 consecutive adenines; all bases upstream of and including that A-run were removed, and PHRED qualities were trimmed accordingly.

For each sample, valid read orientations were merged into a single combined read1 FASTQ and a single combined read2 FASTQ. Read2 was reverse-complemented and concatenated to read1 to reconstruct amplicon sequences for MiXCR alignment and clonotype assembly.

Concatenated reads were processed with MiXCR using the human RNA generic amplicon preset (-p generic-amplicon --species hsa --rna) with original reads retained (-OsaveOriginalReads=true) and no trimming-quality threshold (--trimming-quality-threshold 0). For TRB, --floating-left-alignment-boundary was set to VEnd; for IGH, it was set to FR1Begin, together with permissive settings (-OallowPartialAlignments=true -OallowNoCDR3PartAlignments=true), while the right boundary was set to C for both. Alignments were assembled into clonotypes with mixcr assemble --write-alignments, and per-read alignment details were exported using mixcr exportAlignments.

Exported alignment tables were post-processed in Python to remove non-clonal reads (cloneId = -1) and to retain only chain-consistent records, i.e., chains beginning with TRB for TRB libraries and IGH for IGH libraries.

**Detailed Spatial Barcode Recovery and UMI Extraction**

Filtered MiXCR alignment tables were matched to the Cell Ranger/Space Ranger barcode whitelist (barcodes.tsv) using a custom barcode-recovery pipeline. All whitelist barcodes were read while preserving the original -1 suffix. For each barcode, the suffix was removed and the remaining sequence was reverse-complemented, generating a two-column metadata table containing spatial_barcode_ori (original Visium barcode) and spatial_barcode_mixcr (reverse-complement barcode).

For each MiXCR alignment record, the first entry in targetSequences and its corresponding targetQualities were used. Candidate barcode windows were generated as 16-mers starting from the 6th nucleotide of the sequence and sliding with a 1-bp step across a search region of up to 27 bp. Each candidate window was compared with the whitelist set (spatial_barcode_mixcr).

Exact matches (Levenshtein distance = 0) were accepted directly. If no exact match was identified, a single-edit correction was attempted (Levenshtein distance ≤ 1). For mismatch-type edits (base substitutions), correction was performed only when all discrepant positions had PHRED quality scores < 30. Insertion/deletion events were recorded and labeled as insertion/deletion, together with the corrected sequence and matched whitelist barcode.

UMIs were extracted simultaneously for each read. Using the matched barcode window as an anchor, the 12 bp immediately upstream of that window were taken as the UMI; if fewer than 12 bp were available, the UMI was left-padded with adenines to a fixed length of 12. Matching and correction were executed in parallel across alignment records using a multithreaded ThreadPoolExecutor, generating a read-level table containing matched barcode, correction type, edit distance, barcode-window quality scores, and UMI sequence.

At the spatial-barcode level, all reads assigned to the same matched_spatial_barcode were aggregated by summing match_count and concatenating read-level annotations using brace-delimited strings to preserve read-level information. The aggregated table was then merged back to barcode metadata (spatial_barcode_mixcr → spatial_barcode_ori) to restore standard Visium barcode notation (including the -1 suffix). Additional consistency filtering was then applied as follows:
(i) only matches with edit distance ≤ 1 were retained;
(ii) within each spatial barcode, per-UMI read counts were computed and propagated back to each record;
(iii) events supported by fewer than three reads per UMI were removed; and
(iv) for each (spatial_barcode_ori, UMI) combination, the entry with the highest cumulative support within the corresponding cloneId was retained to reduce ambiguous assignments arising from barcode correction or sequencing noise.

**Additional Parameters for Cell2location Analysis**

Cell-type reference signatures were obtained according to tissue type. For tonsil and lymph node samples, reference signatures were derived from the tutorial-provided single-cell reference dataset distributed with cell2location (sc.h5ad) together with its pretrained RegressionModel. For gastric cancer samples, reference signatures were derived from a published human gastric cancer single-cell RNA-seq dataset (Sun et al., 2022).

The Visium Seurat object was converted to AnnData (.h5ad). Gene symbols from the spatial expression matrix were mapped to Ensembl gene IDs using biomaRt, and Ensembl IDs were added to feature annotations to ensure compatibility with the corresponding reference gene identifier system. The spatial AnnData object was loaded in Scanpy, gene identifiers were replaced with Ensembl IDs, and mitochondrial genes (prefix MT-) were flagged and excluded from the feature set used for cell2location fitting while remaining in the object for quality-control bookkeeping.

Reference signatures were extracted as the inferred mean expression per cell state/cell type (means_per_cluster_mu_fg), and both the spatial data and reference signatures were restricted to the intersecting gene set before model fitting. After running cell2location.models.Cell2location.setup_anndata, the model was trained on the full spatial dataset using batch_size=None, train_size=1, N_cells_per_location = 15, and detection_alpha = 20, for 20,000 epochs. Posterior cell abundance estimates were obtained using export_posterior(num_samples = 1000). Spot-level abundance summaries, including means_cell_abundance_w_sf, q05_cell_abundance_w_sf, and q95_cell_abundance_w_sf, were exported from adata_vis.obsm for downstream analyses. Unless otherwise specified, q05_cell_abundance_w_sf was used as the spot-level abundance estimate for each cell type.

**Additional Parameters for Monocle3 Pseudotime Analysis**

Pseudotime analysis was performed on LN2, which exhibited a well-defined GC architecture and a clear continuum across DZ, LZ, and Inter LZ-DZ regions. Raw spatial counts and spot-level metadata were extracted from the Seurat object, and a Monocle3 cell_data_set was constructed together with a gene_metadata table containing gene_short_name. Based on Seurat-derived regional annotations (NewClusters), analysis was restricted to Follicular DZ, Follicular LZ, and Inter LZ-DZ spots.

The subsetted cell_data_set was preprocessed using preprocess_cds(num_dim = 200), embedded with UMAP (reduce_dimension, reduction_method = "UMAP"), clustered using cluster_cells, and used to learn the principal trajectory graph with learn_graph. Spots were ordered using order_cells, and pseudotime values were written back to Seurat metadata and visualized in tissue space.

Trajectory-associated genes were identified using graph_test(neighbor_graph = "principal_graph", cores = 6), and genes with q < 0.05 were retained. Significant genes were grouped into co-expression modules using find_gene_modules(resolution = 0.007). Module-level expression was aggregated by NewClusters using aggregate_gene_expression and visualized as a module-by-cluster heatmap. The top 100 genes ranked by Moran’s I were visualized along pseudotime using plot_genes_in_pseudotime.

**Additional Parameters for Spatial HdWGCNA**

Spatial hdWGCNA was performed using hdWGCNA v0.4.05. Spots annotated as non-GC (GC_part = "Others") were removed before network construction. Genes were pre-selected using SetupForWGCNA(gene_select = "fraction", fraction = 0.05, wgcna_name = "vis"). To reduce spot-level noise, spatial spots were aggregated into metaspots by GC_Maturity_Label using MetaspotsByGroups, followed by NormalizeMetacells. The Spatial assay was then used as network input with SetDatExpr.

Candidate soft-thresholding powers were evaluated using TestSoftPowers, and a power compatible with approximate scale-free topology was selected. Networks were constructed using ConstructNetwork(tom_name = "test", overwrite_tom = TRUE), followed by ModuleEigengenes and ModuleConnectivity. Module eigengenes were added to Seurat metadata and visualized using DotPlot and SpatialFeaturePlot.

Module co-expression structure was summarized using RunModuleUMAP(n_hubs = 5, n_neighbors = 5, min_dist = 0.3, spread = 1). Hub genes were identified with GetHubGenes(n_hubs = 150) on the basis of module membership (kME). To avoid circularity in downstream interpretation, mitochondrial genes and GC maturity marker genes used for stage labeling were excluded before enrichment analyses.

GO enrichment was performed using clusterProfiler v4.12.0 and org.Hs.eg.db v3.19.1. For each module, filtered hub-gene lists were used as input to enrichGO(OrgDb = org.Hs.eg.db, keyType = "SYMBOL", ont = "ALL", pAdjustMethod = "BH", readable = TRUE). Significant GO terms were defined by p.adjust < 0.05. For visualization, a composite combined_score was calculated as GeneRatio × [−log10(p.adjust)].

**Additional Parameters for SHM-Stratified Analysis of Follicular DZ Spots**

Spots annotated as Follicular DZ in the Seurat object (NewClusters) were extracted. SHM was quantified using MiXCR-derived IGH VDJ_mutation_rate. For each spot, SHM was summarized as the mean VDJ_mutation_rate across all IGH entries assigned to that spot (avg_15G_VDJ_mutation_rate). Spots were ranked by this value and classified into three categories: High_Mutation (>0.05), Low_Mutation (0–0.05), and No_Mutation (0). These labels were stored in Seurat metadata as SHM_VDJ.

To identify SHM-associated transcriptional programs, differential feature screening across SHM_VDJ groups was performed using FindAllMarkers(test.use = "roc"), generating an AUC-like discriminative metric (myAUC) for each gene. For each gene, the record with the maximal myAUC across groups was retained, and mitochondrial genes were excluded. SHM-associated genes were visualized using AUC-ranked plots, and predefined gene sets were displayed across SHM groups using DotPlot.

The three SHM-associated gene sets were then treated as functional modules and scored per DZ spot using AddModuleScore (Score_High, Score_Low, Score_No, nbin = 2). Module scores were correlated with spot-level SHM using Spearman rank correlation.

**Additional Details for SpatialSNV and Candidate Somatic Variant Prioritization**

Spatial transcriptomic RNA data and spatial SNV matrices were integrated using Scanpy (v1.9.0). Spatial coordinates were retained to preserve tissue architecture. Variant allele frequency (Alt) and sequencing depth matrices were aligned to RNA-seq barcodes, and only shared spots were retained.

Variants were annotated using the SpatialSNV pipeline together with ANNOVAR and the refGene (hg38) database. Functional consequences, including genomic region and mutation type, were assigned to each variant. SNV matrices were normalized against total RNA counts using normalize_with_rna to account for technical variation in transcript capture across spots.

To prioritize candidate somatic variants, a multi-step filtering strategy was applied:
(i) only exonic, nonsynonymous SNVs resulting in amino acid substitutions were retained;
(ii) spatial clustering was evaluated using Moran’s I, and variants with Moran’s I > 0.1 were prioritized; and
(iii) tumor-to-TLS enrichment was used as an additional filter, retaining variants with a tumor-to-TLS enrichment ratio > 2.0 or variants detected exclusively within tumor-rich regions.

Candidate variants were further resolved at the transcript level by expanding ANNOVAR annotations into one-to-many mappings to specific Ensembl/RefSeq transcripts.

**Additional Details for Spatial Co-Localization and TCR–SNV Pairing**

SNV and TCR matrices were integrated into a unified Seurat object. To identify candidate TCR–SNV interactions, a UMI-based spatial pairing algorithm was applied to calculate UMI Precision, defined as the proportion of co-localized UMIs within shared capture spots. High-confidence pairs were retained using the following criteria: UMI Precision ≥ 0.5, spatial co-localization P < 0.05 based on permutation testing, and at least three co-localized UMIs.

To evaluate immune-context perturbation associated with specific TCR–mutation engagement, cell-type abundances per spot derived from cell2location were compared between interaction sites and background regions. For each validated pair, log-fold changes of specific immune cell lineages were calculated. TCR sequences were assigned unique identifiers based on CDR3 sequence identity to ensure consistency across mutation niches.

**Additional Details for Neoantigen Prediction Using DeepAntigen**

For each candidate variant identified by SpatialSNV, the corresponding protein sequence was retrieved from UniProt, and a 20-residue sequence window flanking the mutation site was extracted. Candidate neoepitopes of 9 and 10 amino acids were generated using a sliding-window approach, retaining only peptides containing the substituted residue. Candidate neoantigens and their spatially co-localized TCR clonotypes were exhaustively paired and evaluated using deepAntigen to generate interaction scores for all candidate peptide–TCR combinations.
