## Supplementary Figure for "SpaCir-VDJ: a broadly compatible circularization strategy for spatial immune repertoire profiling"

### Figure S1

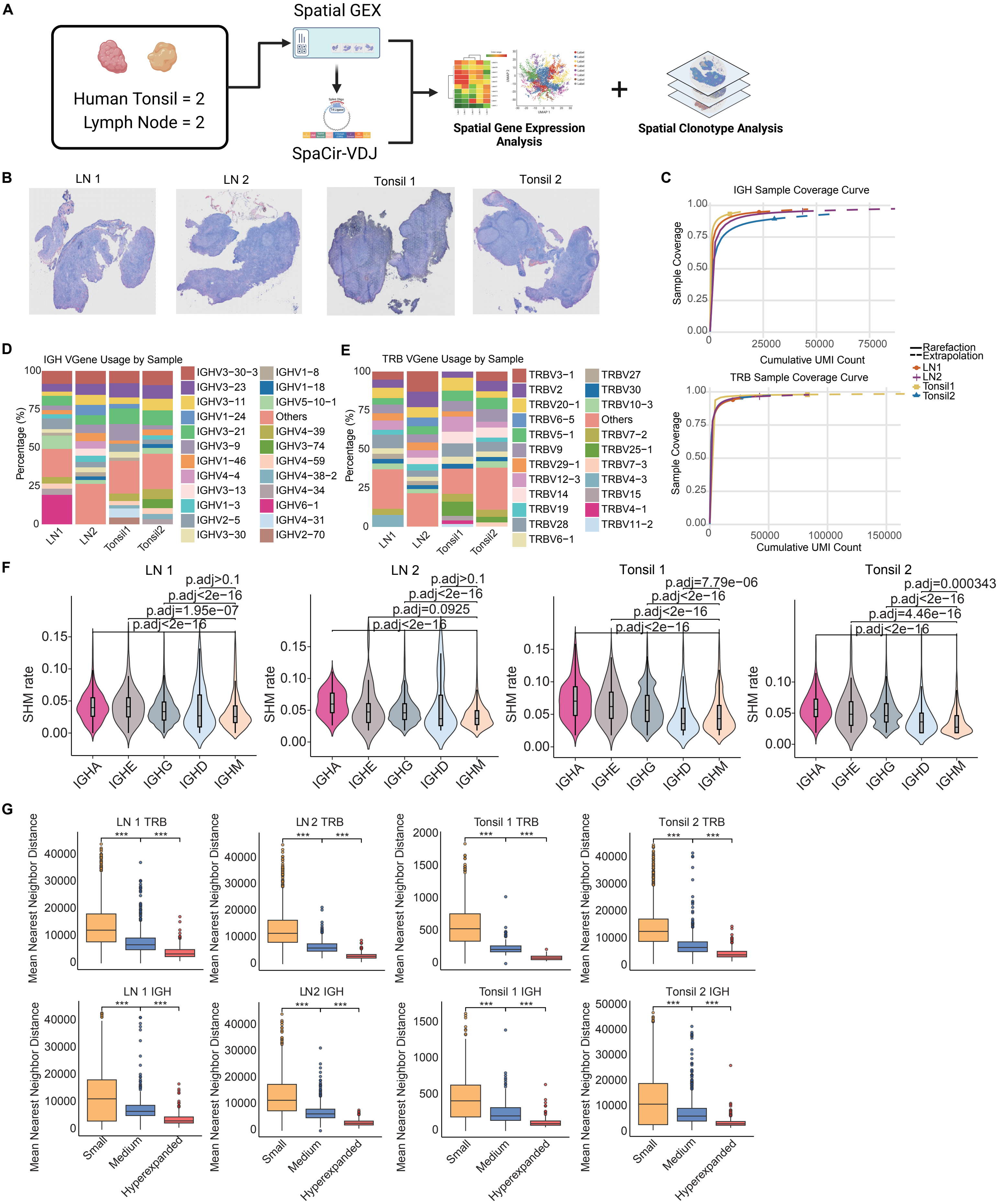
A
Spatial GEX
Human Tonsil = 2
Lymph Node = 2
SpaCir-VDJ
Spatial Gene Expression Analysis
Spatial Clonotype Analysis
B
LN 1
LN 2
Tonsil 1
Tonsil 2
C
IGH Sample Coverage Curve
Sample Coverage
Cumulative UMI Count
TRB Sample Coverage Curve
Rarefaction
Extrapolation
LN1
LN2
Tonsil1
Tonsil2
D
IGH VGene Usage by Sample
Percentage (%)
E
TRB VGene Usage by Sample
F
LN 1
LN 2
Tonsil 1
Tonsil 2
SHM rate
IGHA
IGHE
IGHG
IGHD
IGHM
G
LN 1 TRB
LN 2 TRB
Tonsil 1 TRB
Tonsil 2 TRB
LN 1 IGH
LN2 IGH
Tonsil 1 IGH
Tonsil 2 IGH
Mean Nearest Neighbor Distance
Small
Medium
Hyperexpanded

#### Figure S2

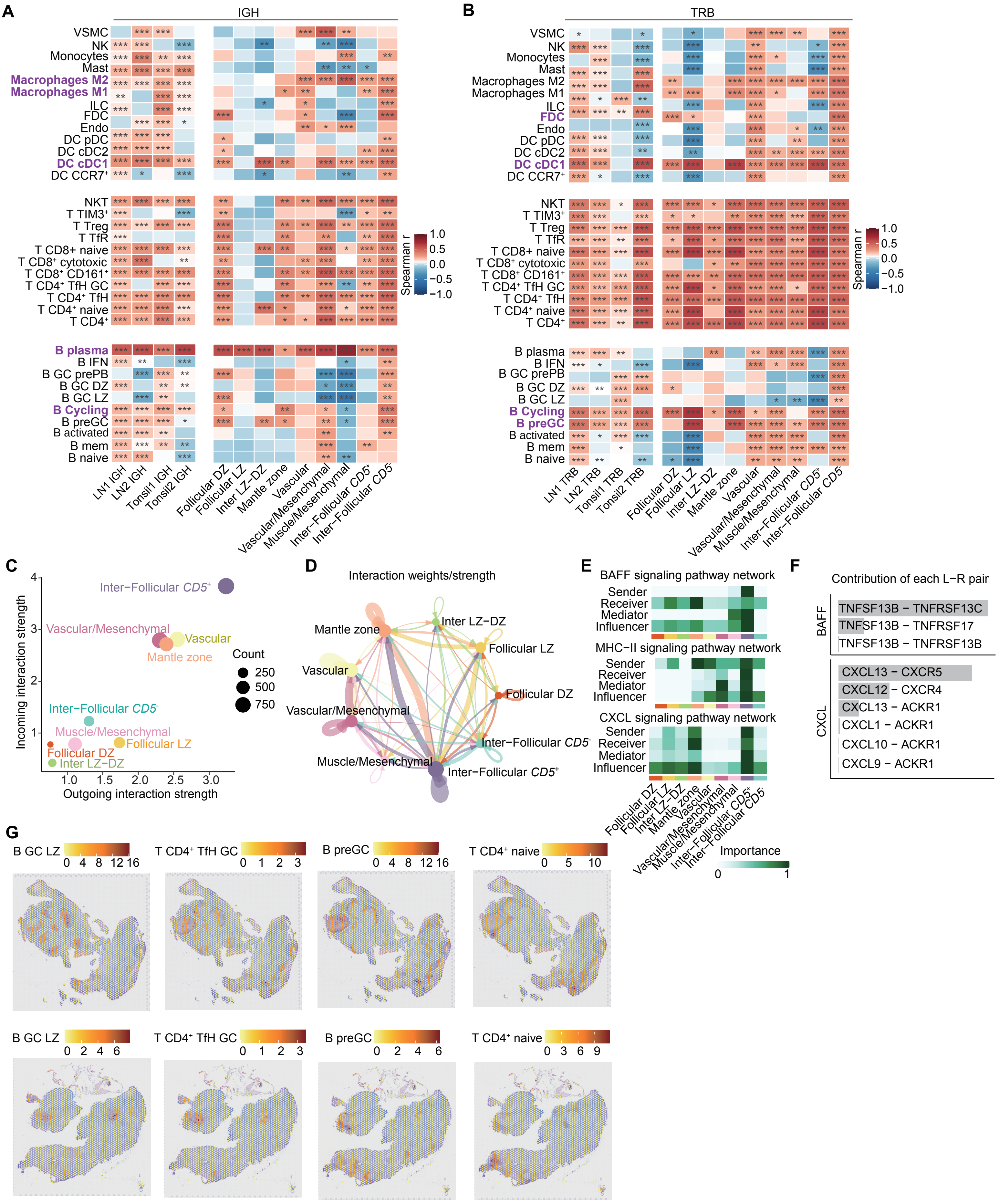

A
IGH
B
TRB
Spearman r
C
Incoming interaction strength
Outgoing interaction strength
Count
D
Interaction weights/strength
E
BAFF signaling pathway network
MHC-II signaling pathway network
CXCL signaling pathway network
Sender
Receiver
Mediator
Influencer
Importance
F
Contribution of each L-R pair
BAFF
TNFSF13B − TNFRSF13C
TNFSF13B − TNFRSF17
TNFSF13B − TNFRSF13B
CXCL
CXCL13 − CXCR5
CXCL12 − CXCR4
CXCL13 − ACKR1
CXCL1 − ACKR1
CXCL10 − ACKR1
CXCL9 − ACKR1
G
B GC LZ
T CD4⁺ TfH GC
B preGC
T CD4⁺ naive

**Figure S3**

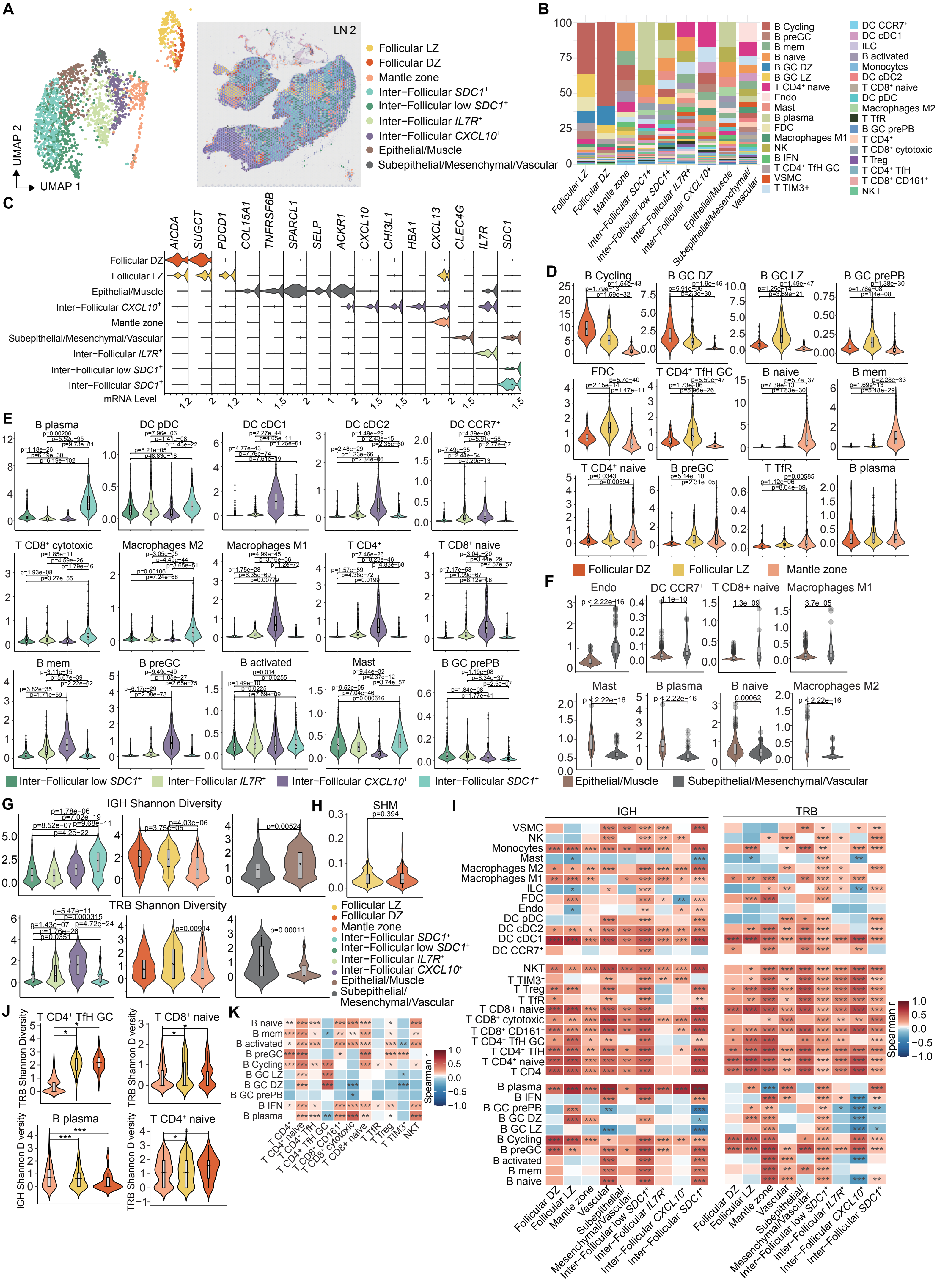
A
LN 2
UMAP 1
UMAP 2
Follicular LZ
Follicular DZ
Mantle zone
Inter−Follicular SDC1+
Inter−Follicular low SDC1+
Inter−Follicular IL7R+
Inter−Follicular CXCL10+
Epithelial/Muscle
Subepithelial/Mesenchymal/Vascular
B
B Cycling
B preGC
B mem
B naive
B GC DZ
B GC LZ
T CD4+ naive
Endo
Mast
B plasma
FDC
Macrophages M1
NK
B IFN
T CD4+ TfH GC
VSMC
T TIM3+
DC CCR7+
DC cDC1
ILC
B activated
Monocytes
DC cDC2
T CD8+ naive
DC pDC
Macrophages M2
T TfR
B GC prePB
T CD4+
T CD8+ cytotoxic
T Treg
T CD4+ TfH
T CD8+ CD161+
NKT
C
AICDA
SUGCT
PDCD1
COL15A1
TNFRSF6B
SPARCL1
SELP
ACKR1
CXCL10
CHI3L1
HBA1
CXCL13
CLEC4G
IL7R
SDC1
mRNA Level
D
B Cycling
B GC DZ
B GC LZ
B GC prePB
FDC
T CD4+ TfH GC
B naive
B mem
T CD4+ naive
B preGC
T TfR
B plasma
Follicular DZ
Follicular LZ
Mantle zone
E
B plasma
DC pDC
DC cDC1
DC cDC2
DC CCR7+
T CD8+ cytotoxic
Macrophages M2
Macrophages M1
T CD4+
T CD8+ naive
B mem
B preGC
B activated
Mast
B GC prePB
Inter−Follicular low SDC1+
Inter−Follicular IL7R+
Inter−Follicular CXCL10+
Inter−Follicular SDC1+
F
Endo
DC CCR7+
T CD8+ naive
Macrophages M1
Mast
B plasma
B naive
Macrophages M2
Epithelial/Muscle
Subepithelial/Mesenchymal/Vascular
G
IGH Shannon Diversity
TRB Shannon Diversity
H
SHM
p=0.394
I
IGH
TRB
Spearman r
J
T CD4+ TfH GC
T CD8+ naive
B plasma
T CD4+ naive
TRB Shannon Diversity
IGH Shannon Diversity
K
Spearman r

#### Figure S4

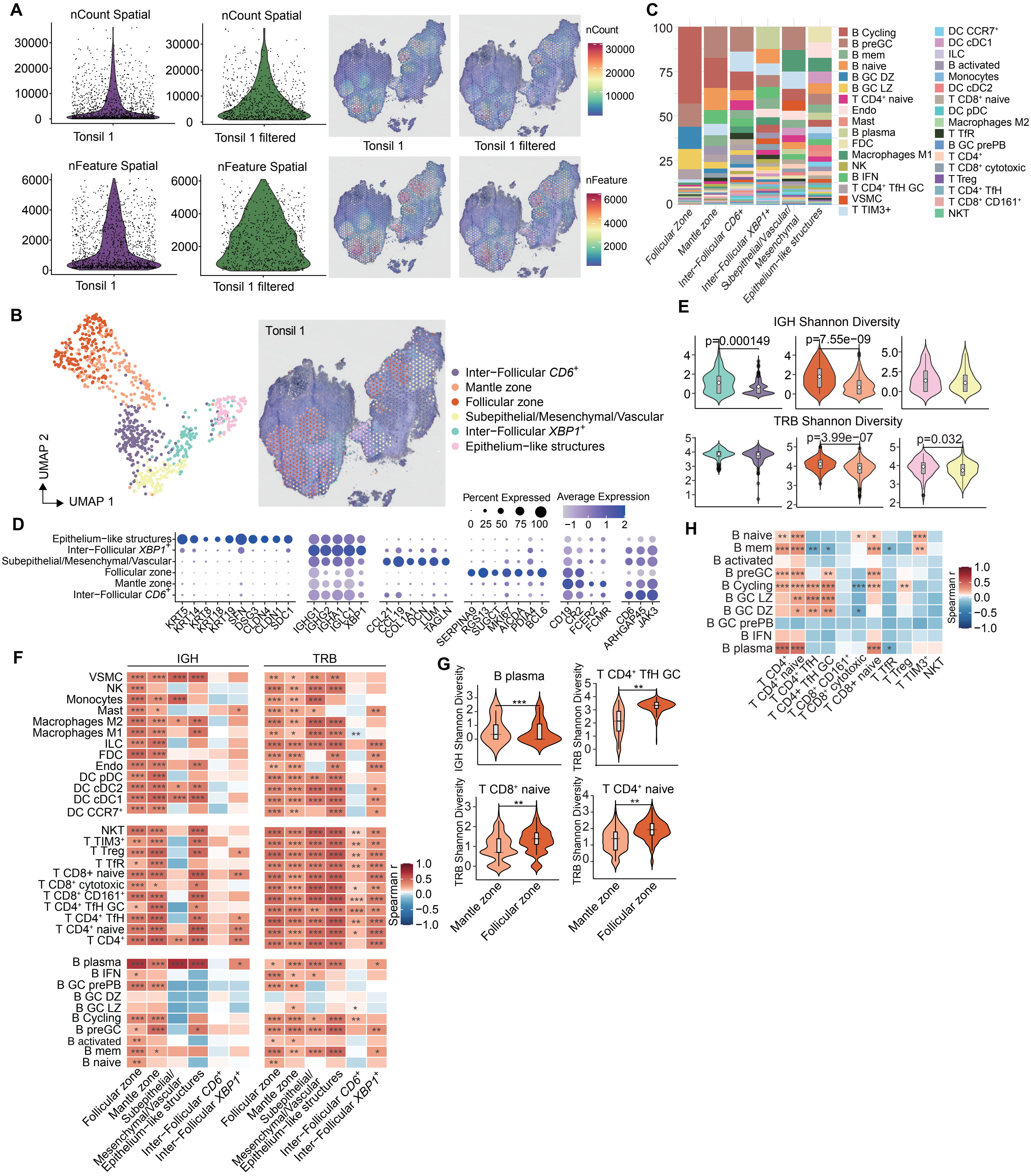
A
nCount Spatial
nCount Spatial
Tonsil 1
Tonsil 1 filtered
nFeature Spatial
nFeature Spatial
Tonsil 1
Tonsil 1 filtered
nCount
nFeature
B
UMAP 1
UMAP 2
Tonsil 1
Inter−Follicular CD6+
Mantle zone
Follicular zone
Subepithelial/Mesenchymal/Vascular
Inter−Follicular XBP1+
Epithelium−like structures
C
Follicular Zone
Mantle zone
Inter−Follicular CD6+
Inter−Follicular XBP1+
Subepithelial/Vascular/Mesenchymal
Epithelium−like structures
B Cycling
B preGC
B mem
B naive
B GC DZ
B GC LZ
T CD4+ naive
Endo
Mast
B plasma
FDC
Macrophages M1
NK
B IFN
T CD4+ TfH GC
VSMC
T TIM3+
DC CCR7+
DC cDC1
ILC
B activated
Monocytes
DC cDC2
T CD8+ naive
DC pDC
Macrophages M2
T TfR
B GC prePB
T CD4+
T CD8+ cytotoxic
T Treg
T CD4+ TfH
T CD8+ CD161+
NKT
D
Percent Expressed
Average Expression
KRT5
KRT14
KRT8
KRT18
KRT19
SFN
DSG3
CLDN4
CLDN1
SDC1
IGHG1
IGHG2
IGHA1
IGLC1
XBP1
CCL21
CCL19
COL1A1
DCN
LUM
TAGLN
SERPINA9
RGS13
SUGCT
MKI67
AICDA
PDIA4
BCL6
CD19
CR2
FCER2
FCMR
CD6
ARHGAP45
JAK3
E
IGH Shannon Diversity
p=0.000149
p=7.55e−09
TRB Shannon Diversity
p=3.99e−07
p=0.032
F
IGH
TRB
Spearman r
G
B plasma
T CD4+ TfH GC
T CD8+ naive
T CD4+ naive
IGH Shannon Diversity
TRB Shannon Diversity
Mantle zone
Follicular zone
H
Spearman r

#### Figure S5

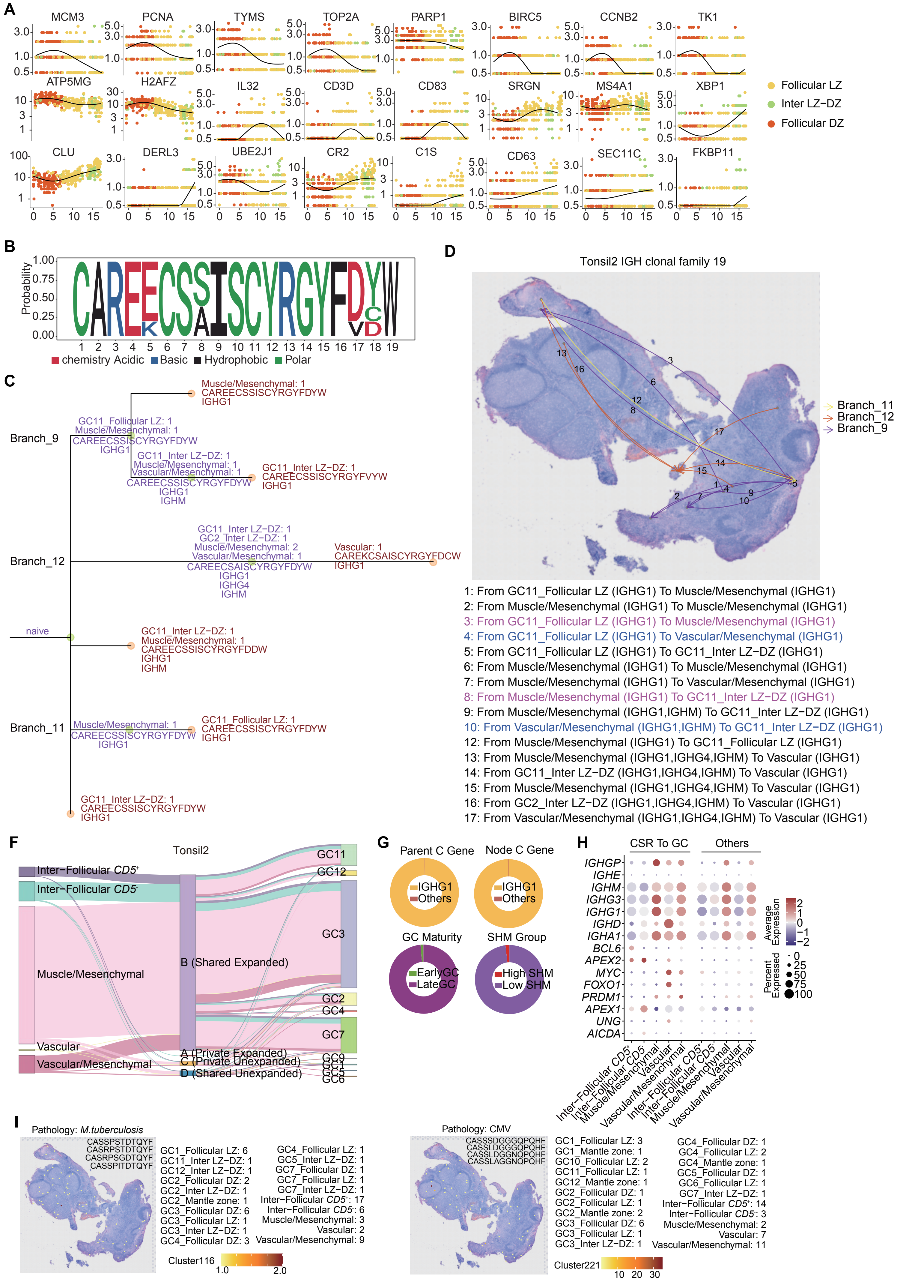
A
MCM3
PCNA
TYMS
TOP2A
PARP1
BIRC5
CCNB2
TK1
ATP5MG
H2AFZ
IL32
CD3D
CD83
SRGN
MS4A1
XBP1
CLU
DERL3
UBE2J1
CR2
C1S
CD63
SEC11C
FKBP11
Follicular LZ
Inter LZ−DZ
Follicular DZ
B
Probability
chemistry Acidic Basic Hydrophobic Polar
C
Branch_9
Branch_12
Branch_11
naive
Muscle/Mesenchymal: 1
CAREECSSISCYRGYFDYW
IGHG1
GC11_Follicular LZ: 1
Muscle/Mesenchymal: 1
CAREECSSISCYRGYFDYW
IGHG1
GC11_Inter LZ−DZ: 1
Muscle/Mesenchymal: 1
Vascular/Mesenchymal: 1
CAREECSSISCYRGYFDYW
IGHG1
IGHM
GC11_Inter LZ−DZ: 1
CAREECSSISCYRGYFVYW
IGHG1
GC11_Inter LZ−DZ: 1
GC2_Inter LZ−DZ: 1
Muscle/Mesenchymal: 2
Vascular/Mesenchymal: 1
CAREECSAISCYRGYFDYW
IGHG1
IGHG4
IGHM
Vascular: 1
CAREKCSAISCYRGYFDCW
IGHG1
GC11_Inter LZ−DZ: 1
Muscle/Mesenchymal: 1
CAREECSSISCYRGYFDDW
IGHG1
IGHM
Muscle/Mesenchymal: 1
CAREECSSISCYRGYFDYW
IGHG1
GC11_Follicular LZ: 1
CAREECSSISCYRGYFDYW
IGHG1
GC11_Inter LZ−DZ: 1
CAREECSSISCYRGYFDYW
IGHG1
D
Tonsil2 IGH clonal family 19
Branch_11
Branch_12
Branch_9
1: From GC11_Follicular LZ (IGHG1) To Muscle/Mesenchymal (IGHG1)
2: From Muscle/Mesenchymal (IGHG1) To Muscle/Mesenchymal (IGHG1)
3: From GC11_Follicular LZ (IGHG1) To Muscle/Mesenchymal (IGHG1)
4: From GC11_Follicular LZ (IGHG1) To Vascular/Mesenchymal (IGHG1)
5: From GC11_Follicular LZ (IGHG1) To GC11_Inter LZ−DZ (IGHG1)
6: From Muscle/Mesenchymal (IGHG1) To Muscle/Mesenchymal (IGHG1)
7: From Muscle/Mesenchymal (IGHG1) To Vascular/Mesenchymal (IGHG1)
8: From Muscle/Mesenchymal (IGHG1) To GC11_Inter LZ−DZ (IGHG1)
9: From Muscle/Mesenchymal (IGHG1,IGHM) To GC11_Inter LZ−DZ (IGHG1)
10: From Vascular/Mesenchymal (IGHG1,IGHM) To GC11_Inter LZ−DZ (IGHG1)
12: From Muscle/Mesenchymal (IGHG1) To GC11_Follicular LZ (IGHG1)
13: From Muscle/Mesenchymal (IGHG1,IGHG4,IGHM) To Vascular (IGHG1)
14: From GC11_Inter LZ−DZ (IGHG1,IGHG4,IGHM) To Vascular (IGHG1)
15: From Muscle/Mesenchymal (IGHG1,IGHG4,IGHM) To Vascular (IGHG1)
16: From GC2_Inter LZ−DZ (IGHG1,IGHG4,IGHM) To Vascular (IGHG1)
17: From Vascular/Mesenchymal (IGHG1,IGHG4,IGHM) To Vascular (IGHG1)
F
Tonsil2
Inter−Follicular CD5+
Inter−Follicular CD5-
Muscle/Mesenchymal
Vascular
Vascular/Mesenchymal
B (Shared Expanded)
A (Private Expanded)
C (Private Unexpanded)
D (Shared Unexpanded)
GC11
GC12
GC3
GC2
GC4
GC7
GC9
GC1
GC5
GC6
G
Parent C Gene
Node C Gene
IGHG1
Others
GC Maturity
SHM Group
EarlyGC
LateGC
High SHM
Low SHM
H
CSR To GC
Others
IGHGP
IGHE
IGHM
IGHG3
IGHG1
IGHD
IGHA1
BCL6
APEX2
MYC
FOXO1
PRDM1
APEX1
UNG
AICDA
Average Expression
Percent Expressed
Inter−Follicular CD5+
Inter−Follicular CD5-
Muscle/Mesenchymal
Vascular
Vascular/Mesenchymal
I
Pathology: M.tuberculosis
CASSPSTDTQYF
CASRPSTDTQYF
CASRPSGDTQYF
CASSPITDTQYF
GC1_Follicular LZ: 6
GC11_Inter LZ−DZ: 1
GC12_Inter LZ−DZ: 1
GC2_Follicular DZ: 2
GC2_Inter LZ−DZ: 1
GC2_Mantle zone: 1
GC3_Follicular DZ: 6
GC3_Follicular LZ: 1
GC3_Inter LZ−DZ: 1
GC4_Follicular DZ: 3
GC4_Follicular LZ: 1
GC5_Inter LZ−DZ: 1
GC7_Follicular DZ: 1
GC7_Follicular LZ: 1
GC7_Inter LZ−DZ: 1
Inter−Follicular CD5+: 17
Inter−Follicular CD5-: 6
Muscle/Mesenchymal: 3
Vascular: 2
Vascular/Mesenchymal: 9
Cluster116
Pathology: CMV
CASSSDGGGQPQHF
CASSLDGGGQPQHF
CASSLDGGNQPQHF
CASSLAGGNQPQHF
GC1_Follicular LZ: 3
GC1_Mantle zone: 1
GC10_Follicular LZ: 2
GC11_Follicular LZ: 1
GC12_Mantle zone: 1
GC2_Follicular DZ: 1
GC2_Follicular LZ: 1
GC2_Mantle zone: 2
GC3_Follicular DZ: 6
GC3_Follicular LZ: 1
GC3_Inter LZ−DZ: 1
GC4_Follicular DZ: 1
GC4_Follicular LZ: 2
GC4_Mantle zone: 1
GC5_Follicular DZ: 1
GC6_Follicular LZ: 1
GC7_Inter LZ−DZ: 1
Inter−Follicular CD5+: 14
Inter−Follicular CD5-: 3
Muscle/Mesenchymal: 2
Vascular: 7
Vascular/Mesenchymal: 11
Cluster221

### Figure S6

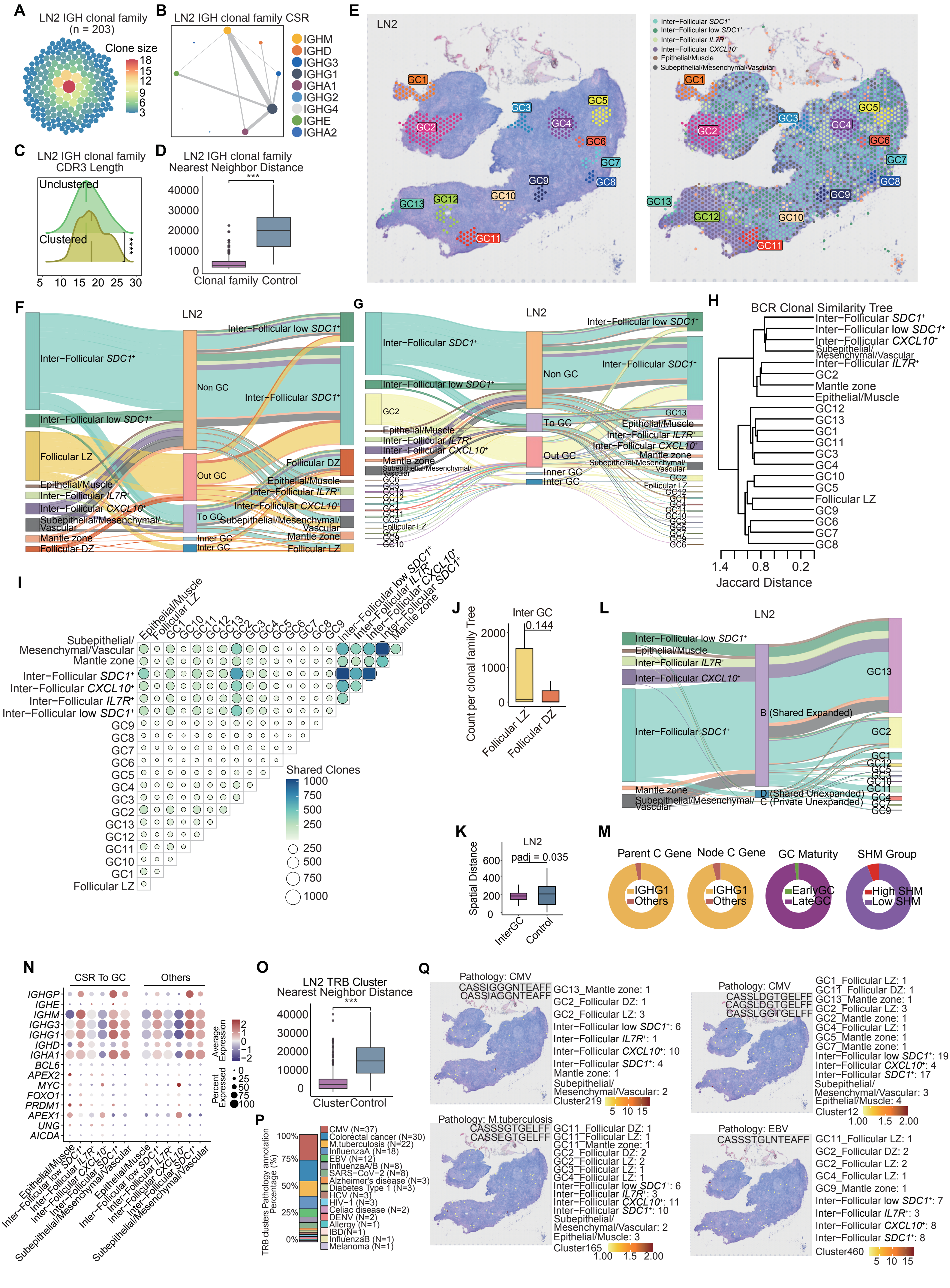
A
LN2 IGH clonal family
(n = 203)
Clone size
18
15
12
9
6
3
B
LN2 IGH clonal family CSR
IGHM
IGHD
IGHG3
IGHG1
IGHA1
IGHG2
IGHG4
IGHE
IGHA2
C
LN2 IGH clonal family
CDR3 Length
Unclustered
Clustered
****
5 10 15 20 25 30
D
LN2 IGH clonal family
Nearest Neighbor Distance
***
40000
30000
20000
10000
0
Clonal family Control
E
LN2
Inter−Follicular SDC1+
Inter−Follicular low SDC1+
Inter−Follicular IL7R+
Inter−Follicular CXCL10+
Epithelial/Muscle
Subepithelial/Mesenchymal/Vascular
GC1
GC2
GC3
GC4
GC5
GC6
GC7
GC8
GC9
GC10
GC11
GC12
GC13
F
LN2
Inter−Follicular SDC1+
Inter−Follicular low SDC1+
Follicular LZ
Epithelial/Muscle
Inter−Follicular IL7R+
Inter−Follicular CXCL10+
Subepithelial/Mesenchymal/Vascular
Mantle zone
Follicular DZ
Non GC
Out GC
To GC
Inner GC
Inter GC
G
LN2
H
BCR Clonal Similarity Tree
Inter−Follicular SDC1+
Inter−Follicular low SDC1+
Inter−Follicular CXCL10+
Subepithelial/Mesenchymal/Vascular
Inter−Follicular IL7R+
GC2
Mantle zone
Epithelial/Muscle
GC12
GC13
GC1
GC11
GC3
GC4
GC10
GC5
Follicular LZ
GC9
GC6
GC7
GC8
1.4 0.8 0.2
Jaccard Distance
I
Shared Clones
1000
750
500
250
J
Inter GC
0.144
Count per clonal family Tree
2000
1000
0
Follicular LZ
Follicular DZ
K
LN2
padj = 0.035
Spatial Distance
600
400
200
0
InterGC
Control
L
LN2
B (Shared Expanded)
D (Shared Unexpanded)
C (Private Unexpanded)
M
Parent C Gene
Node C Gene
GC Maturity
SHM Group
IGHG1
Others
EarlyGC
LateGC
High SHM
Low SHM
N
CSR To GC
Others
IGHGP
IGHE
IGHM
IGHG3
IGHG1
IGHD
IGHA1
BCL6
APEX2
MYC
FOXO1
PRDM1
APEX1
UNG
AICDA
Average Expression
2
1
0
−1
−2
Percent Expressed
0
25
50
75
100
O
LN2 TRB Cluster
Nearest Neighbor Distance
***
40000
30000
20000
10000
0
Cluster Control
P
TRB clusters Pathology annotation Percentage (%)
100%
75%
50%
25%
0%
CMV (N=37)
Colorectal cancer (N=30)
M.tuberculosis (N=22)
InfluenzaA (N=18)
EBV (N=12)
InfluenzaA/B (N=8)
SARS−CoV−2 (N=8)
Alzheimer's disease (N=3)
Diabetes Type 1 (N=3)
HCV (N=3)
HIV−1 (N=3)
Celiac disease (N=2)
DENV (N=2)
Allergy (N=1)
IBD (N=1)
InfluenzaB (N=1)
Melanoma (N=1)
Q
Pathology: CMV
CASSIGGGNTEAFF
CASSIAGGNTEAFF
GC13_Mantle zone: 1
GC2_Follicular DZ: 1
GC2_Follicular LZ: 3
Inter−Follicular low SDC1+: 6
Inter−Follicular IL7R+: 1
Inter−Follicular CXCL10+: 10
Inter−Follicular SDC1+: 4
Mantle zone: 1
Subepithelial/Mesenchymal/Vascular: 2
Cluster219
Pathology: CMV
CASSLDGTGELFF
CAGSLDGTGELFF
CASSLGGTGELFF
GC1_Follicular LZ: 1
GC11_Follicular DZ: 1
GC13_Mantle zone: 1
GC2_Follicular LZ: 3
GC2_Mantle zone: 1
GC4_Follicular LZ: 1
GC5_Mantle zone: 1
GC7_Mantle zone: 1
Inter−Follicular low SDC1+: 19
Inter−Follicular CXCL10+: 4
Inter−Follicular SDC1+: 17
Subepithelial/Mesenchymal/Vascular: 3
Epithelial/Muscle: 4
Cluster12
Pathology: M.tuberculosis
CASSGTGELFF
CASSEGTGELFF
GC11_Follicular DZ: 1
GC11_Follicular LZ: 1
GC11_Mantle zone: 1
GC2_Follicular DZ: 2
GC2_Follicular LZ: 2
GC3_Follicular LZ: 1
GC4_Follicular LZ: 1
Inter−Follicular low SDC1+: 6
Inter−Follicular IL7R+: 3
Inter−Follicular CXCL10+: 11
Inter−Follicular SDC1+: 10
Subepithelial/Mesenchymal/Vascular: 2
Epithelial/Muscle: 3
Cluster165
Pathology: EBV
CASSSTGLNTEAFF
GC11_Follicular LZ: 1
GC2_Follicular DZ: 2
GC2_Follicular LZ: 2
GC4_Follicular LZ: 1
GC9_Mantle zone: 1
Inter−Follicular low SDC1+: 7
Inter−Follicular IL7R+: 3
Inter−Follicular CXCL10+: 8
Inter−Follicular SDC1+: 8
Cluster460

#### Figure S7

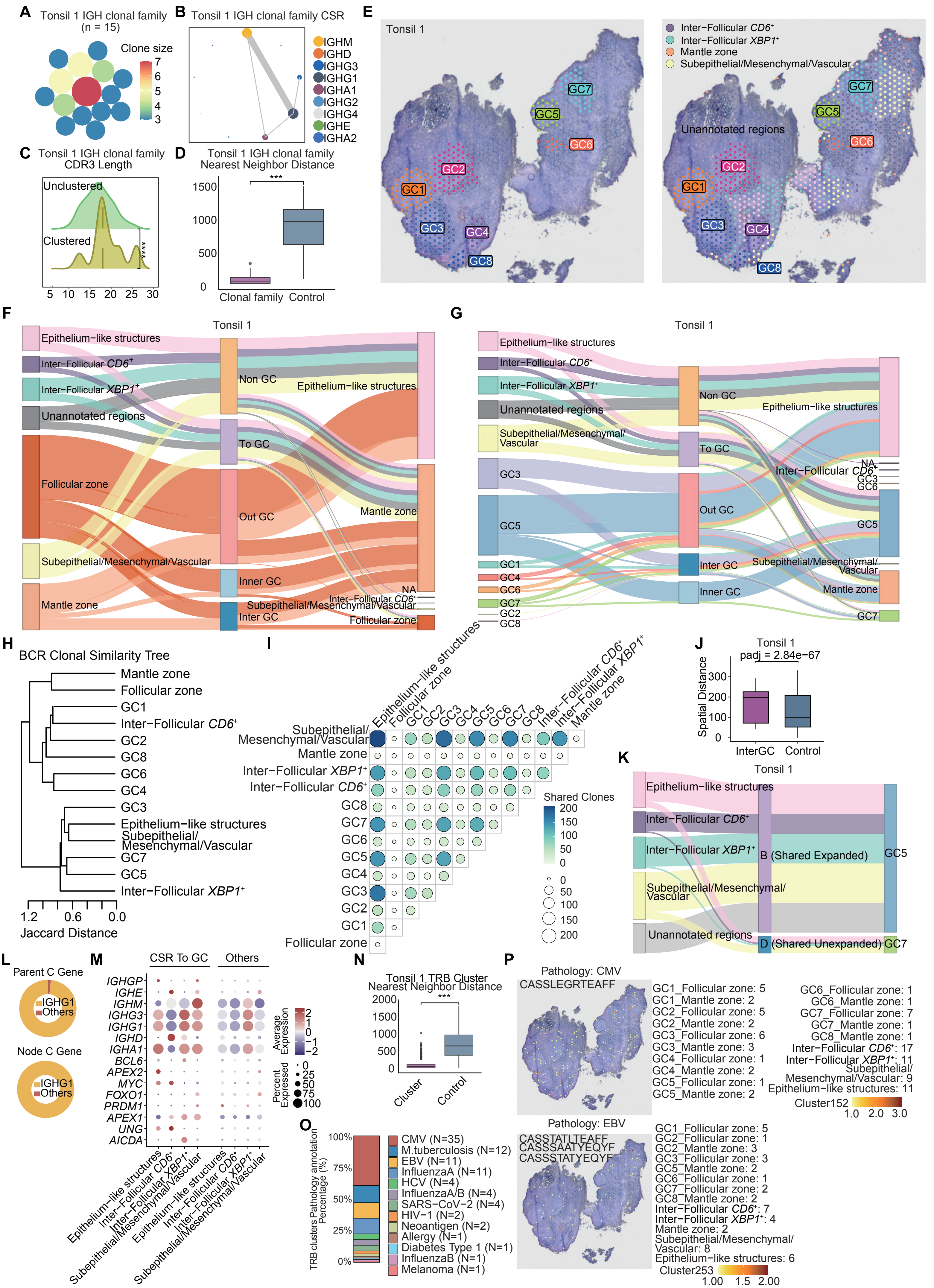

A Tonsil 1 IGH clonal family (n = 15)
Clone size
B Tonsil 1 IGH clonal family CSR
IGHM
IGHD
IGHG3
IGHG1
IGHA1
IGHG2
IGHG4
IGHE
IGHA2
C Tonsil 1 IGH clonal family CDR3 Length
Unclustered
Clustered
D Tonsil 1 IGH clonal family Nearest Neighbor Distance
Clonal family
Control
E Tonsil 1
Inter−Follicular CD6+
Inter−Follicular XBP1+
Mantle zone
Subepithelial/Mesenchymal/Vascular
Unannotated regions
F Tonsil 1
Epithelium−like structures
Unannotated regions
Follicular zone
Mantle zone
Non GC
To GC
Out GC
Inner GC
Inter GC
G Tonsil 1
H BCR Clonal Similarity Tree
Jaccard Distance
I Shared Clones
J Tonsil 1
padj = 2.84e−67
Spatial Distance
InterGC
Control
K Tonsil 1
B (Shared Expanded)
D (Shared Unexpanded)
L Parent C Gene
Node C Gene
IGHG1
Others
M CSR To GC
Others
Average Expression
Percent Expressed
N Tonsil 1 TRB Cluster Nearest Neighbor Distance
Cluster
Control
O TRB clusters Pathology annotation Percentage (%)
CMV (N=35)
M.tuberculosis (N=12)
EBV (N=11)
InfluenzaA (N=11)
HCV (N=4)
InfluenzaA/B (N=4)
SARS−CoV−2 (N=4)
HIV−1 (N=2)
Neoantigen (N=2)
Allergy (N=1)
Diabetes Type 1 (N=1)
InfluenzaB (N=1)
Melanoma (N=1)
P Pathology: CMV
CASSLEGRTEAFF
GC1_Follicular zone: 5
GC1_Mantle zone: 2
GC2_Follicular zone: 5
GC2_Mantle zone: 2
GC3_Follicular zone: 6
GC3_Mantle zone: 3
GC4_Follicular zone: 1
GC4_Mantle zone: 2
GC5_Follicular zone: 1
GC5_Mantle zone: 2
GC6_Follicular zone: 1
GC6_Mantle zone: 1
GC7_Follicular zone: 7
GC7_Mantle zone: 1
GC8_Mantle zone: 1
Inter−Follicular CD6+: 17
Inter−Follicular XBP1+: 11
Subepithelial/Mesenchymal/Vascular: 9
Epithelium−like structures: 11
Cluster152
Pathology: EBV
CASSTATLTEAFF
CASSSAATYEQYF
CASSSTATYEQYF
GC1_Follicular zone: 5
GC2_Follicular zone: 1
GC2_Mantle zone: 3
GC3_Follicular zone: 3
GC5_Mantle zone: 2
GC6_Follicular zone: 1
GC7_Follicular zone: 2
GC8_Mantle zone: 2
Inter−Follicular CD6+: 7
Inter−Follicular XBP1+: 4
Mantle zone: 2
Subepithelial/Mesenchymal/Vascular: 8
Epithelium−like structures: 6
Cluster253

#### Figure S8

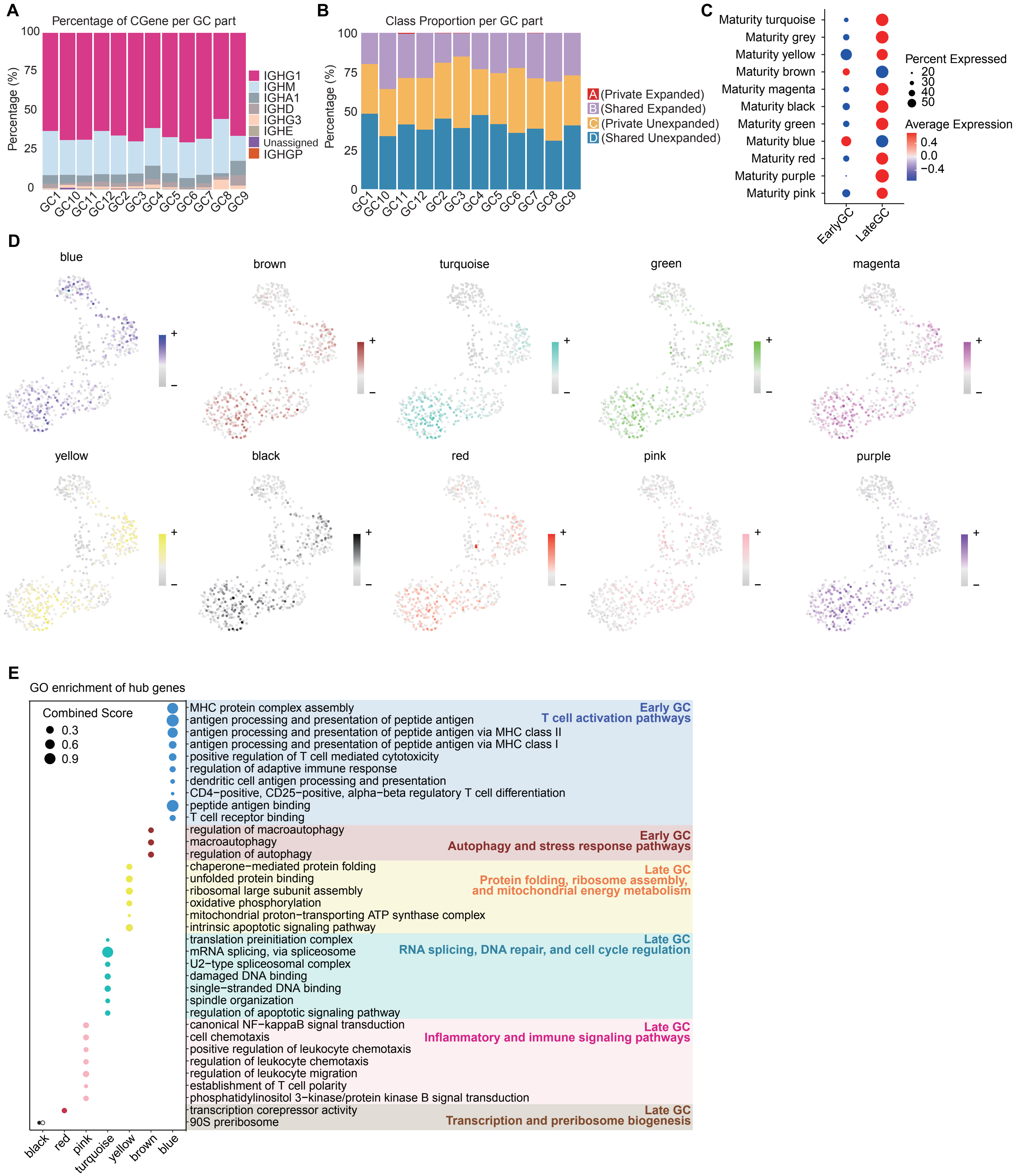

A
Percentage of CGene per GC part
Percentage (%)
IGHG1
IGHM
IGHA1
IGHD
IGHG3
IGHE
Unassigned
IGHGP
B
Class Proportion per GC part
Percentage (%)
A (Private Expanded)
B (Shared Expanded)
C (Private Unexpanded)
D (Shared Unexpanded)
C
Maturity turquoise
Maturity grey
Maturity yellow
Maturity brown
Maturity magenta
Maturity black
Maturity green
Maturity blue
Maturity red
Maturity purple
Maturity pink
Percent Expressed
Average Expression
EarlyGC
LateGC
D
blue
brown
turquoise
green
magenta
yellow
black
red
pink
purple
E
GO enrichment of hub genes
Combined Score
MHC protein complex assembly
antigen processing and presentation of peptide antigen
antigen processing and presentation of peptide antigen via MHC class II
antigen processing and presentation of peptide antigen via MHC class I
positive regulation of T cell mediated cytotoxicity
regulation of adaptive immune response
dendritic cell antigen processing and presentation
CD4−positive, CD25−positive, alpha−beta regulatory T cell differentiation
peptide antigen binding
T cell receptor binding
Early GC
T cell activation pathways
regulation of macroautophagy
macroautophagy
regulation of autophagy
Early GC
Autophagy and stress response pathways
chaperone−mediated protein folding
unfolded protein binding
ribosomal large subunit assembly
oxidative phosphorylation
mitochondrial proton−transporting ATP synthase complex
intrinsic apoptotic signaling pathway
Late GC
Protein folding, ribosome assembly, and mitochondrial energy metabolism
translation preinitiation complex
mRNA splicing, via spliceosome
U2−type spliceosomal complex
damaged DNA binding
single−stranded DNA binding
spindle organization
regulation of apoptotic signaling pathway
Late GC
RNA splicing, DNA repair, and cell cycle regulation
canonical NF−kappaB signal transduction
cell chemotaxis
positive regulation of leukocyte chemotaxis
regulation of leukocyte chemotaxis
regulation of leukocyte migration
establishment of T cell polarity
phosphatidylinositol 3−kinase/protein kinase B signal transduction
Late GC
Inflammatory and immune signaling pathways
transcription corepressor activity
90S preribosome
Late GC
Transcription and preribosome biogenesis
black
red
pink
turquoise
yellow
brown
blue

**Figure S9**

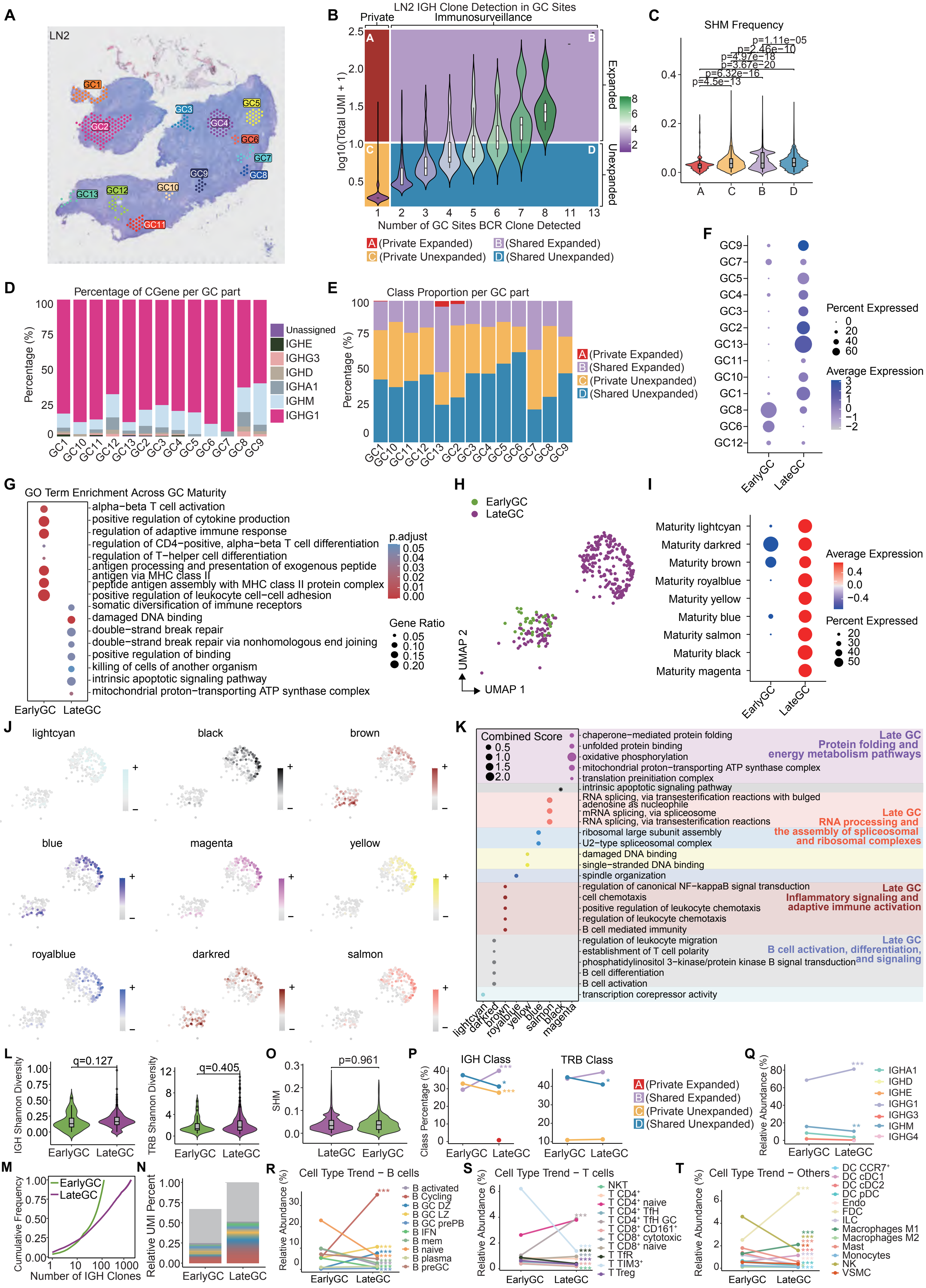
A
LN2
GC1
GC2
GC3
GC4
GC5
GC6
GC7
GC8
GC9
GC10
GC11
GC12
GC13
B
LN2 IGH Clone Detection in GC Sites
Private
Immunosurveillance
log10(Total UMI + 1)
Number of GC Sites BCR Clone Detected
Expanded
Unexpanded
A (Private Expanded)
B (Shared Expanded)
C (Private Unexpanded)
D (Shared Unexpanded)
C
SHM Frequency
p=1.11e−05
p=2.46e−10
p=4.97e−18
p=3.67e−20
p=6.32e−16
p=4.5e−13
D
Percentage of CGene per GC part
Percentage (%)
Unassigned
IGHE
IGHG3
IGHD
IGHA1
IGHM
IGHG1
E
Class Proportion per GC part
A (Private Expanded)
B (Shared Expanded)
C (Private Unexpanded)
D (Shared Unexpanded)
F
Percent Expressed
Average Expression
EarlyGC
LateGC
G
GO Term Enrichment Across GC Maturity
alpha−beta T cell activation
positive regulation of cytokine production
regulation of adaptive immune response
regulation of CD4−positive, alpha−beta T cell differentiation
regulation of T−helper cell differentiation
antigen processing and presentation of exogenous peptide antigen via MHC class II
peptide antigen assembly with MHC class II protein complex
positive regulation of leukocyte cell−cell adhesion
somatic diversification of immune receptors
damaged DNA binding
double−strand break repair
double−strand break repair via nonhomologous end joining
positive regulation of binding
killing of cells of another organism
intrinsic apoptotic signaling pathway
mitochondrial proton−transporting ATP synthase complex
p.adjust
Gene Ratio
H
EarlyGC
LateGC
UMAP 1
UMAP 2
I
Maturity lightcyan
Maturity darkred
Maturity brown
Maturity royalblue
Maturity yellow
Maturity blue
Maturity salmon
Maturity black
Maturity magenta
Average Expression
Percent Expressed
J
lightcyan
black
brown
blue
magenta
yellow
royalblue
darkred
salmon
K
Combined Score
chaperone−mediated protein folding
unfolded protein binding
oxidative phosphorylation
mitochondrial proton−transporting ATP synthase complex
translation preinitiation complex
intrinsic apoptotic signaling pathway
RNA splicing, via transesterification reactions with bulged adenosine as nucleophile
mRNA splicing, via spliceosome
RNA splicing, via transesterification reactions
ribosomal large subunit assembly
U2−type spliceosomal complex
damaged DNA binding
single−stranded DNA binding
spindle organization
regulation of canonical NF−kappaB signal transduction
cell chemotaxis
positive regulation of leukocyte chemotaxis
regulation of leukocyte chemotaxis
B cell mediated immunity
regulation of leukocyte migration
establishment of T cell polarity
phosphatidylinositol 3−kinase/protein kinase B signal transduction
B cell differentiation
B cell activation
transcription corepressor activity
Late GC Protein folding and energy metabolism pathways
Late GC RNA processing and the assembly of spliceosomal and ribosomal complexes
Late GC Inflammatory signaling and adaptive immune activation
Late GC B cell activation, differentiation, and signaling
L
IGH Shannon Diversity
q=0.127
TRB Shannon Diversity
q=0.405
O
SHM
p=0.961
P
IGH Class
TRB Class
Class Percentage (%)
Q
Relative Abundance (%)
IGHA1
IGHD
IGHE
IGHG1
IGHG3
IGHM
IGHG4
M
Cumulative Frequency
Number of IGH Clones
N
Relative UMI Percent
R
Cell Type Trend − B cells
B activated
B Cycling
B GC DZ
B GC LZ
B GC prePB
B IFN
B mem
B naive
B plasma
B preGC
S
Cell Type Trend − T cells
NKT
T CD4+
T CD4+ naive
T CD4+ TfH
T CD4+ TfH GC
T CD8+ CD161+
T CD8+ cytotoxic
T CD8+ naive
T TfR
T TIM3+
T Treg
T
Cell Type Trend − Others
DC CCR7+
DC cDC1
DC cDC2
DC pDC
Endo
FDC
ILC
Macrophages M1
Macrophages M2
Mast
Monocytes
NK
VSMC

#### Figure S10

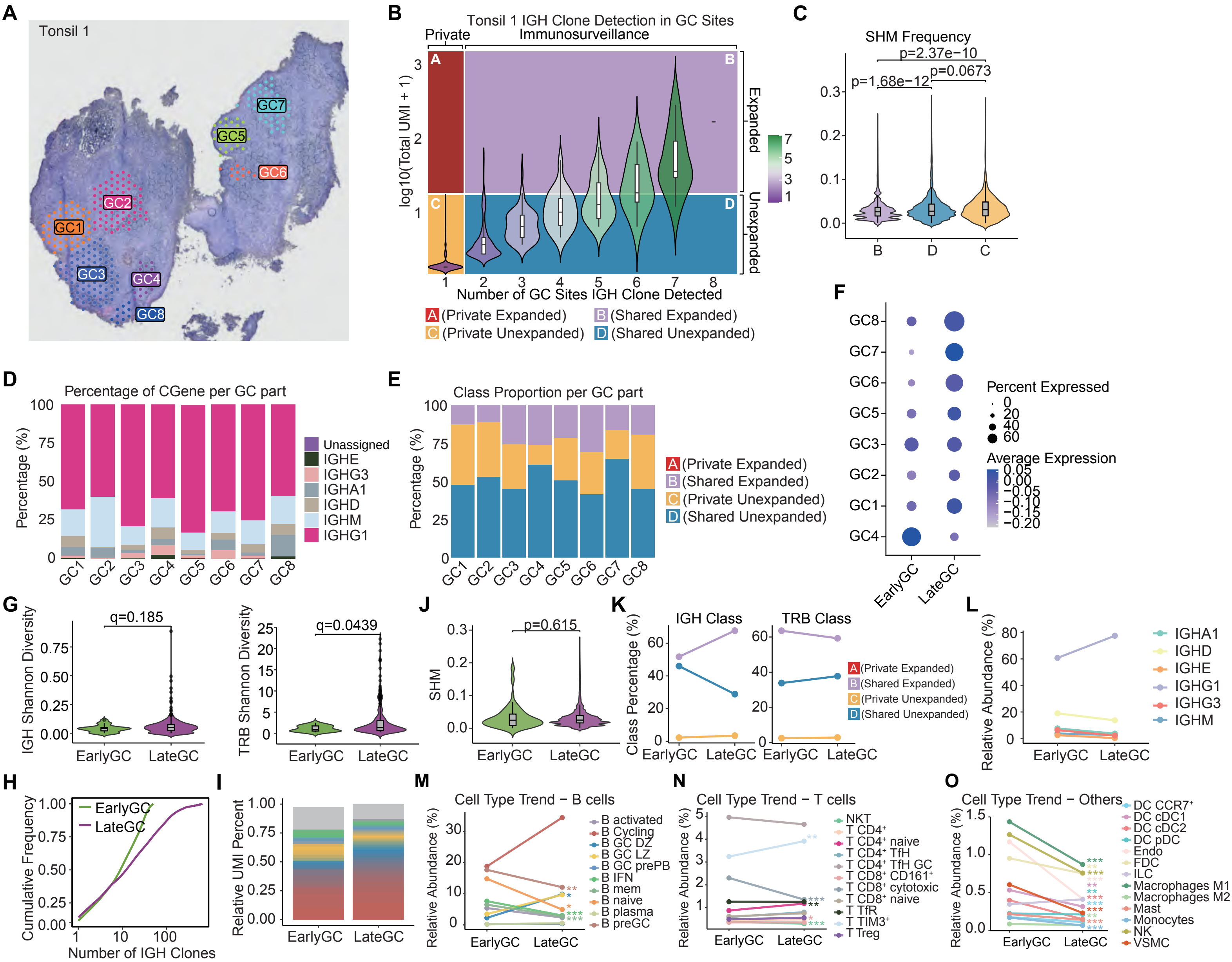

A
Tonsil 1
GC1
GC2
GC3
GC4
GC5
GC6
GC7
GC8
B
Tonsil 1 IGH Clone Detection in GC Sites
Private
Immunosurveillance
Expanded
Unexpanded
log10(Total UMI + 1)
Number of GC Sites IGH Clone Detected
A (Private Expanded)
B (Shared Expanded)
C (Private Unexpanded)
D (Shared Unexpanded)
C
SHM Frequency
p=2.37e−10
p=1.68e−12
p=0.0673
D
Percentage of CGene per GC part
Percentage (%)
Unassigned
IGHE
IGHG3
IGHA1
IGHD
IGHM
IGHG1
E
Class Proportion per GC part
Percentage (%)
A (Private Expanded)
B (Shared Expanded)
C (Private Unexpanded)
D (Shared Unexpanded)
F
Percent Expressed
Average Expression
EarlyGC
LateGC
G
IGH Shannon Diversity
q=0.185
TRB Shannon Diversity
q=0.0439
J
SHM
p=0.615
K
Class Percentage (%)
IGH Class
TRB Class
L
Relative Abundance (%)
IGHA1
IGHD
IGHE
IGHG1
IGHG3
IGHM
H
Cumulative Frequency
Number of IGH Clones
EarlyGC
LateGC
I
Relative UMI Percent
M
Cell Type Trend − B cells
Relative Abundance (%)
B activated
B Cycling
B GC DZ
B GC LZ
B GC prePB
B IFN
B mem
B naive
B plasma
B preGC
N
Cell Type Trend − T cells
NKT
T CD4⁺
T CD4⁺ naive
T CD4⁺ TfH
T CD4⁺ TfH GC
T CD8⁺ CD161⁺
T CD8⁺ cytotoxic
T CD8⁺ naive
T TfR
T TIM3⁺
T Treg
O
Cell Type Trend − Others
DC CCR7⁺
DC cDC1
DC cDC2
DC pDC
Endo
FDC
ILC
Macrophages M1
Macrophages M2
Mast
Monocytes
NK
VSMC

**Figure S11**

**A**

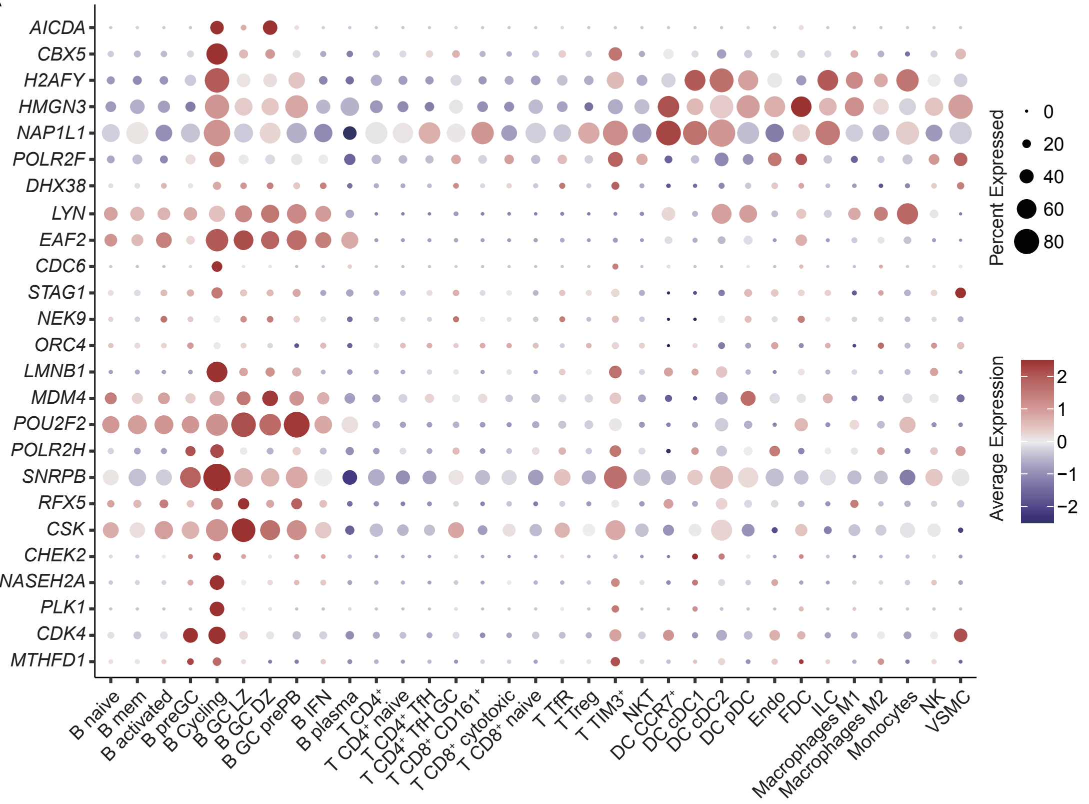

AICDA
CBX5
H2AFY
HMGN3
NAP1L1
POLR2F
DHX38
LYN
EAF2
CDC6
STAG1
NEK9
ORC4
LMNB1
MDM4
POU2F2
POLR2H
SNRPB
RFX5
CSK
CHEK2
RNASEH2A
PLK1
CDK4
MTHFD1
B naive
B mem
B activated
B preGC
B Cycling
B GC LZ
B GC DZ
B GC prePB
B IFN
B plasma
T CD4+
T CD4+ naive
T CD4+ TfH
T CD4+ TfH GC
T CD8+ CD161+
T CD8+ cytotoxic
T CD8+ naive
T TfR
T Treg
T TIM3+
NKT
DC CCR7+
DC cDC1
DC cDC2
DC pDC
Endo
FDC
ILC
Macrophages M1
Macrophages M2
Monocytes
NK
VSMC
Percent Expressed
0
20
40
60
80
Average Expression
2
1
0
−1
−2

#### Figure S12

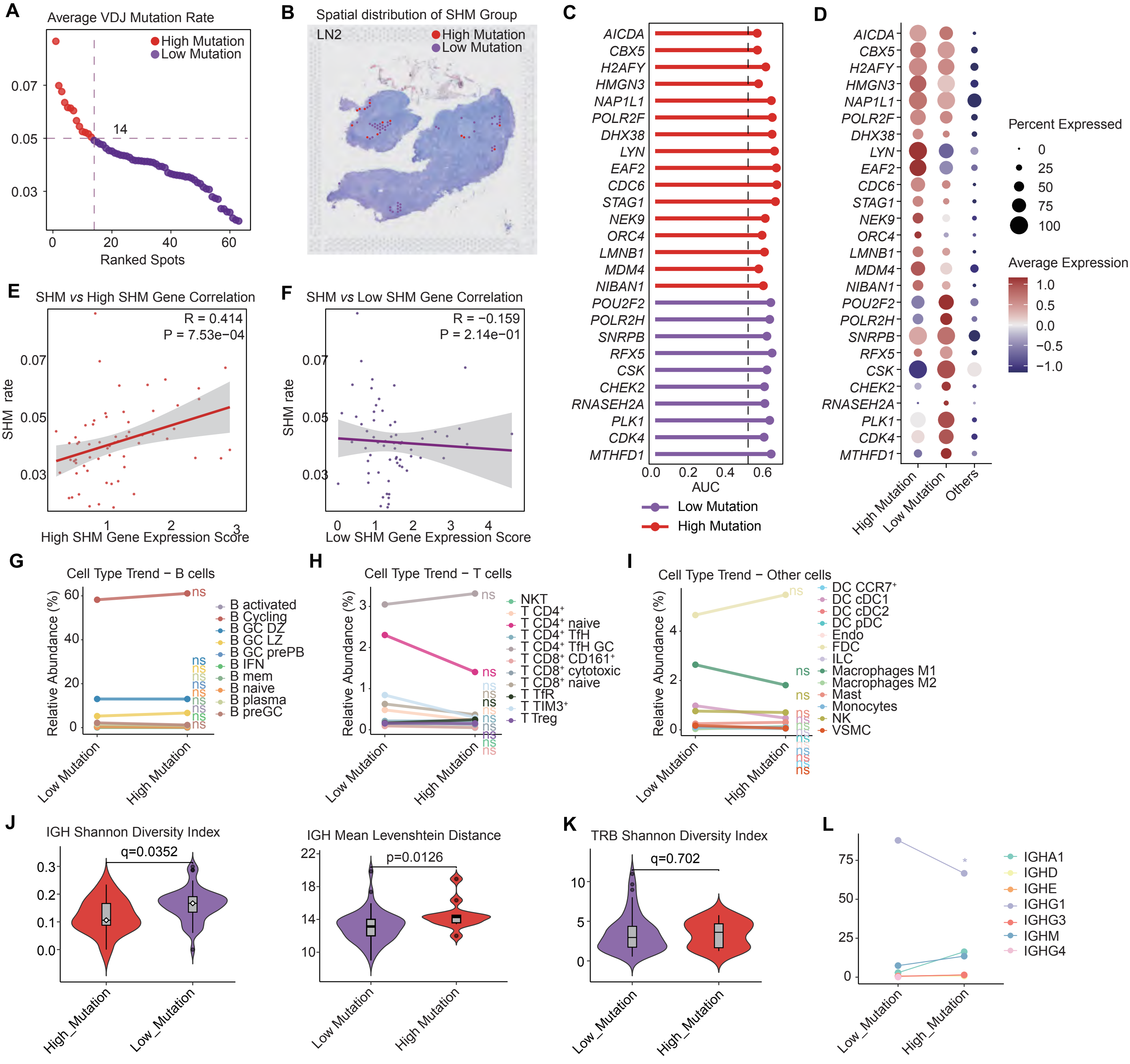

A
Average VDJ Mutation Rate
High Mutation
Low Mutation
14
Ranked Spots
B
Spatial distribution of SHM Group
LN2
High Mutation
Low Mutation
C
AUC
Low Mutation
High Mutation
AICDA
CBX5
H2AFY
HMGN3
NAP1L1
POLR2F
DHX38
LYN
EAF2
CDC6
STAG1
NEK9
ORC4
LMNB1
MDM4
NIBAN1
POU2F2
POLR2H
SNRPB
RFX5
CSK
CHEK2
RNASEH2A
PLK1
CDK4
MTHFD1
D
Percent Expressed
Average Expression
High Mutation
Low Mutation
Others
E
SHM vs High SHM Gene Correlation
R = 0.414
P = 7.53e−04
SHM rate
High SHM Gene Expression Score
F
SHM vs Low SHM Gene Correlation
R = −0.159
P = 2.14e−01
SHM rate
Low SHM Gene Expression Score
G
Cell Type Trend − B cells
Relative Abundance (%)
B activated
B Cycling
B GC DZ
B GC LZ
B GC prePB
B IFN
B mem
B naive
B plasma
B preGC
ns
Low Mutation
High Mutation
H
Cell Type Trend − T cells
NKT
T CD4+
T CD4+ naive
T CD4+ TfH
T CD4+ TfH GC
T CD8+ CD161+
T CD8+ cytotoxic
T CD8+ naive
T TfR
T TIM3+
T Treg
I
Cell Type Trend − Other cells
DC CCR7+
DC cDC1
DC cDC2
DC pDC
Endo
FDC
ILC
Macrophages M1
Macrophages M2
Mast
Monocytes
NK
VSMC
J
IGH Shannon Diversity Index
q=0.0352
High_Mutation
Low_Mutation
IGH Mean Levenshtein Distance
p=0.0126
Low Mutation
High Mutation
K
TRB Shannon Diversity Index
q=0.702
Low_Mutation
High_Mutation
L
IGHA1
IGHD
IGHE
IGHG1
IGHG3
IGHM
IGHG4
Low_Mutation
High_Mutation

### Figure S13

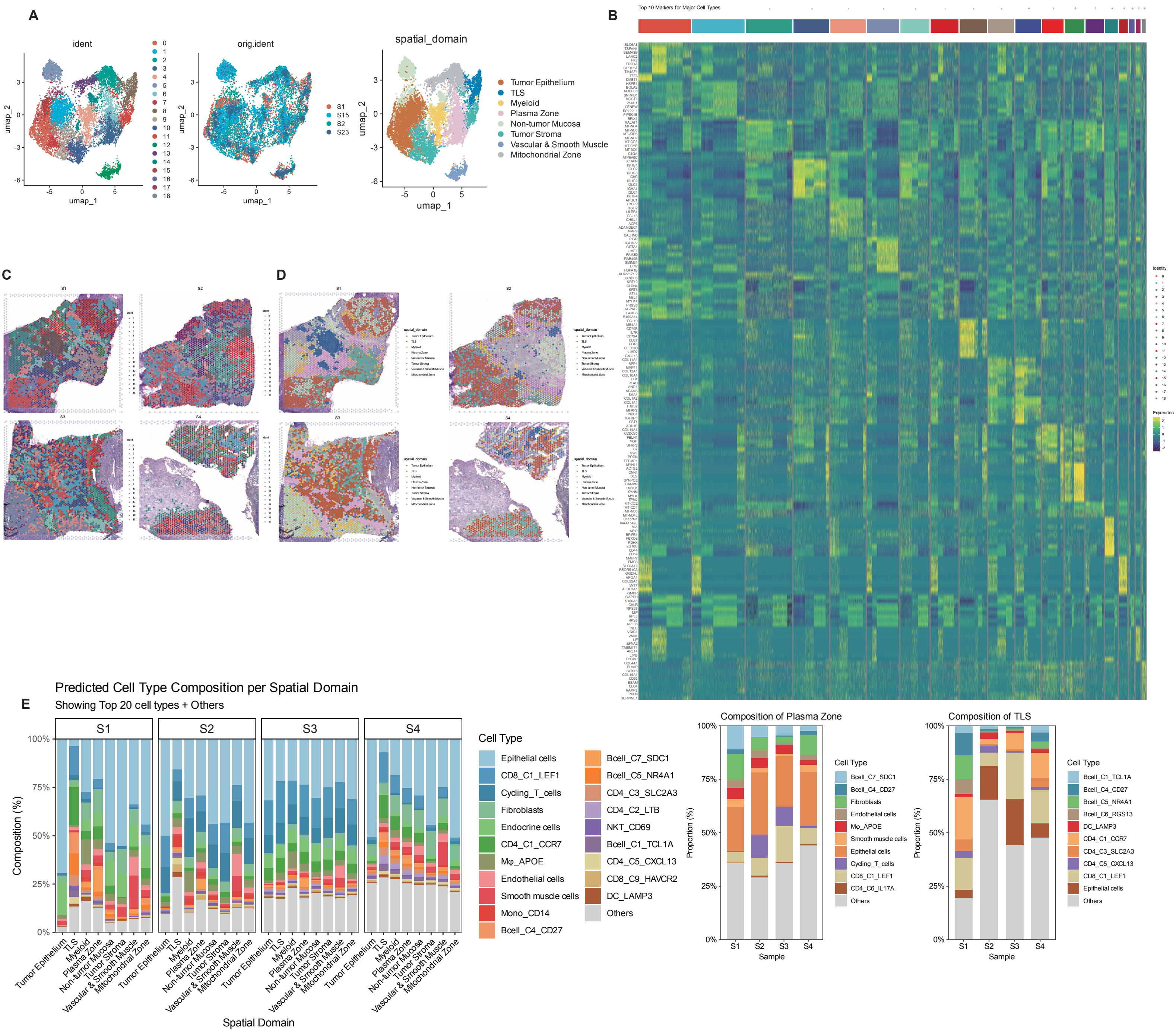
A
ident
orig.ident
spatial_domain
umap_1
umap_2
S1
S15
S2
S23
Tumor Epithelium
TLS
Myeloid
Plasma Zone
Non-tumor Mucosa
Tumor Stroma
Vascular & Smooth Muscle
Mitochondrial Zone
B
Top 10 Markers for Major Cell Types
C
D
Predicted Cell Type Composition per Spatial Domain
E
Showing Top 20 cell types + Others
S1
S2
S3
S4
Composition (%)
100%
75%
50%
25%
0%
Spatial Domain
Cell Type
Epithelial cells
CD8_C1_LEF1
Cycling_T_cells
Fibroblasts
Endocrine cells
CD4_C1_CCR7
Mφ_APOE
Endothelial cells
Smooth muscle cells
Mono_CD14
Bcell_C4_CD27
Bcell_C7_SDC1
Bcell_C5_NR4A1
CD4_C3_SLC2A3
CD4_C2_LTB
NKT_CD69
Bcell_C1_TCL1A
CD4_C5_CXCL13
CD8_C9_HAVCR2
DC_LAMP3
Others
Composition of Plasma Zone
Proportion (%)
Sample
Cell Type
Bcell_C7_SDC1
Bcell_C4_CD27
Fibroblasts
Endothelial cells
Mφ_APOE
Smooth muscle cells
Epithelial cells
Cycling_T_cells
CD8_C1_LEF1
CD4_C6_IL17A
Others
Composition of TLS
Proportion (%)
Sample
Cell Type
Bcell_C1_TCL1A
Bcell_C4_CD27
Bcell_C5_NR4A1
Bcell_C6_RGS13
DC_LAMP3
CD4_C1_CCR7
CD4_C3_SLC2A3
CD4_C5_CXCL13
CD8_C1_LEF1
Epithelial cells
Others

### Figure S14

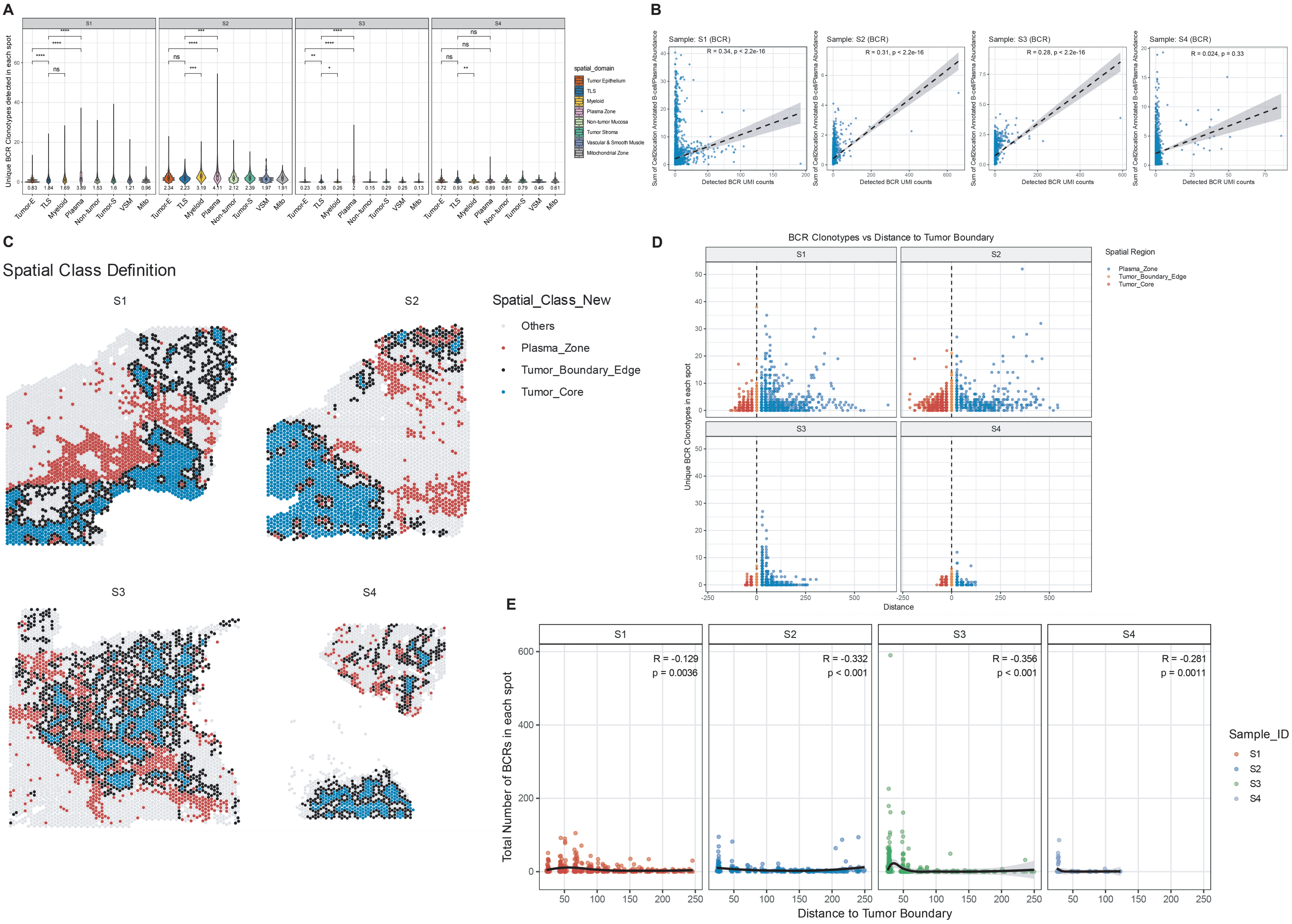

A
S1
S2
S3
S4
Unique BCR Clonotypes detected in each spot
spatial_domain
Tumor Epithelium
TLS
Myeloid
Plasma Zone
Non-tumor Mucosa
Tumor Stroma
Vascular & Smooth Muscle
Mitochondrial Zone
Tumor-E
TLS
Myeloid
Plasma
Non-tumor
Tumor-S
VSM
Mito
B
Sample: S1 (BCR)
R = 0.34, p < 2.2e-16
Sample: S2 (BCR)
R = 0.31, p < 2.2e-16
Sample: S3 (BCR)
R = 0.28, p < 2.2e-16
Sample: S4 (BCR)
R = 0.024, p = 0.33
Sum of Cell2location Annotated B-cell/Plasma Abundance
Detected BCR UMI counts
C
Spatial Class Definition
S1
S2
S3
S4
Spatial_Class_New
Others
Plasma_Zone
Tumor_Boundary_Edge
Tumor_Core
D
BCR Clonotypes vs Distance to Tumor Boundary
S1
S2
S3
S4
Spatial Region
Plasma_Zone
Tumor_Boundary_Edge
Tumor_Core
Unique BCR Clonotypes in each spot
Distance
E
S1
S2
S3
S4
R = -0.129
p = 0.0036
R = -0.332
p < 0.001
R = -0.356
p < 0.001
R = -0.281
p = 0.0011
Total Number of BCRs in each spot
Distance to Tumor Boundary
Sample_ID
S1
S2
S3
S4

### Figure S15

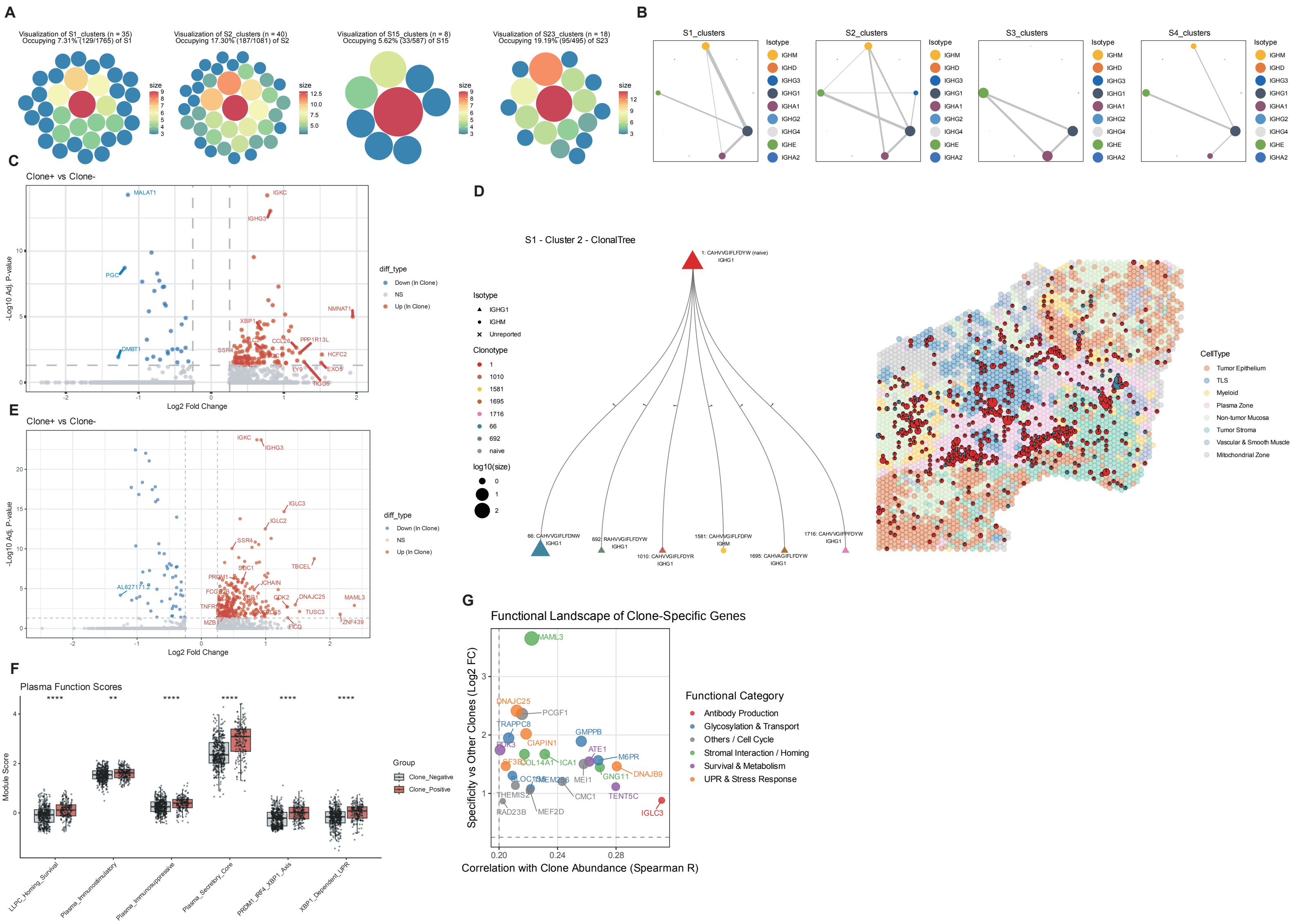
A
Visualization of S1_clusters (n = 35) Occupying 7.31% (129/1765) of S1
Visualization of S2_clusters (n = 40) Occupying 17.30% (187/1081) of S2
Visualization of S15_clusters (n = 8) Occupying 5.62% (33/587) of S15
Visualization of S23_clusters (n = 18) Occupying 19.19% (95/495) of S23
B
S1_clusters
S2_clusters
S3_clusters
S4_clusters
Isotype
IGHM
IGHD
IGHG3
IGHG1
IGHA1
IGHG2
IGHG4
IGHE
IGHA2
C
Clone+ vs Clone-
Log2 Fold Change
-Log10 Adj. P-value
diff_type
Down (In Clone)
NS
Up (In Clone)
D
S1 - Cluster 2 - ClonalTree
Isotype
Unreported
Clonotype
log10(size)
CellType
Tumor Epithelium
TLS
Myeloid
Plasma Zone
Non-tumor Mucosa
Tumor Stroma
Vascular & Smooth Muscle
Mitochondrial Zone
E
Clone+ vs Clone-
F
Plasma Function Scores
Module Score
Group
Clone_Negative
Clone_Positive
LLPC_Homing_Survival
Plasma_Immunostimulatory
Plasma_Immunosuppressive
Plasma_Secretory_Core
PRDM1_IRF4_XBP1_Axis
XBP1_Dependent_UPR
G
Functional Landscape of Clone-Specific Genes
Specificity vs Other Clones (Log2 FC)
Correlation with Clone Abundance (Spearman R)
Functional Category
Antibody Production
Glycosylation & Transport
Others / Cell Cycle
Stromal Interaction / Homing
Survival & Metabolism
UPR & Stress Response

### Figure S16

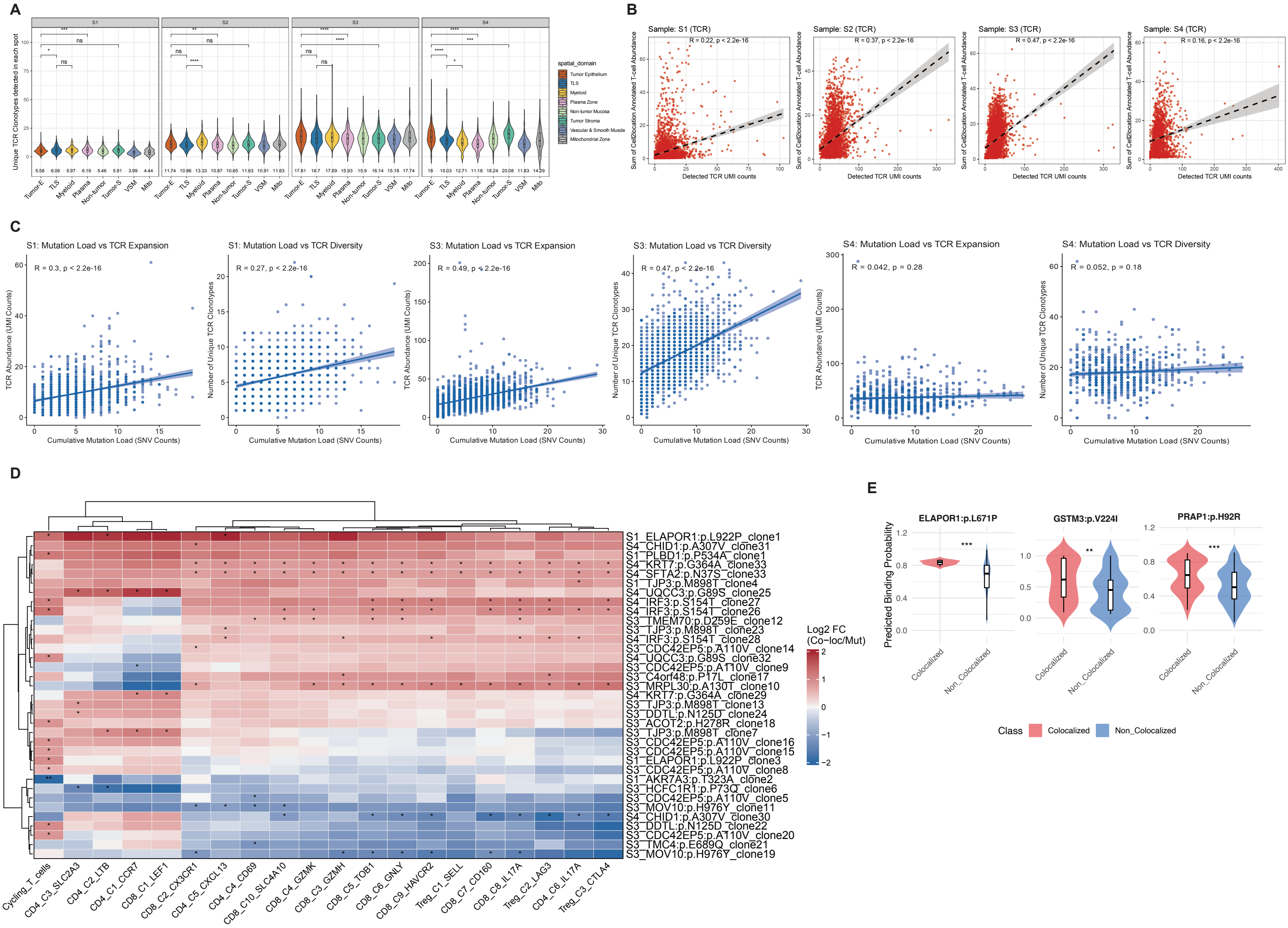
A
B
C
D
E
Sample: S1 (TCR)
R = 0.22, p < 2.2e-16
Sample: S2 (TCR)
R = 0.37, p < 2.2e-16
Sample: S3 (TCR)
R = 0.47, p < 2.2e-16
Sample: S4 (TCR)
R = 0.16, p < 2.2e-16
Detected TCR UMI counts
S1: Mutation Load vs TCR Expansion
R = 0.3, p < 2.2e-16
S1: Mutation Load vs TCR Diversity
R = 0.27, p < 2.2e-16
S3: Mutation Load vs TCR Expansion
R = 0.49, p < 2.2e-16
S3: Mutation Load vs TCR Diversity
R = 0.47, p < 2.2e-16
S4: Mutation Load vs TCR Expansion
R = 0.042, p = 0.28
S4: Mutation Load vs TCR Diversity
R = 0.052, p = 0.18
TCR Abundance (UMI Counts)
Number of Unique TCR Clonotypes
Cumulative Mutation Load (SNV Counts)
Log2 FC
(Co−loc/Mut)
ELAPOR1:p.L671P
GSTM3:p.V224I
PRAP1:p.H92R
Predicted Binding Probability
Colocalized
Non_Colocalized
Class
